# Impact of intercalators on the properties of DNA analyzed by molecular dynamics simulations

**DOI:** 10.64898/2026.04.02.716032

**Authors:** Hisashi Ishida, Hidetoshi Kono

## Abstract

Intercalation of small molecules between DNA base pairs affects DNA conformation, disrupting essential cellular processes including replication, transcription, and repair. We investigated conformational changes in 18-mer DNA upon intercalation of doxorubicin, SYBR Gold and YOYO-1 using extensive MD simulations. Two main patterns for the intercalation were identified: RISE-type intercalation occurs between adjacent base pairs and extends the DNA helix with decreased twist angles, while OPEN-type intercalation proceeds through base-pair opening without significant DNA extension. Kinetic analysis revealed that association rates for intercalation followed the order: first YO-moiety (mono-intercalation) > SYBR Gold > doxorubicin > YOYO-1 (bis-intercalation). Free energy landscape showed that forces at DNA termini reached up to 117 pN during stretching. Notably, base pairs adjacent to intercalators were protected from strand separation, accompanied by additional helical unwinding. MM-PBSA/GBSA analysis revealed that the driving force for intercalation is the stacking energy, and the binding affinity was highest for minor groove binding. Persistence length decreased with single molecule binding but recovered with two molecules due to their electrostatic repulsion. Mechanical properties of intercalated DNA showed position-dependence, demonstrating that multiple intercalation modes coexist in solution. The heterogeneous nature of intercalation explains why experimental measurements reflect ensemble averages rather than single binding configurations.

## Introduction

DNA intercalators are small organic molecules that are non-covalently inserted into the hydrophobic space between adjacent DNA base pairs of double-stranded DNA (dsDNA). The planar aromatic parts of the intercalators stabilize the dsDNA helix by new stacking interactions between them and adjacent base pairs [1]. DNA intercalation is considered to affect the conformation of the dsDNA, resulting in the disruption of essential cellular processes including replication, transcription and repair [2] [3].

Intercalators are used for many purposes such as DNA staining probes such as SYBR Gold (referred to as SYBR) and YOYO-1 (the oxazole yellow (YO) dimer, referred to as YOYO), and drugs such as doxorubicin (also called Adriamycin, referred to as DOXO). SYBR is a mono-intercalating cyanine dye. YOYO is a bis-intercalating dye that consists of two identical YO intercalative moieties connected by a linker and intercalates at two positions. DOXO, a mono-intercalator, is one of the anthracycline antitumor antibiotics which are among the most effective drugs for cancer chemotherapy [2] [3].

Intercalation can occur in two different ways: intercalation that occurs between adjacent base pairs and extends the DNA helix (we refer to this intercalation as RISE-type) and intercalation which flips out base pairs (we refer to this intercalation as OPEN-type). This is also known as intercalation through base-pair eversion or base-pair flipping out in molecular dynamics (MD) simulations [4] [5], or insertion in an experiment [6].

Many single molecule measurements observed the DNA extension upon intercalation. Optical tweezer experiments and atomic force microscopy showed that intercalation of DOXO [7] [8], SYBR [9] [10], and YOYO [9] [11] [12] extends the DNA and unwinds the double helix by 11º [7], 19.1º [10], and 24º [12] per molecule, respectively. However, it is difficult to analyze the precise location of the intercalation in the base pairs of the DNA at the atomic level by experiments.

Many computational calculations of the DNA with intercalators such as Ethidium bromide [13] [14], Copper complexes [4], Daunomycin [14] [15] [16] [17], and DOXO [5] [18] [19] were performed to analyze the conformational changes in the adjacent base pairs at the intercalation site at the atomic level. However, comprehensive research comparing different intercalation mechanisms and their structural effects on DNA remains limited.

To understand how the intercalation impacts DNA conformation at the atomic level, we carried out all-atom MD simulations of an 18-mer DNA with intercalators (DOXO, SYBR and YOYO). An adaptively biased molecular dynamics (ABMD) method was employed to facilitate the intercalation by repeating the extension and contraction of the DNA. This generated a variety of intercalated DNA at different base pair positions and major/minor grooves. For representative conformations of the intercalated DNA, umbrella sampling simulations were carried out to obtain the free energy curves (FECs) which characterize the free energy landscape of the DNA intercalation for the extension of the DNA. The molecular mechanics Poisson-Boltzmann surface area (MM-PBSA) approach was carried out to analyze the binding free energies of the intercalator–DNA complexes.

In this article, we report (1) diverse interaction modes between intercalators (DOXO, SYBR, and YOYO) and DNA base pairs, (2) free energy landscapes for DNA upon intercalation, (3) mechanical properties of intercalated DNA such as stretch modulus and persistence length, and (4) energetic contributions to binding affinity through MM-PBSA analysis. These findings elucidate how intercalation mode determines the mechanical response of DNA, providing molecular-level insights into intercalator-DNA interactions.

## Methods

### The force field parameters for the intercalators

The force field parameters describing the SYBR, YOYO, and DOXO molecules (Figure 1) were defined using the Antechamber package [20] and the general Amber force field (GAFF) [21] for organic molecules. These molecules were optimized at the HF/6-31G+ level by the Gaussian 16 software [22]. SYBR, YOYO and DOXO were protonated so that the net charge for SYBR, YOYO and DOXO were set at +2, +4, and +1, respectively. Discrete partial atomic charges were assigned to each atomic center in the molecules using the RESP methodology [23] at the HF/6-31G* level.

**Figure 1.**
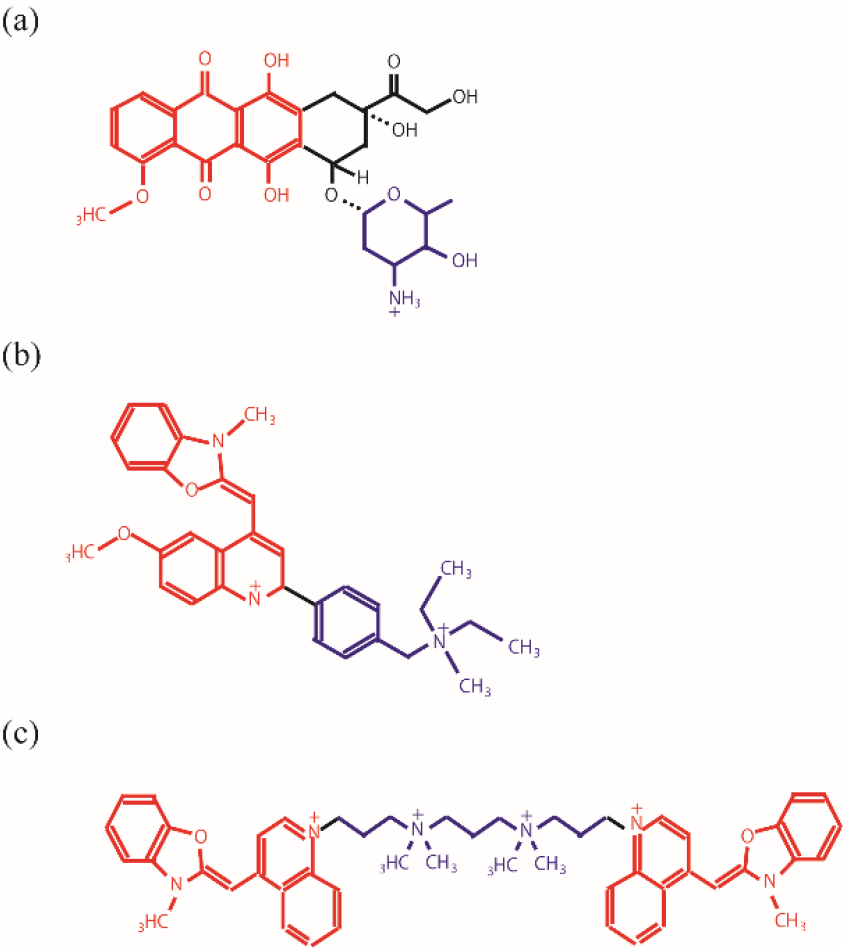
Structures of (a) DOXO, (b) SYBR, and (c) YOYO. The red group of atoms is defined as the intercalation part that intercalates into the double-stranded DNA helix. The blue group of atoms is defined as the bulky out-of-plane substitute to distinguish whether it is located at the major or minor groove.

### Construction of DNA

The initial structures of 18mer-DNA and 18mer-DNA with SYBR, DOXO, and YOYO were created using the Nucleic Acid Builder (NAB) module in the AMBER program [24]. The 18mer-DNA sequence for SYBR and DOXO was set to be d(GCATGAACGAACGAACGC)_2_ which contains a variety of base steps to analyze whether SYBR and DOXO have a specific preference for the sequence (Table I). The 18mer-DNA sequence for YOYO was selected so that CGCTAGCG is at the center as in the crystal structure of d(CGCTAGCG)_2_ with TOTO, YOYO-analog (PDB code:108d [25]).

**Table I.**
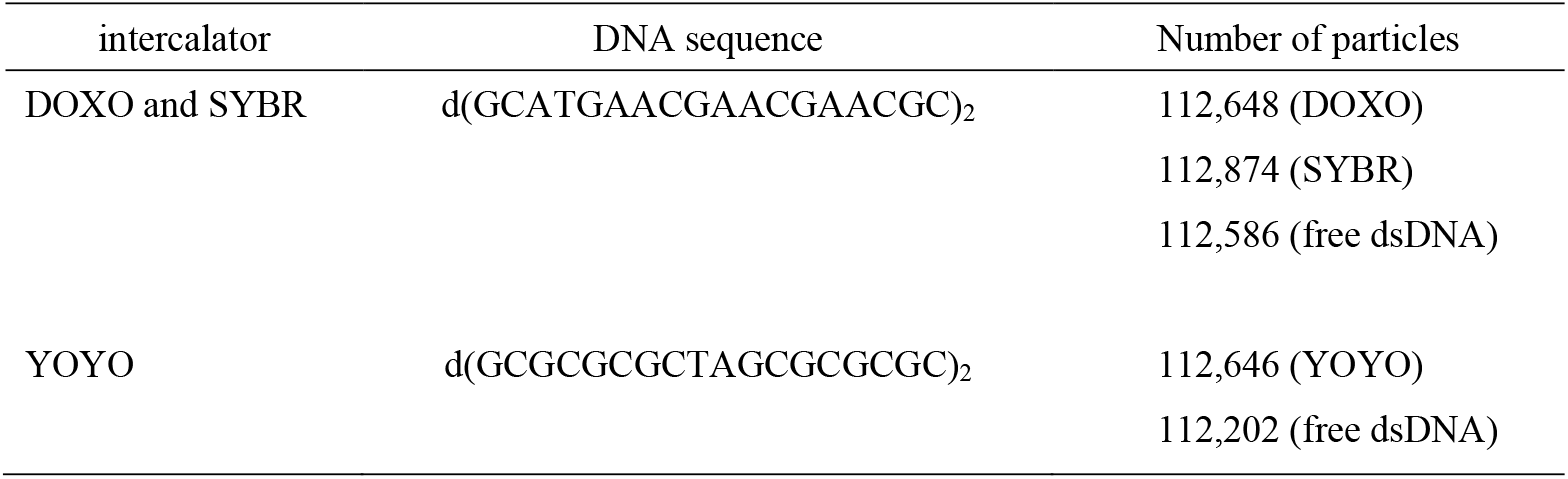
DNA sequence for intercalation.

### MD simulations of the intercalator – DNA complexes

After creating the 18mer-DNA structure, four intercalation molecules were placed around the center of the DNA, following the previous study by Galindo-Murillo and Cheatham [13]. It is known that many anthracycline antibiotics including DOXO tend to stack with each other in a high concentration, as is shown in experimental research [26]. In this study, distance restraints were imposed between intercalator molecules to alleviate strong stacking with each other. The potential for the distance restraints, *U*_*restr*_, was set at zero for *X* > *X2* = 20.0 Å for DOXO and SYBR, and for *X* > *X2* = 10.0 Å for YOYO, where *X* is the distance between the centers of mass of the intercalative parts of two DOXO and SYBR molecules for DOXO and SYBR, and the distance between the centers of mass of the intercalative parts of the YO-moieties of two YOYO molecules for YOYO. Therefore, no distance restraints are imposed on the intercalators for *X* > *X2*. For *X1* < *X* < *X2, U*_*restr*_ = *k*(*X* – *X2*)^2^ was used to produce the force of –2*k*(*X* – *X2*), where *X1* was set at 15.0 Å for SYBR and DOXO, and 5.0 Å for YOYO, respectively. The force constant of *k* was set at 0.1 kcal/mol/Å^2^. For *X* < *X1, U*_*restr*_ = *k*(*X* – *X1*) + *k*(*X1* – *X2*)^2^ was used to produce the constant force of 0.1 kcal/mol/Å. This restraint also worked to repel the four intercalator molecules from each other around the center of the DNA in the initial structure, so they instantly moved away from the DNA in almost all the cases. Therefore, the initial location of the four intercalator molecules would not affect the estimation of the value of *k*_*on*_, compared with the value in a system where the four molecules are randomly distributed.

Moreover, to avoid base pair fraying at the ends of the DNA, which would disturb stable sampling of the reaction coordinate in the following ABMD and umbrella simulations, the hydrogen bonds at the first and 18^th^ base pairs at the ends of the DNA were maintained through all the simulations by imposing a distance restraint with a harmonic force constant, which was set at 0.10 kcal/mol/Å^2^.

All simulations were carried out using an MD simulation program called GENESIS [27] [28] with the AMBER bsc1 [29] and *ff*99ions08 [30] force-fields for the DNAs and ions, respectively. The MD simulations of the DNA and the DNAs with the intercalators were carried out in a cubic box ∼105 Å × 105 Å × 105 Å at a constant pressure of 1 bar and a temperature of 300 K. Each system comprised ∼113,000 atoms with ∼37,000 TIP3P water molecules [31]. To neutralize the charges of the system, sodium ions were placed in the box, and then additional sodium and chloride ions were added at a concentration of 0.15 M NaCl. The summary of the procedure of the simulations is shown in Table SI.

The dielectric constant used was 1.0, and the van der Waals interactions were evaluated with a cut-off radius of 9 Å. The PME method [32] [33] was used for the electrostatic interactions for the direct space cutoff of 9 Å. The Langevin dynamics algorithm was utilized to control the temperature and pressure of the system. The temperature and pressure control coupling times were set at 2 ps^-1^. The SHAKE algorithm [34] [35] was used to constrain all the bond lengths involving hydrogen atoms. The leap-frog algorithm with a time step of 2 fs was used throughout the simulation to integrate the equations of motion.

### Adaptively Biased Molecular dynamics (ABMD) simulation to generate the DNA intercalation

It has been shown that the affinity constant of DNA intercalators increases exponentially with applied force [9]. To facilitate the intercalation and generate a variety of conformations of intercalated DNA, the adaptively biased MD (ABMD) simulations [36] of 24 replicas were independently carried out for an 18mer-DNA with intercalator molecules (Tables SI) for 700ns at a constant pressure of 1atm and a temperature of 300 K.

The reaction coordinate *d* was defined as the distance between the two centers of mass of the base-pairs at the ends of the DNA. The range of the reaction coordinate was set at 54.0 ≤ *d* ≤ 67.0 Å. High energy walls of 1,000 kcal/mol were set at *d* = 54.0 and 67.0 Å to restrict the extension and contraction of the DNA within the range of the reaction coordinates. The resolution of the reaction coordinate, Δ*d*, and the relaxation time for the free-energy profile, *τ*, were set at 1.0 Å and 1,000, 2500, 5000 ps and infinite (conventional MD without a biasing potential), respectively. The conformation of the DNA and intercalators was stored every 1 ps for analysis.

### ABMD simulation to estimate the biasing potential for the DNA intercalation

After the ABMD simulations, six, six, and three models of distinct DNAs with the intercalator molecule at different sequences in the major or minor groove were selected for SYBR, DOXO, and YOYO, respectively (Table SIV). The systems in which the DNA was severely deformed due to other intercalation molecules or excessive extension of the DNA were not selected.

From the conformational ensembles in the DNA intercalation of each model, a series of 24 conformations were selected. To estimate the biasing potential for the DNA intercalation for the selected conformations in the models, another ABMD simulation was carried out for 30 ns per model. The range of the reaction coordinate for the ABMD simulations was set at 48.0 ≤ *d* ≤ 71.0 Å for the free dsDNA, 51.0 ≤ *d* ≤ 74.0 Å for intercalated DNA with SYBR, DOXO, and one YOYO molecule, and 60.0 ≤ *d* ≤ 83.0 Å for intercalated DNA with two YOYO molecules. High energy potentials of 1,000 kcal/mol were set at the ends of the reaction coordinate to confine the system within the range of the reaction coordinate. Δ*d* and *τ* were set at 1.0 Å and 10,000 ps, respectively.

### Umbrella sampling simulations for refinement of the biasing potential and the production run

Using the conformations that were sampled in the ABMD simulation, 10 ns-umbrella sampling simulations were carried out from twice to four times to refine the biasing potential. To refine the biasing potential, the procedure of an evaluation of a temporal free energy curve (FEC) and using the FEC as a refined biasing potential was repeated several times (see reference [37] for the detailed procedure). The final umbrella sampling simulations were carried out for 500 ns to obtain the FEC for the extension of the intercalated DNA.

The umbrella potential for the *i*-th window used is

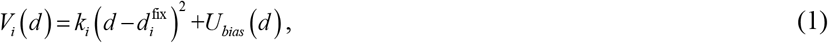

where 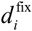 is a fixed distance from 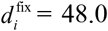 to 71.0 Å for the free dsDNA, 51.0 to 74.0 Å for the intercalated DNA with one DOXO and SYBR molecule, 54.0 to 77.0 Å for the intercalated DNA with one YOYO molecule, and 60.0 to 83.0 Å for the intercalated DNA with two YOYO molecules, at the interval of 1.0 Å. *k*_i_ is an arbitrary harmonic force constant, which was set at 0.20 kcal/mol/Å^2^. *U*_bias_ is the basing potential which was obtained in the ABMD simulation and replaced with the new basing potential obtained in the repeated 10 ns-umbrella sampling simulations. The WHAM algorithm [38] was used to obtain the FEC [37]. The last 450 ns of the trajectory was used for the analysis. The values of the free energy at *d* were estimated using sampling data (more than 100 data points per 0.1 Å out of 2,160,000 data points per 24 windows).

In our study, the data for the window at *d*_fix_ = 51.0 Å for DOXO Model 1, and the window at *d*_fix_ = 61.0 Å for SYBR Model 1 were excluded for the analysis because of the state change from RISE-type to OPEN-type during the umbrella sampling simulations.

### Conformational entropy of the DNA

The conformational entropies of the two ends of DNA_1_ or _2_ were calculated using the quasiharmonic approximation [39] as follows:

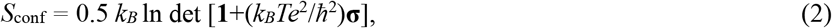

where *e* is *Euler*’s number, *ħ* is Planck’s constant divided by 2π. **σ** = < ***x x***^T^ > represents the mass-weighted covariance matrix, where ***x*** is the coordinates of the atoms in the base-pair step.

For the calculation of each covariance matrix for the base-pair step, ***x*** was best-fit in the reference coordinates. Each of the coordinates of the base-pair step in the initial structure in the *i*-th window of the umbrella sampling simulation was used as the reference coordinates for the best-fit at 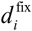. The trajectories in the *i*-th window of the umbrella sampling simulation where the desired position of 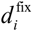 was set in Eq. (2) were used for the calculation of the conformational entropies against 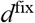. 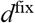, not *d*, was used to keep the number of sampled conformations the same for all 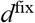. It should be noted that the conformational entropies calculated by the quasiharmonic approximation can be markedly overestimated because of the anharmonicity in protein dynamics [40].

### Definition of intercalation

*Rise* and *Opening* of the base-pair step parameter were analyzed using the DNA conformational analysis module in SCUBA [41] based on the definition of X3DNA [42].

1. If the value of *Rise* is greater than 5.0 Å at the DNA base-pair step of the *i*-th and *j*-th bases / (*i*+1)-th and (*j*–1)-th bases, then this conformation is a candidate for RISE-type (Figure 2(a)).
2. If the value of *Open* is greater than 90° or less than –90° at the base pair of the *i*-th and *j*-th bases, then this conformation is a candidate for OPEN-type (Figure 2(b)).

These conditions could spontaneously happen even in a single nucleosome. To confirm whether intercalation occurred at the base-pairs or not, the following quantities were monitored.

1. The center of mass of the intercalative part and the bulky out-of-plane substituent of the intercalator as COM(intercalative) and COM(substituent), respectively.
2. The centers of mass of the all the base pairs were calculated as the COM (*i,j*) by calculating the COM of *i-*th and *j*-th bases at the base pair.

If the distance between COM(intercalative) and COM(*i,j*), and the distance between COM(intercalative) and COM(*i*+1,*j*–1) are less than 4.0 Å, then the candidate is defined as the RISE-type at the DNA base step of *i*-th and *j*-th bases / (*i*+1)-th and (*j*–1)-th bases. If the distance between COM(intercalative) and COM(*i*–1,*j*+1), and the distance between COM(intercalative) and COM(*i*+1,*j*–1) are less than 4.0 Å, then the candidate is defined as the OPEN-type at the base pair of *i*-th and *j*-th bases.

**Figure 2.**
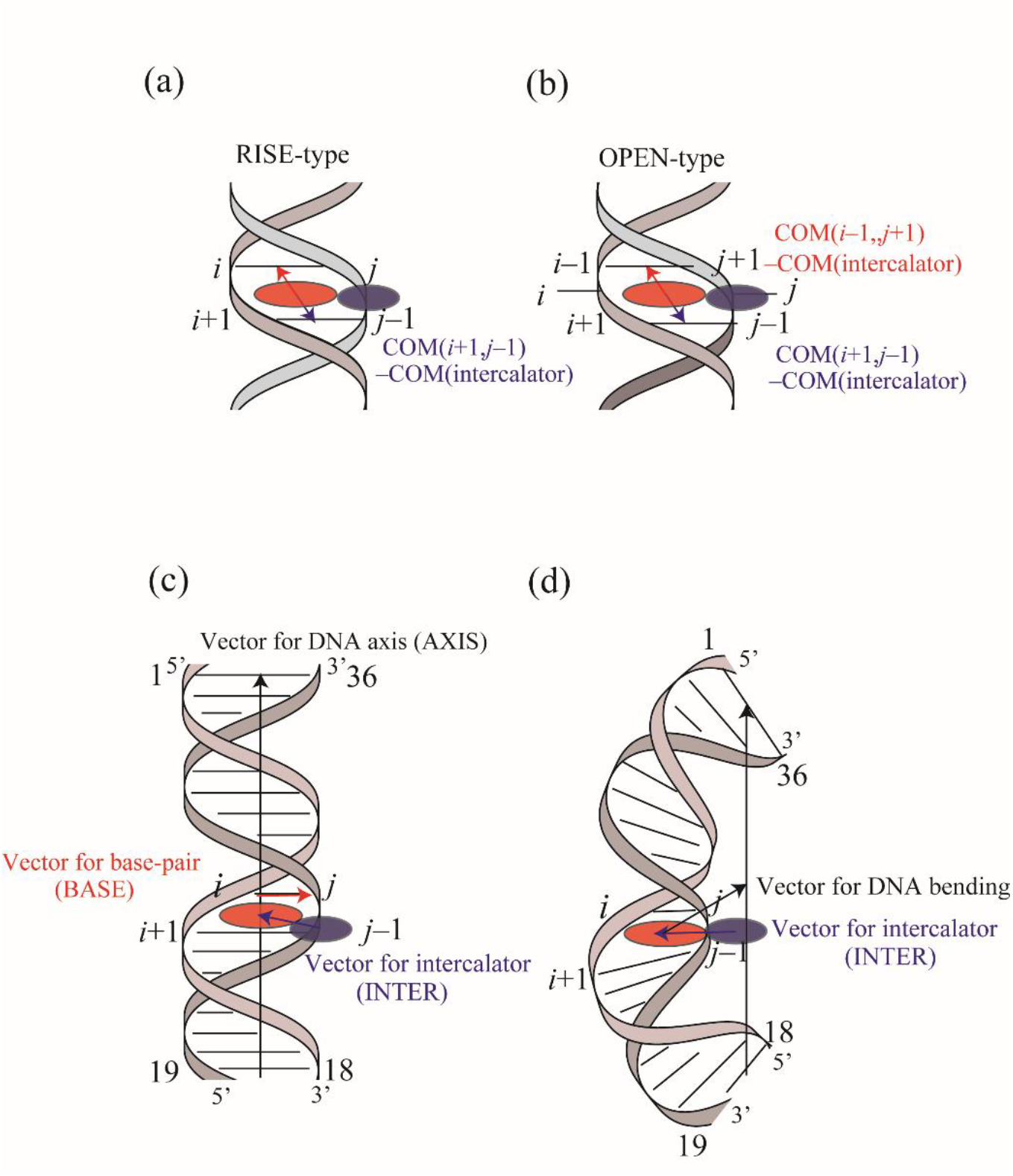
Schematic representations of (a) the RISE-type (extension of the base-pair stacking along the DNA axis) and (b) OPEN-type (opening of the base pairs towards the major or minor groove), (c) intercalator location at the major or minor groove, and (d) DNA bending. The planar aromatic part that intercalates in the double-stranded DNA helix is shown in red. The bulky out-of-plane substituent which is located at the major or minor groove is shown in blue (see also Figure 1).

### Definition of the location of the intercalators at the major and minor grooves

The location of the intercalators at the major or minor groove was defined by monitoring the location of the other part from the intercalative part at the major or minor groove, as follows (See also Figure 2(c));

1. COM(RISE-type) is the COM of the *i*-th and *j*-th bases and (*i*+1)-th and (*j*–1)-th bases. COM(OPEN-type) is the COM of the base pair of (*i*–1)-th and (*j*+1)-th bases / (*i*+1)-th and (*j*–1)-th bases.
2. INTER denotes the vector from COM(intercalative) to COM(RISE-tpye/OPEN-type).
3. BASE denotes the vector from the COM of *i*-th base to the COM of the *j*-th base.
4. AXIS denotes the vector from COM(1,36) and COM(18,19) on the edge of the DNA in the system of the 18mer-DNA.

If the value of (AXIS · (BASE × INTER)) is greater than 0, and less than 0, then it was defined that the intercalation occurred at the major groove and minor groove, respectively. In the case of the OPEN-type at the major and minor grooves, the base pairs flipped out towards these grooves.

### The direction of bent DNA towards the major and minor grooves

We monitored the direction of the DNA bent by comparing the direction of intercalation, INTER, and the vector from intercalator to the middle point of the DNA axis (see Figure 2(d)). If the dot product of these vectors > 0 then the DNA bend is in the direction of the intercalator; If the intercalator is at the major and minor grooves, the DNA bends towards the major and minor grooves, respectively. If the dot product of these vectors < 0 then the DNA bends in the opposite direction of the location of the intercalator; If intercalator is at the major and minor grooves, the DNA bends in the opposite direction.

### Kinetic analysis of k_on_

To analyze the kinetic rate of *k*_on_ for the intercalation process, we defined the time of occurrence of intercalation. First, we monitor whether the condition of intercalation is satisfied from time = 0 to 700 ns. When the condition is satisfied at time = *t*, then we counted the event for 100 ns from *t*, to *t* + 100 ns (or *t* to the final simulation time of 700 ns if *t* is > 600 ns). If the event was observed at the rate of more than 50 % in the range of 100 ns (or the same occurrence was observed after *t* = 600 ns), then we judged that intercalation stably occurred at time = *t*.

It should be noted that the *k*_*on*_ and Δ*G* are not directly related as Δ*G* = –ln *K* where *K* = *k*_*on*_ / *k*_*off*_. The value of *k*_*off*_ was not estimated in this study because its time scale is expected to be in the order of milli-seconds [9] [43] which is beyond the simulation time of a micro-second.

### Definition of fraction of intercalation

Figure S1 indicates that two intercalative molecules out of four in the system were substantially consumed to stack on the ends of the DNA, and the other two molecules can move relatively freely around the other base pairs from 2 to 17 and participate in the intercalation into the DNA. This phenomenon has been observed in other MD simulations [4] [5] [13].

Consequently, the total possible number of the intercalation sites was assumed to be two, and the number of the intercalation sites was normalized by two. Thus, the fraction of intercalation is defined as 1.0 when two DOXO or SYBR molecules are intercalated throughout all the 24 ABMD simulations, or when four YO-moieties of two YOYO molecules are intercalated throughout all the 24 ABMD simulations.

### Calculation of the stretch modulus

The extension of the DNA under a wide range of forces can be well described by an extensible worm-like chain (WLC) model [44]:

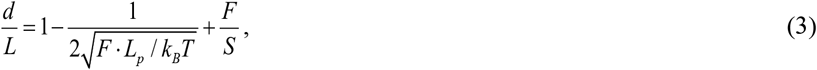

where *d* is the end-to-end distance of the DNA, *L* is the contour length of the DNA, *F* is the force applied to the DNA, *L*_*p*_ is the persistence length, *S* is the stretch modulus of the molecule. *k*_*B*_ is the Boltzmann constant, and *T* is the absolute temperature. In this equation, the first and second terms represent the entropic contribution while the third term represents the enthalpic contribution from the stretch modulus *S*. At low forces, only the first and second terms in Eq. (3) should contribute to the observed mechanics of the DNA. The force-extension curve is non-linear; a small increase in force leads to a significant change in the end-to-end distance as the DNA is pulled from a folded state.

At high forces, the second term of Eq. (3) becomes small, allowing the approximation *d* = *L*_0_ (1 + *F*/*S*), where *L*_0_ was set at the contour length at the free energy minimum state in our study. Here we assume that the fractional elongation for the contour length is the same as that for the end-to-end distance,

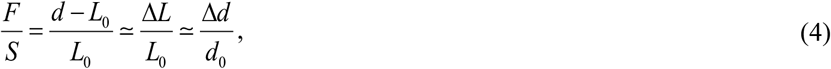

where Δ*L* = *L* – *L*_0_ is the extension from its original contour length, and Δ*d* = *d* – *d*_0_ is the extension from its free-energy minimum end-to-end distance. In this regime, the force-extension curve becomes relatively linear, and its slope corresponds to the stretch modulus *S*.

As stretching molecule experiment observes the DNA with a long chain (about kb), *L*_*p*_ is measured at a low force (< 10 pN) in the entropic regime and *S* is measured in the enthalpic regime. In our study, the length of the DNA (about 6 nm) is much shorter than the persistence length so we measure both *L*_*p*_ and *S* in the enthalpic regime.

### Calculation of the persistence length

The persistence length *L*_*p*_ was estimated to characterize the bending rigidity of the DNA. Based on the WLC model, the bending persistence length *l*_B_ of dsDNA are determined by fitting the bending angle distribution to a quadratic function of the bending angle [45] [46] as follows:

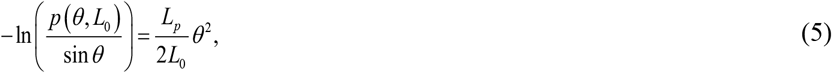

Here, *p*(*θ, L*_*0*_) is an angular probability distribution, where *θ* is the bending angle, and *L*_*0*_ is the average of the instantaneous contour length of 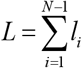, where *l*_*i*_ is the distance between the centers of successive base pairs in a step, and *N* is the total number of base pairs. *l*_*i*_ is calculated by 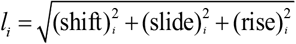 using X3DNA [42]. *N* =18 in the case of the RISE-type, and *N* = 17 in the case of OPEN-type as the flipped-out base pair was ignored. The bending angle was determined by the angle between a vector from the center of mass (COM) of the 2^nd^ base pair to the COM of the 1^st^ base pair and another vector from the COM of 17^th^ base pair to the COM of the 18^th^ base pair. To estimate the persistence length of the DNA, only the trajectory which is within the elastic range (Table II) was used to avoid the force dependency. In the calculations of bending angle probabilities *p*(*θ, L*_*0*_), a bin width of 1.0 degree was employed for bending angle *θ*.

### Calculation of the stretch modulus from the contour length

Stretch modulus, *S*, is obtained from the contour length distribution as follows [45] [46]:

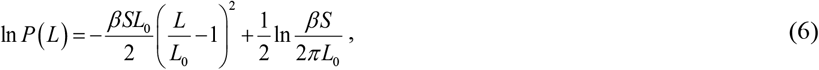

where *β* = 1/*k*_*B*_*T*. From the slope of the graph of ln *P*(*L*) versus (*L*/*L*_0_ – 1)^2^, the stretch modules can be measured. To estimate the stretch modulus, trajectories within the elastic region in *d* were used in our study.

### MM-PBSA/GBSA

MM-PBSA/GBSA [47] is described as follows:

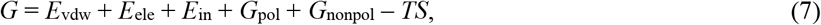

where *G* is the calculated average free energy, and *E*_vdw_, *E*_ele_ and *E*_in_ are the average molecular mechanical energy of the van der Waals interaction, the electrostatic interaction, and the internal energy such as the bond, angle and torsion energy of the solute, respectively. The MMPBSA.py tool was used to calculate PBSA/GBSA [48]. The mbond2 was selected as atomic radii [49]. The solute and solvent dielectric constants were set at 1 and 80, respectively. *G*_pol_ is the polar part of the solvation free energy. *G*_nonpol_ is the nonpolor contribution to the solvation free energy calculated using the solvent accessible surface area (SASA) of the solute. *G*_nonpol_ is obtained from the SASA as follows, *G*_nonpol_ = *γ*SASA + *b*, where *γ* is a surface tension of 0.00542 kcal/mol/Å^2^ and *b* is 0.92 kcal/mol for MM-PBSA [50], while *γ* is 0.0072 kcal/mol/Å^2^ [51] and *b* is 0.0 for MM-GBSA [51],. Salt concentration was set at 0.150 M. –*TS* is the entropic contribution, where *S* is the entropy of the solute and *T* is the temperature. MM-GBSA analysis was performed on the trajectories at an interval of 100 ps.

In spite of the uncertainty of the accuracy of the absolute binding free energy, it has often been suggested that MM-GBSA can qualitatively evaluate the relative binding free energy for protein-DNA complexes [52] [53] without taking account of the entropic contribution. Consequently, the entropy term was omitted in this study.

The biding free energy, Δ*G*_bind_ was calculated as:

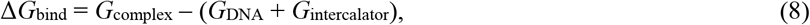

from the difference between the free energy of the DNA – intercalator complex, G_complex_, and the free energy of the DNA/intercalator, *G*_DNA/intercalator_, using the trajectory of the complex. The trajectories for *G*_DNA_ and *G*_intercalator_ were extracted from the trajectory for the complex. The value of Δ*G*_bind_ in the MM-PBSA/GBSA calculation is known to be much larger than that of the experimental Δ*G*_bind_ [52] [53]. However, it has been pointed out that it was quantitatively valid to discuss the relative stability of the two states from the difference in their free-energies [52] [53].

Further, to account for the deformation energy of the DNA (*G*_def_), *G*_def_ was calculated as the difference in energy between the free dsDNA and the intercalated DNA [17]. The final corrected Gibbs binding free energy (Δ*G*_corr_) is given by:

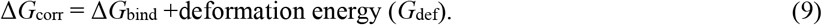

### Computational time

The simulation times for (1) the ABMD simulation to generate the DNA intercalation, (2) the ABMD simulation to estimate the biasing potential for the extension of the DNA, (3) the umbrella sampling simulations to refine the biasing potential, and (4) the umbrella sampling simulation for the production run are summarized in Table II. All the simulations, requiring a total time of 14.6 μs (3.1 μs for the ABMD simulations and 11.5 μs for the umbrella sampling simulations), were mainly performed on the Polaire supercomputer at Hokkaido University and the SGI8600 supercomputer at Japan Atomic Energy Agency.

## Results

### A variety of intercalated DNA generated by ABMD simulations

We considered three intercalators, DOXO, SYBR, and YOYO, and generated a variety of intercalated DNA conformations using the adaptively biased MD (ABMD) simulation for an 18mer-DNA with intercalator molecules for 700 ns at 1 atm pressure and 300 K (see Methods and Table SI), as well as a conventional MD simulation.

We classified the obtained conformations into four categories dependent on RISE or OPEN type at the major or minor grooves: RISE(major), RISE(minor), OPEN(major), and OPEN(minor) (Figure 3 (a) and (b)).

**Figure 3.**
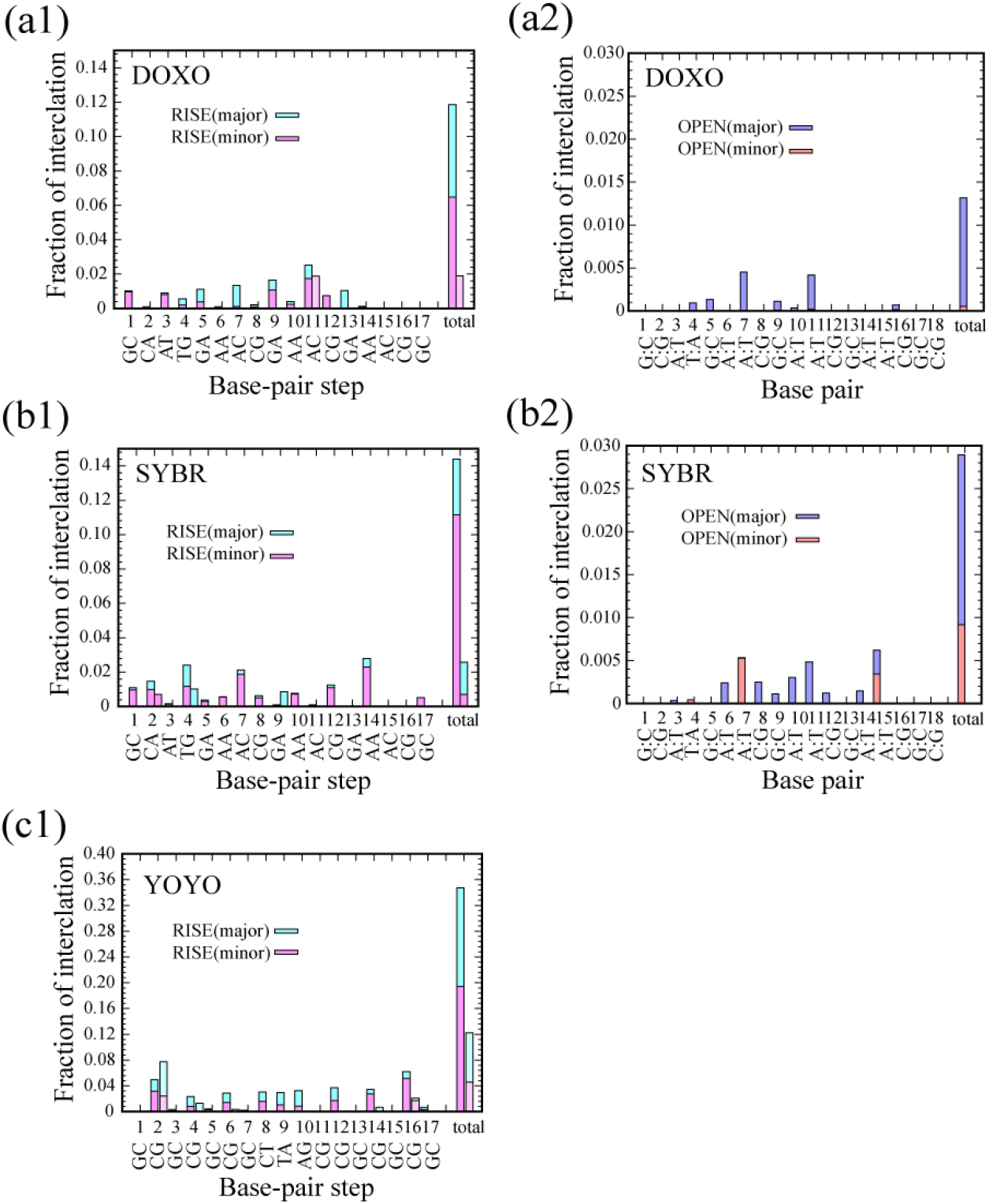
The fraction of intercalation for RISE-type and OPEN-type. The fraction of intercalation for RISE-type for (a1) DOXO, (b1) SYBR, and (c1) YOYO along the DNA base-pair step, and OPEN-type for (a2) DOXO and (b2) SYBR along the DNA base-pair, respectively. At each step, a bar for the average of tau = 1500 ps, 2500 ps, 5000 ps, and another bar for the conventional MD are shown from left to right. RISE-type in the major and minor grooves, and OPEN-type in the major and minor grooves are shown in blue, red, cyan, and magenta, respectively. The probability of OPEN-type at *i*-th and *j*-th base pair is shown at *i*-th DNA bp-step. The fraction of intercalation is normalized by 24, the number of replicas, and 2, the number of DOXO molecules which participate in intercalation (48 in total). This means that the fraction of intercalation is one if two molecules always intercalated in the DNA. The total fraction of intercalation from 1 to 17 is shown at 19 of the base-pair steps.

We defined the intercalation fraction such that 1.0 represents the state in which two of the four intercalator molecules are continuously involved in intercalation (see Method). For DOXO, ABMD yielded significantly higher intercalation fractions compared with conventional MD: RISE(major): 5.37×10^−2^ vs 1.89×10^−2^, RISE(minor): 6.48×10^−2^ vs 0.0, OPEN(major): 1.26×10^−2^ vs 0.0, and OPEN(minor): 5.36×10^−4^ vs 0.0. Notably, OPEN-type conformations were completely absent in conventional MD (Figures 3(a1), (a2), and Table SII). Intercalation was facilitated by the ABMD simulation. RISE-type was observed 1.5 times more frequently than expected based on sequence compositions at the AC, GA, and AT steps, while it was less often at the AA, CA(=TG), and CG steps. We also analyzed the base pair composition at RISE-type sites. In the 18-mer DNA, A:T and C:G base pairs comprised 8 (44.4%) and 10 (55.6%) pairs, respectively. The observed intercalation distributions in ABMD (47.8% A:T, 52.1% C:G) and conventional MD (50.1% A:T, 49.9% C:G) simulations were consistent with this composition, indicating no significant sequence preference for either base pair type.

In contrast to the RISE-type interaction, OPEN-type at the A:T and G:C base pairs was 76.0% and 24.0% in the ABMD simulation, respectively, indicating that the A:T base pair more easily facilitates OPEN-type (Figure 3(a2)). This is probably due to the weaker hydrogen bonds in A:T pairs compared with G:C pairs. The 18-mer DNA contains three 5’-GAAC-3’ segments (base pairs 5-8, 9-12, and 13-16). OPEN-type intercalation occurred at the 7th and 11th A:T base pairs (the second A in each GAAC motif) for DOXO, and at the 7th, 11th, and 15th positions for SYBR, but was absent at the 6th, 10th, and 14th A:T base pairs (the first A in each motif). This indicates that OPEN-type intercalation preferentially occurs at A:T base pairs in the middle of 5’-AAC-3’ to 5’-GAA-3’. Notably, these OPEN-type conformations were not observed in conventional MD simulations.

For SYBR, the fraction of RISE(major), RISE(minor), OPEN(major), and OPEN(minor) in the ABMD simulation was 3.23×10^−2^, 1.12×10^−1^, 1.98×10^−2^, and 9.17×10^−3^, compared with 1.88×10^−2^, 7.09×10^−3^, 0.0, and 0.0 in the conventional MD simulation (Figures 3(b1) and (b2), and Table SII), indicating that SYBR intercalation is also facilitated by the ABMD simulation. RISE-type intercalation occurred1.5 times more frequently than expected at the AA, CA(=TG) steps, while occurring less frequently at the GA and AT steps. The base pair composition at RISE-type intercalation sites showed 60.7% A:T and 39.3% C:G in ABMD simulation, and 50.1% A:T and 49.9% C:G in conventional MD simulation (Table SII). Compared with the sequence composition (44.4% A:T and 55.6% C:G), this indicates a moderate preference for A:T base pairs in SYBR intercalation. OPEN-type intercalation occurred predominantly at A:T base pairs (83.1%) compared with C:G base pairs (16.9%) in ABMD simulation, similar to DOXO, but was not observed in conventional MD simulation.

YOYO bis-intercalation occurred in two steps: first, one YO moiety intercalated between DNA base pairs; subsequently, the second YO moiety intercalated at a nearby site. For the YO-moieties of YOYO, the fraction of RISE(major), RISE(minor), OPEN(major), and OPEN(minor) in the ABMD simulation was 1.53×10^−1^, 1.94×10^−1^, 0.0, and 0.0, compared with 7.67×10^−2^, 4.58×10^−2^, 0.0, and 0.0 in the conventional MD simulation. Intercalation was also observed 1.5 times more frequently than expected at the AG(=CT) and CG steps, while it was less frequently at GC step (Figures 3(c1) and Table SII). The first intercalation of the YO moieties clearly showed a strong preference for intercalating to the CG steps but little preference for intercalating to the GC steps (Figure 3(c1)). The AG(=CT) steps are positioned in the center of d(CGCTAGCG)_2_, which we took from the complex structure of DNA and TOTO (YOYO analog of the thiazole orange (TO) dimer) (PDB code: 108d [25]). We observed bis-intercalation events where the two moieties intercalated at sites separated by one base pair: specifically, at C6G7/C8T9, A10G11/C12G13, and C4G5/C6G7 steps. Notably, bis-intercalation did not occur at the C8T9/A10G11 step in the center of the 5’-CGCTAGCG-3’ sequence, despite this being the known intercalation site for TOTO in crystal structures [21]. No OPEN-type intercalation was observed for YOYO even in the ABMD simulation.

In summary, the ABMD simulation revealed distinct base-pair step and sequence preferences for each intercalator: RISE-type occurred preferentially at AC, GA, and AT steps for DOXO, AA and CA steps for SYBR, and AG and CG steps for YOYO. OPEN-type intercalation was predominantly observed at A:T base-pairs for both DOXO and SYBR, but was absent for YOYO even in ABMD simulations.

### Intercalation kinetics of DNA–intercalator complexes

We examined the intercalation kinetics under the DNA extension and contraction force with ABMD. Figure 4 shows that *k*_on_ for *τ* = 1500 ps is larger than that for *τ* = 2500 and 5000 ps except for YOYO (first intercalation). This suggests that more frequent extension -contraction cycles (shorter *τ*) generally enhance intercalation rates. Notably, *k*_on_ values could not be reliably determined from conventional MD simulations due to insufficient intercalation events, except for YOYO mono-intercalation.

**Figure 4.**
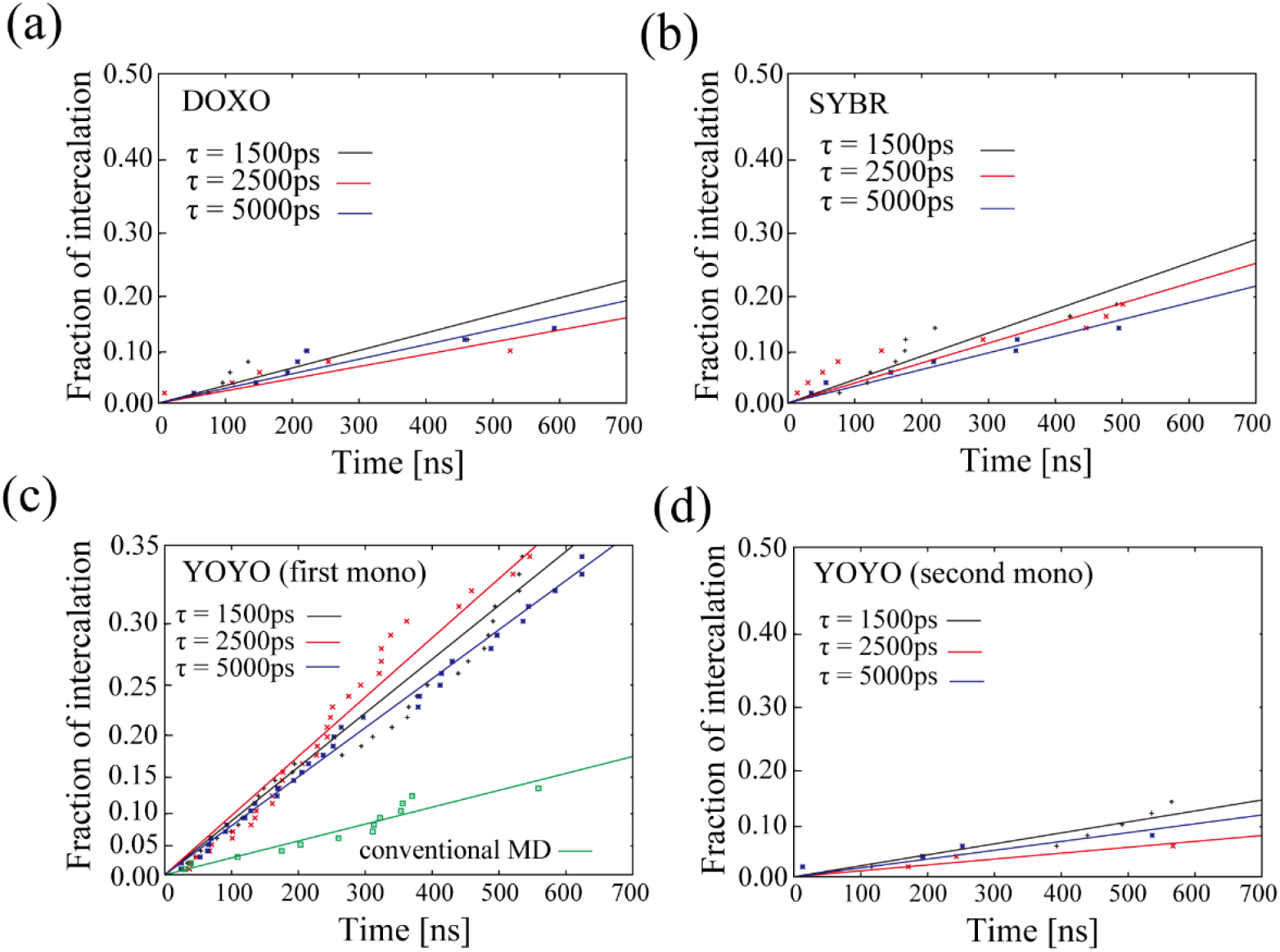
Kinetics of intercalated DNA for (a) DOXO, (b) SYBR, (c) YOYO (first mono-intercalation), and (d) YOYO (second mono-intercalation). The plot shows the fraction of intercalation, exp (-*k*_on_ *t*), with the kinetic rate of *k*_on_ and the time of intercalation. The time of intercalation is defined in Methods. *k*_on_ was estimated based on a simple two-state model for the reaction of an intercalator (I) with the DNA (D), i.e., [D] + [I] = [DI], where [I] has a first order exponential kinetics as [I]_*t*_ = [I]_0_ exp (−*k*_*on*_ · *t*). The data in the graphs represent the intercalation events observed in the 24 MD simulations of 700 ns for each time of τ. For DOXO, the values of *k*_on_ for *τ* = 1500, 2500, and 5000 ps were estimated to be 271, 394, and 325 ns, respectively. For SYBR, they were estimated to be 204, 238, and 285 ns^-1^, respectively. For the first mono-intercalation of a YO moiety of YOYO, the values of *k*_on_ for τ = 1500, 2500, 5000ps, and infinity were estimated to be 50.8, 46.2, 55.8, and 162 ns^-1^, respectively. For the second intercalation of the other YO moiety (bis-intercalation of YOYO), the values of *k*_on_ for τ = 1500, 2500, and 5000 ps were estimated to be 434, 807, and 539 ns^-1^, respectively.

The values of *k*_on_ were calculated by converting the slope [s^-1^] in Figure 4 to [M^-1^s^-1^] in Table SIII under the assumption that the concentration of intercalators in the MD systems is 2,970 mM based on the simulation box volume (see Methods). The kon values (×108 M^-1^s^-1^) for τ = 1500, 2500, and 5000 ps were: 1.24, 0.86 and 1.04 (DOXO), 1.65, 1.41 and 1.18 (SYBR), and 6.63, 7.29 and 6.03 (YOYO first or mono-intercalation). For comparison, conventional MD yielded 2.08×108 M-1s-1 for YOYO mono-intercalation. The kon value for the mono-intercalation is higher than that of kon for SYBR and DOXO. This is reasonable because YOYO has two YO moieties that can independently attempt intercalation, effectively doubling the binding opportunity. However, the second intercalation step (kon values of 7.76×107, 4.17×107 and 6.25×107 M^-1^s^-1^ for τ = 1500, 2500, and 5000 ps, respectively) was approximately one order of magnitude slower than mono-intercalation, likely due to conformational constraints imposed by the first bound moiety and the requirement for proper spatial positioning of the linker.

Overall, our results showed the values of kon were large in the order of first YO-moiety of YOYO (mono-intercalation) > SYBR > DOXO > second YO-moiety of YOYO (bis-intercalation).

### Free energy profiles of the intercalated DNA for the extension

We calculated free energy profiles for specific intercalated DNA models with intercalators at different base pair positions: 6 models each for DOXO (Figure 5A) and SYBR molecule (Figure 5B), 2 models for one YOYO, and two YOYO molecules (Figure 5C). These models were selected because they maintained intact base pairing throughout the repeated extension and contraction in ABMD simulations. Non-intercalated molecules were removed from subsequent simulations.

**Figure 5.**
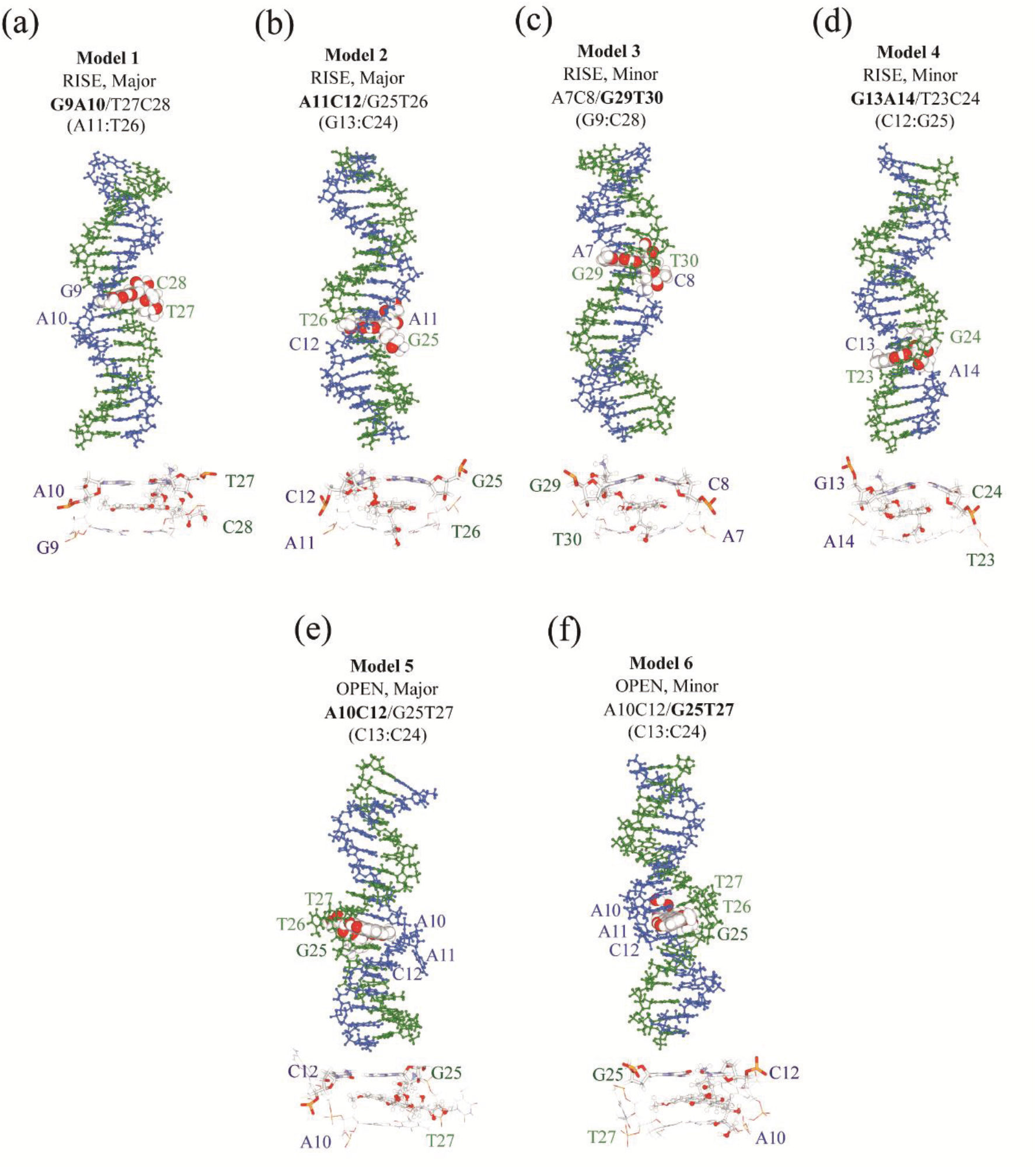
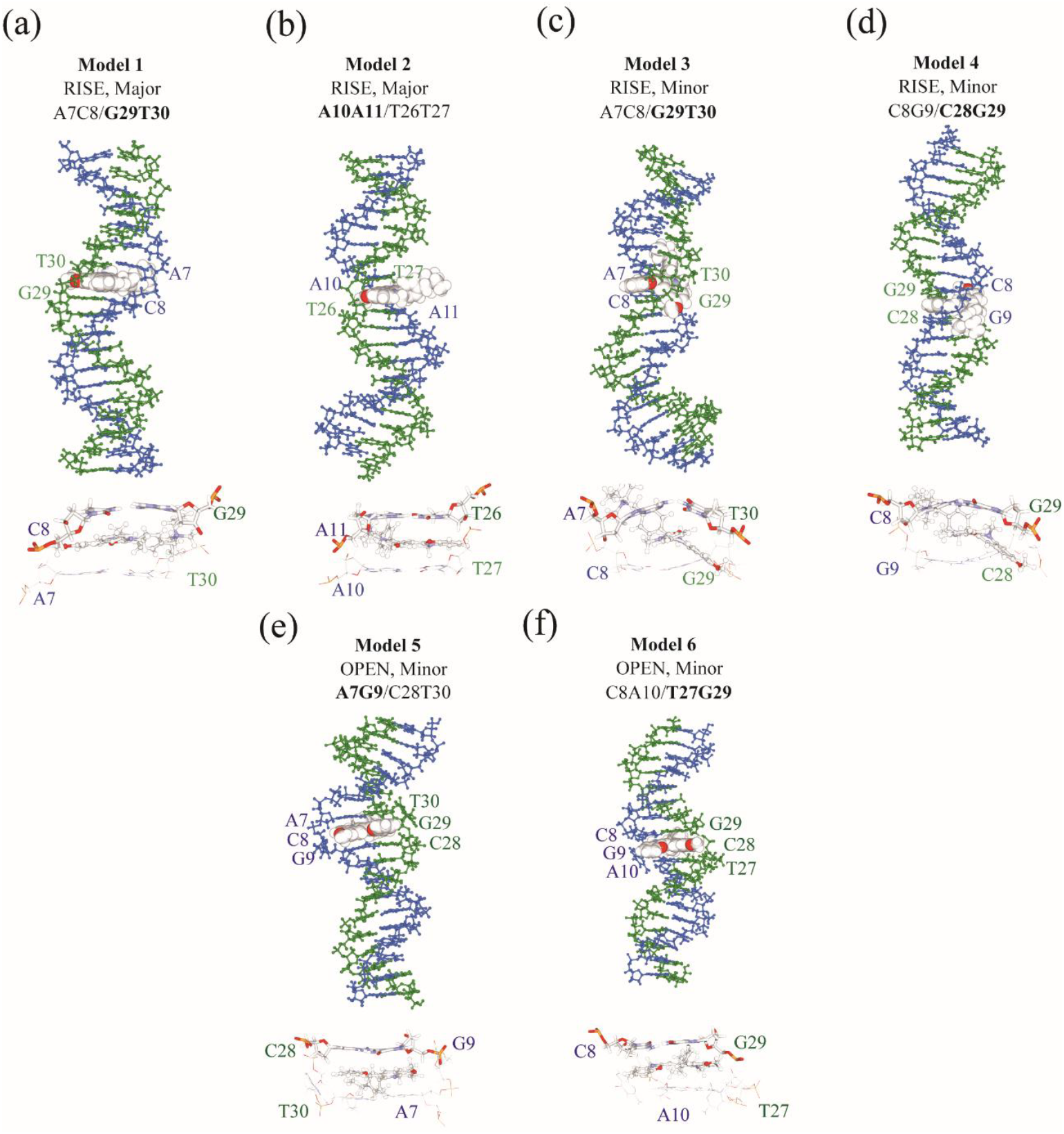
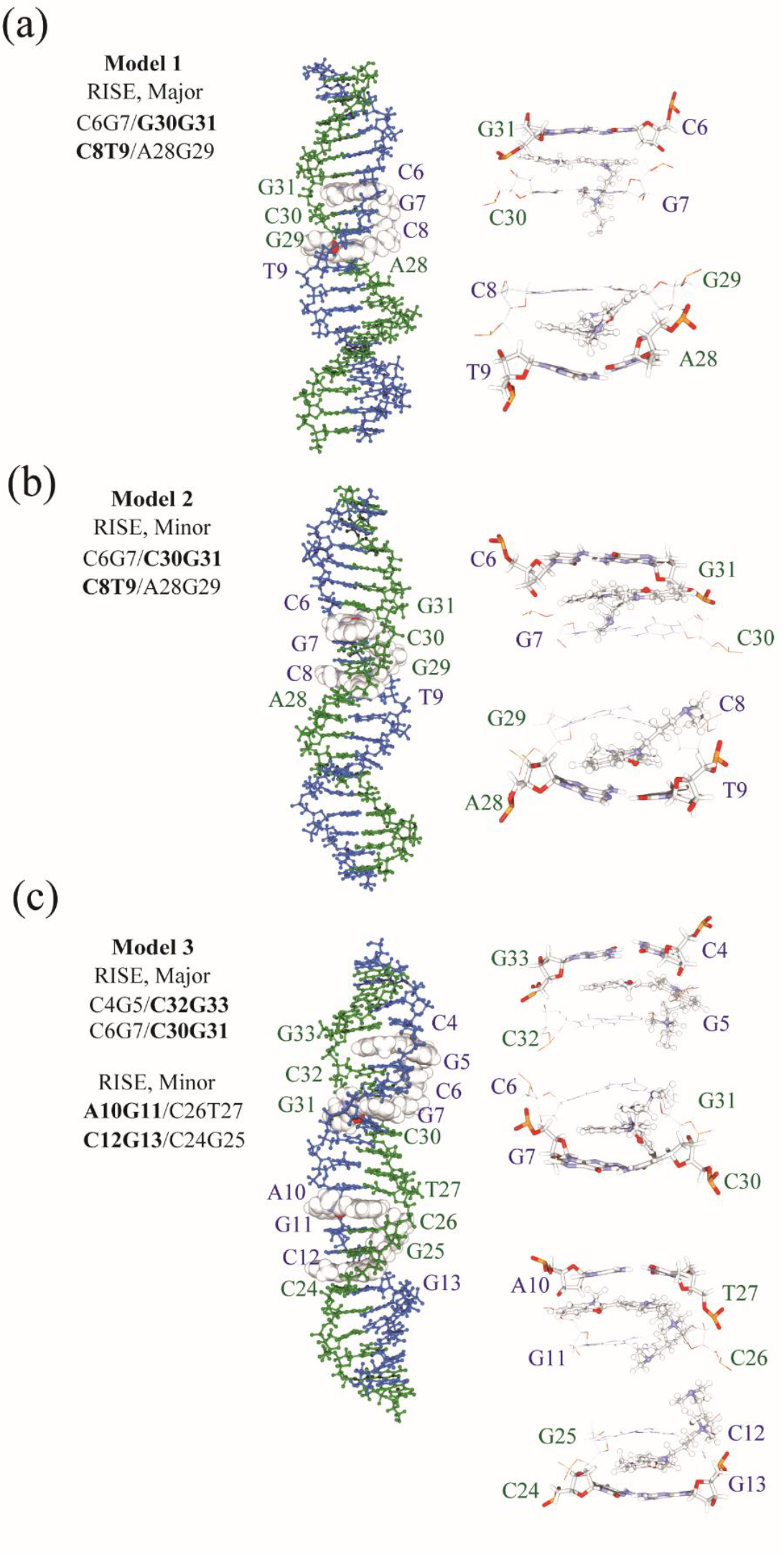
Atomic models of the intercalated DNA for DOXO, SYBR, and YOYO. A snapshot of the conformation of the selected models during the umbrella sampling simulation is depicted. (A) Models 1–6 (a–f) for DOXO are a snapshot of the window for *d*_fix_ = 60.0, 61,0, 61.0, 60.0, 57.0, 57.0 Å, which is nearest to *d*_min_, respectively. For model 1, *Twist* at 8:CG = 28.1°, 9:GA = 24.0°, 10:AA = 28.5°, and *Buckle* at 9:G-C = 4.5°, 10:A-T = – 1.3°. For model 2, *Twist* at 10:A = 34.2°, 11:AC = 40.7°, 12:CG = 24.5°, and *Buckle* at 11:A-T = 34.6°, 12:C-G = –24.9°. For model 3, *Twist* at 6:AA = 30.0°, 7:AC = 39.8°, 8:CG = 27.0°, and *Buckle* at 7:A-T = 27.0°, 8:C-G = –25.5°. For model 4, *Twist* at 12:CG = 25.7°, 13:GA = 44.6°, 14: AA = 25.3°, and *Buckle* at 13:G-C = 36.7°, 14:A-T = –33.1°. For model 5, *Twist* at 9:GA = 36.0°, 10:AC = 62.2°, 11:CG = 26.9°, and *Buckle* at 10:A-T = 13.7°, 11:C-G = –0.8°. For model 6, *Twist* at 9:GA = 38.3°, 10:AC = 52.7°, 11:CG = 26.1°, and *Buckle* at 10:A-T = –5.5°, 11:C-G = –2.7°. (B) Models 1–6 (a–f) for SYBR are for *d*_fix_ = 60.0, 60.0, 60.0, 60.0, 56.0, and 56.0 Å, respectively. For model 1, *Twist* at 6:AA = 32.3°, 7:AC = 2.8°, 8:CG = 44.9°, and *Buckle* at 7:A-T = 12.2°, 8:C-G = 9.7°. For model 2, *Twist* at 9:GA = 38.4°, 10:AA = 16.5°, 11:AC = 34.5°, and *Buckle* at 10:A-T = 3.6°, 11:A-T = –4.4°. For model 3, *Twist* at 6:AA = 33.5°, 7:AC = 40.8°, 8:CG = 33.7°, and *Buckle* at 7:A-T = 43.3°, 8:C-G = –23.7°. For model 4, *Twist* at 7:AC = 28.6°, 8:CG = 39.6°, 9:GA = 35.0°, and *Buckle* at 8:C-G = 34.9°, 9:G-C = –29.9°. For model 5, *Twist* at 6:AA = 24.4°, 7:AG = 40.2°, 8:GA = 46.8°, and *Buckle* at 7:A-T = 1.8°, 8:G-C = 12.8°. For model 6, *Twist* at 7:AC = 34.8°, 8:CA = 36.5°, 9:AA = 30.7°, and *Buckle* at 8:C-G = – 0.2°, 9:A-T = 2.5°. (C) Models 1–3 (a–c) for YOYO are for *d*_fix_ = 63.0, 63.0, and 70.0 Å, respectively. For Model 1, *Twist* at 5:GC = 32.2°, 6:CG = 0.9°, 7:GC = 37.5°, 8:CT = 28.7°, 9:TA = 26.7°, and *Buckle* at 6:C-G = –0.7°, 7:G-C = –0.5°, 8:C-G = 8.5°, 9:T-A = –34.0°. For Model 2, *Twist* at 5:GC = 28.8°, 6:CG = 21.3°, 7:GC = 25.7°, 8:CT = 38.6°, 9:TA = 37.7°, and *Buckle* at 6:C-G = 4.4°, 7:G-C = 14.3°, 8:C-G = 29.6°, 9:T-A = –22.6°. For Model 3 (first YOYO), *Twist* at 3:GC = 38.7°, 4:CG = 13.0°, 5:GC = 37.4°, 6:CG = 39.3°, 7:GC = 30.6°, and *Buckle* at 4:C-G = 26.6°, 5:G-C = 22.5°, 6:C-G = 33.4°, 7:G-C = –30.7°. For Model 3 (second YOYO), *Twist* at 9:TA = 48.8°, 10:AG = 11.8°, 11:GC = 34.2°, 12:CG = 39.9°, 13:GC = 32.9°, and *Buckle* at 10:A-T = 22.3°, 11:G-C = 0.6°, 12:C-G = 13.9°, 13:G-C = –25.9°.

First, we carried out ABMD simulations to extend and contract the intercalated DNA by ∼ 15 and ∼ 10 Å from the end-to-end distance of the intercalated DNA, respectively, and estimated preliminary free energy curves (FECs) for extension as a function of the end-to-end distance *d*. These FECs were subsequently refined using umbrella sampling simulations (See Methods). Figures 6(a) to 6(c) show FECs for RISE-type (For clarity, only models, 1, 3 and 5 are shown for DOXO and SYBR; individual models in Figure S2.). The lowest free energies were at *d* = 60.5 ± 0.23, 60.2 ± 0.13, 63.1 ± 0.10, and 70.1 Å for DOXO, SYBR, one-YOYO, and two-YOYO, respectively (Table II), while the equilibrium end-to-end distance of free dsDNA (without intercalators) was 56.8 Å for the sequence used with DOXO/SYBR and 56.7 Å for the sequence used with YOYO. This extension of approximately 3.4–3.7 Å per mono-intercalator is consistent with the insertion of one planar aromatic moiety between base pairs, corresponding to approximately one base-pair step height (∼3.2 Å). For YOYO, the extensions of 6.4 Å (one-YOYO) and 13.4 Å (two-YOYO) demonstrate additive contributions from each intercalated YOYO molecule. In contrast to RISE-type, the lowest free energies for OPEN-type occurred at *d* = 57.2 ± 0.05 (DOXO) and 56.2 ± 0.25 Å (SYBR), representing slight changes (0.4 Å and –0.6 Å, respectively) relative to free dsDNA. This indicates that in OPEN-type intercalation, the intercalator disrupts base stacking without significantly increasing the helical rise.

**Figure 6.**
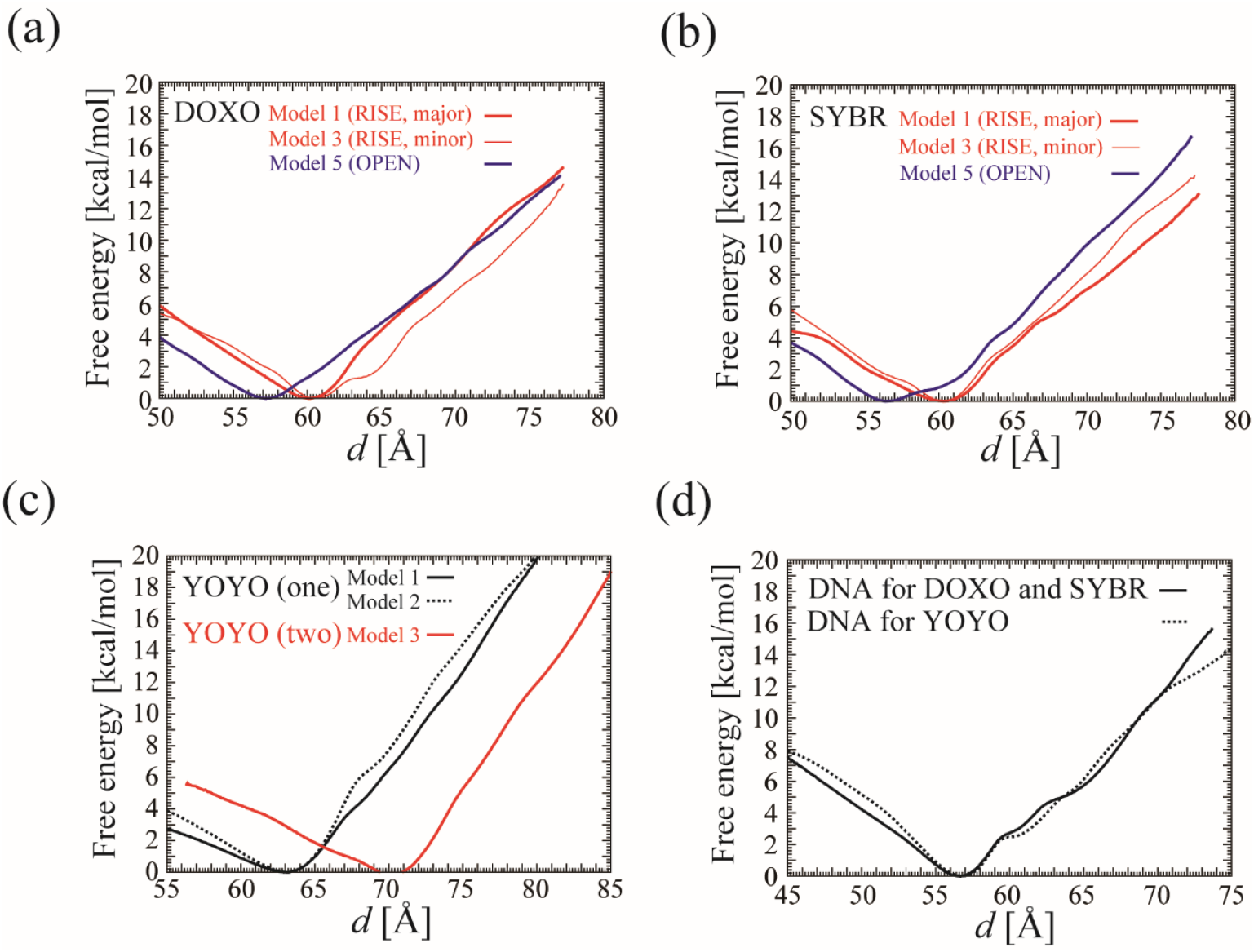
Free energy curve for the extension and contraction of the intercalated DNA. The FECs of the DNA intercalation against *d* (in units of Å) for (a) DOXO, (b) SYBR, (c) YOYO, and (d) the free dsDNA, respectively. The values of the free energies were set at zero at *d* = *d*_min_. For clarity, only models, 1, 3 and 5 are shown for DOXO and SYBR. Individual values for all the models are shown in Figure S2.

### Stretch moduli estimated from FECs

The FECs exhibit approximately quadratic curvature near the minimum *d*_min_ and approximately linear behavior at both sides. This indicates that the dynamics of the intercalated DNA is elastic around *d*_min_. In the quadratic region, we estimated the stretch modulus, which characterizes the elastic stiffness of the DNA, from the curvature of the FECs.

The stretch moduli for DNA extension were estimated to be (in units of 10^3^ pN): DOXO: 3.27 ± 0.66 (RISE-type) and 2.17 ± 0.34 (OPEN-type), SYBR: 2.31 ± 0.34 (RISE-type) and 1.45 ± 0.07 (OPEN-type), and YOYO: 2.46 ± 0.35 (one-YOYO, RISE-type) and 2.71 (two-YOYO, RISE-type) (Figure 7; individual model values in Figure S3). For reference, the stretch moduli which were estimated from the contour length (see Methods and Table SIV) are shown in Figure S4. It should be noted that they do not necessarily match the values in our study.

**Figure 7.**
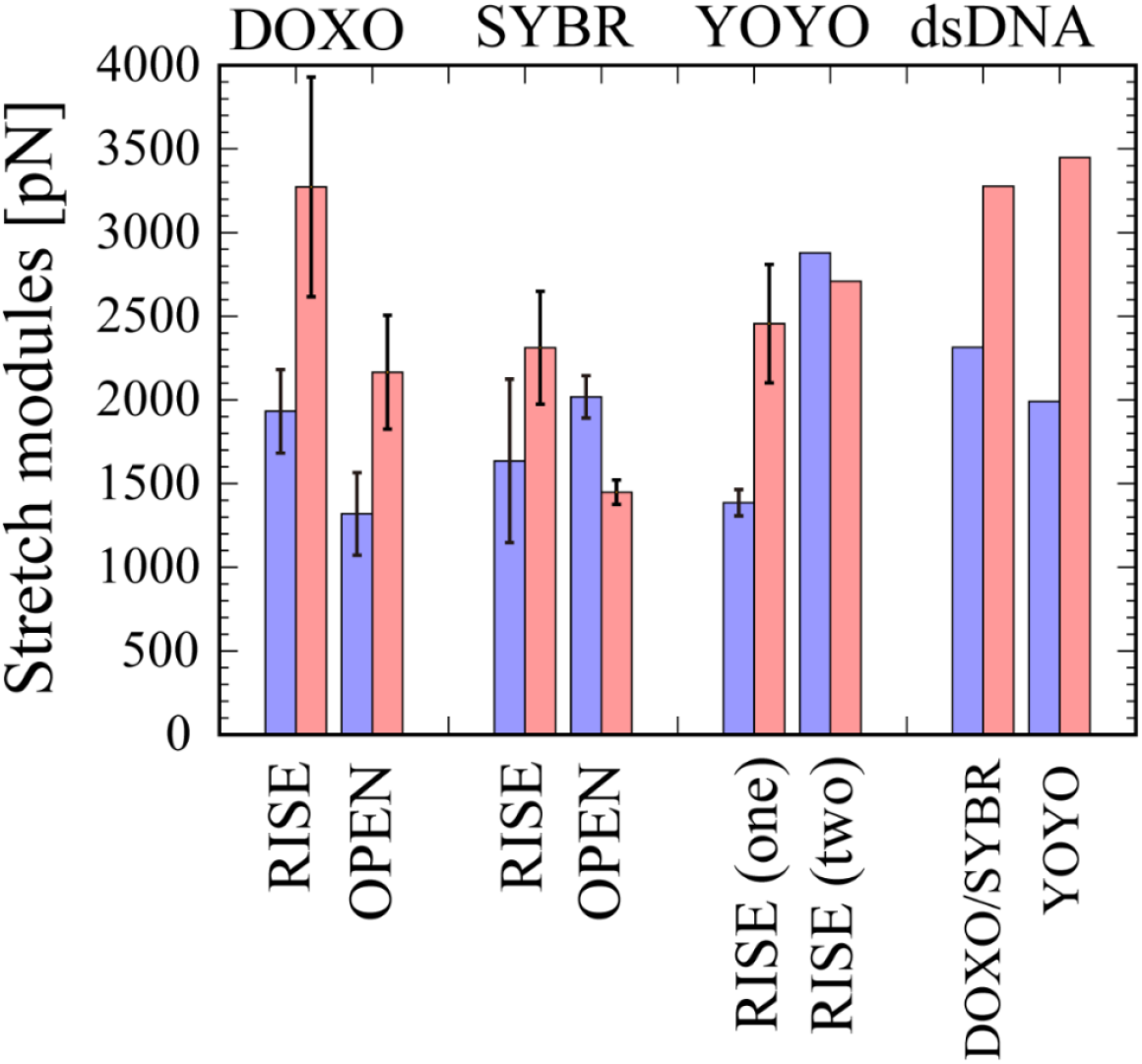
The stretch modulus for the intercalated DNA. The stretch module for extension (in red) and contraction (in blue) of the intercalated DNA for DOXO, SYBR, YOYO, and the free dsDNA are shown. See also Table SIV. Individual values for all the models are shown in Figure S3.

Most of the intercalated models showed reduced stretch moduli compared with free dsDNA (3.28 for the DOXO/SYBR sequences and 3.45 for the YOYO sequence), indicating that intercalation generally increases DNA flexibility during extension. However, DOXO models 2 and 3 (both RISE-type) showed larger stretch moduli. These models had *d*_min_ = 60.7 Å, compared with < 60.3 Å for other DOXO models, suggesting that these DNAs were already near-maximally extended at equilibrium. Consequently, further extension would require overcoming additional structural constraints, resulting in higher apparent stiffness.

The stretch moduli for DNA contraction were estimated to be (in units of 10^3^ pN): DOXO: 1.93 ± 0.25 (RISE-type) and 1.32 ± 0.25 (OPEN-type), SYBR: 1.64 ± 0.49 (RISE-type) and 2.02 ± 0.13 (OPEN-type), and YOYO: 1.39 ± 0.08 (one-YOYO, RISE-type) and 2.88 (two-YOYO, RISE-type) (Figure 7). Overall, OPEN-type consistently exhibited lower stretch moduli than RISE-type for both extension and contraction, reflecting their more disrupted base-stacking arrangements. Compared with free dsDNA (2.32 for DOXO/SYBR sequence and 1.99 for YOYO sequence), the higher contraction modulus for two-YOYO indicates that two-YOYO significantly restricts DNA flexibility for contraction. The asymmetry between extension and contraction moduli reflects the different molecular mechanisms governing DNA stretching versus compression in the presence of intercalators.

### Force in the elastic regime

Figure 8 shows the force extension as a function of *d*. For free dsDNA, the force increased linearly from 0 pN to a first peak marking the end of the elastic regime: 94.4 pN at *d* = 58.6 Å (DOXO/SYBR sequence) and 92.9 pN at *d* = 58.7 Å (YOYO sequence). For intercalated DNA, the first peak forces were shown in Figure 9 and Table SIV: DOXO: 78.3 ± 16.1 pN at d = 62.5 ± 0.2 Å (RISE-type) and 70.2 ± 22.9 pN at *d* = 59.3 ± 0.2 Å (OPEN-type), SYBR: 78.4 ± 10.2 pN at *d* = 62.5 ± 0.5 Å (RISE-type) and 36.3 ± 10.8 pN at *d* = 57.8 Å (OPEN-type), and YOYO: 114 ± 22.1 pN at *d* = 66.6 ± 0.15 Å (one-YOYO, RISE-type) and 117 pN at *d* = 73.6 Å (two-YOYO, RISE-type) (Individual model values in Figure S5).

**Figure 8.**
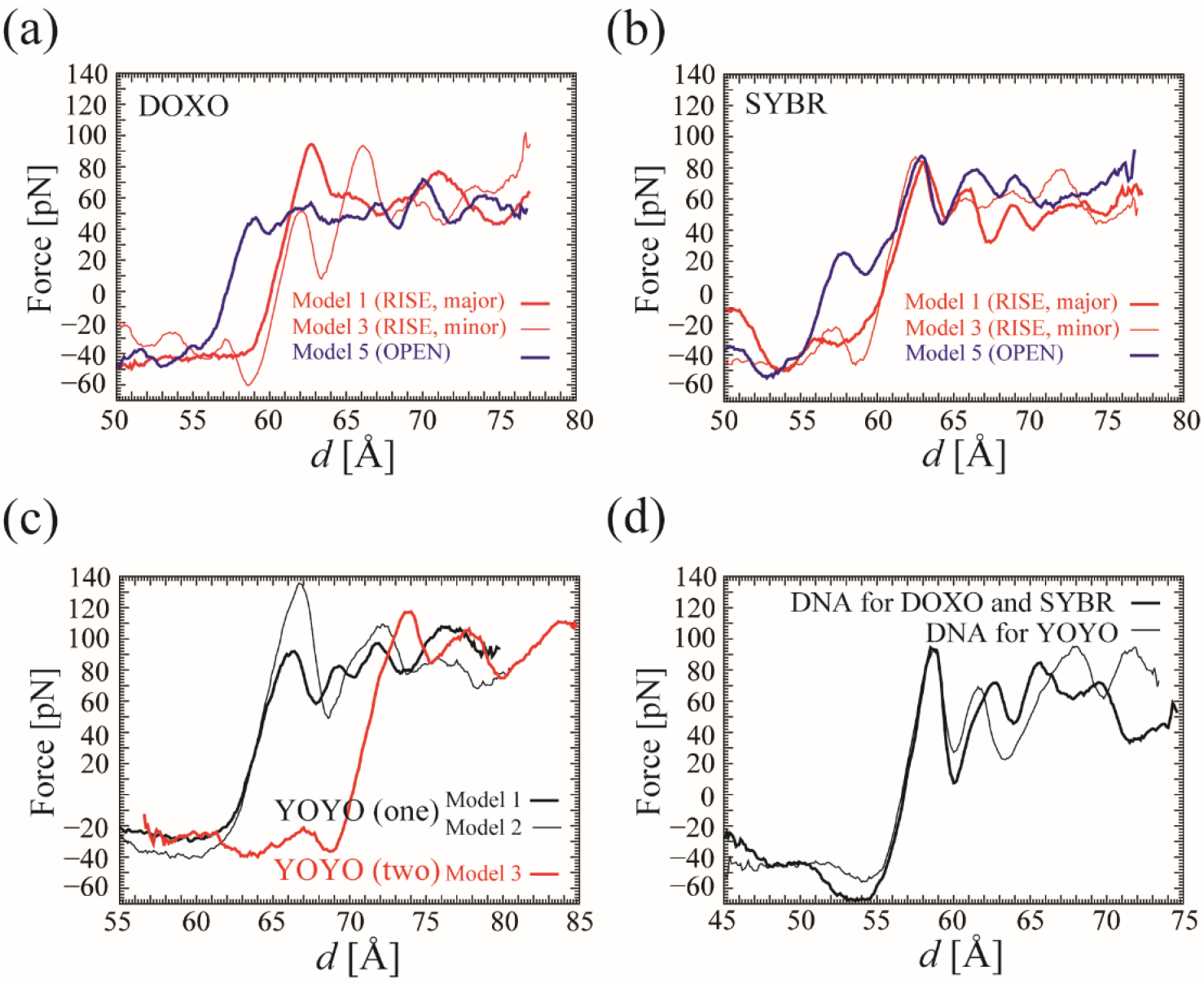
Force for the extension and contraction of the intercalated DNA along *d*. Force for the extension and contraction of the intercalated DNA along *d* for (a) DOXO, (b) SYBR, (c) YOYO, and (d) the free dsDNA, respectively. The force was calculated as the derivative of the free energy curve, obtained by the least-square fitting of seven data points (the current points with three preceding and three subsequent points) at the interval of 0.1 Å. For clarity, only models, 1, 3 and 5 are shown for DOXO and SYBR. Individual values for all the models are shown in Figure S5.

**Figure 9.**
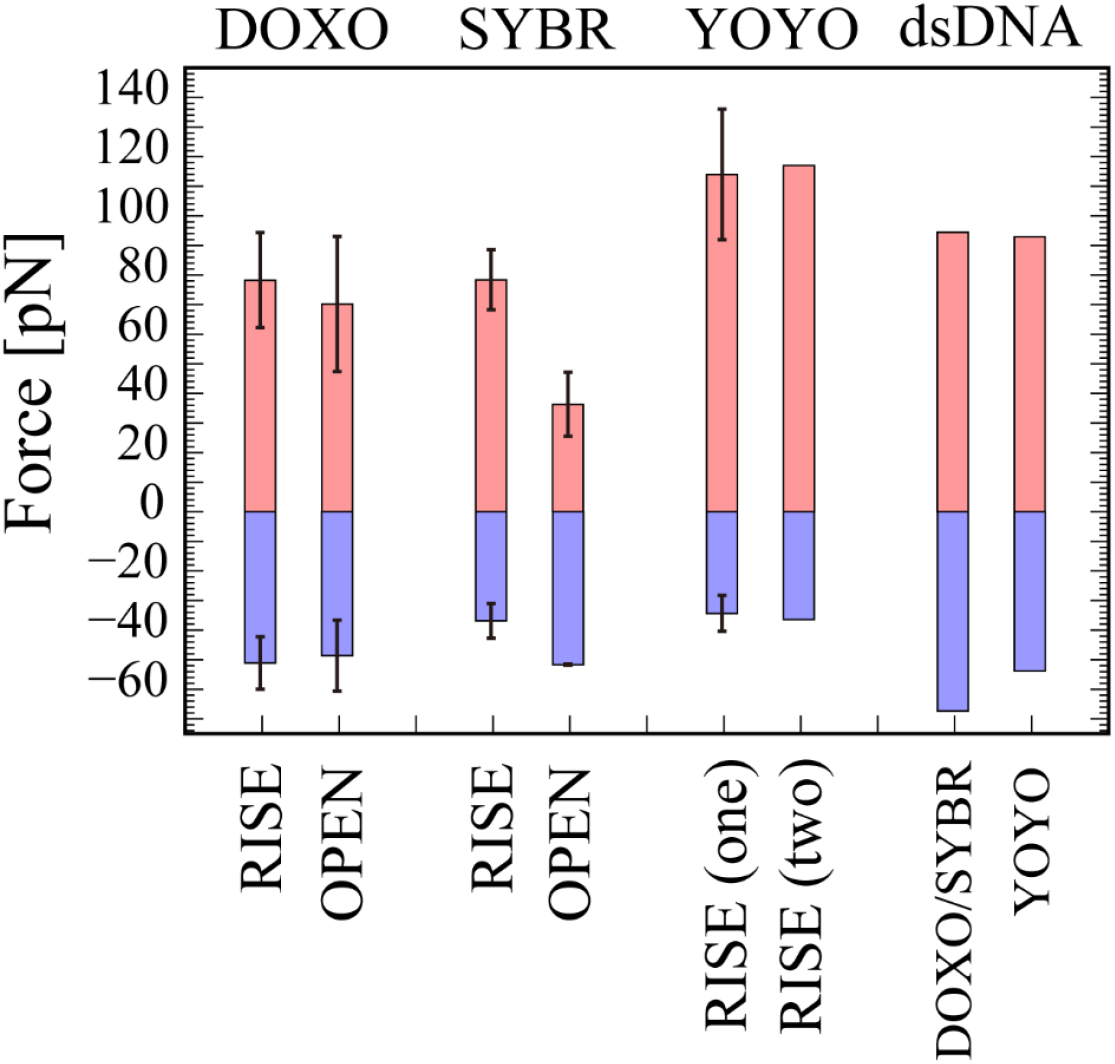
Force at the first peak for the extension and contraction of the intercalated DNA. The force at the first peak for the extension (in red) and contraction (in blue) of the intercalated DNA for DOXO, SYBR, and YOYO, and the free dsDNA are shown. See also Table SIV. Individual values for all the models are shown in Figure S6.

Notably, despite variations in stretch moduli among different models, the first peak forces remained relatively consistent within each intercalator type (DOXO and SYBR). This is because models with lower stretch moduli exhibit wider elastic ranges, while those with higher moduli show narrower ranges, such that the force × distance product at the elastic limit remains approximately constant. This suggests that intercalated DNA with lower stretch moduli can accommodate larger deformations before leaving the elastic regime, likely due to local structural rearrangements and stabilization of base pairs surrounding the intercalation site. The first peak forces for YOYO (>100 pN for both one-YOYO and two-YOYO intercalation) were notably higher than those for DOXO and SYBR (∼70–80 pN for RISE-type). This indicates that YOYO’s bis-intercalative structure with its flexible linker provides enhanced DNA stabilization compared with the mono-intercalators.

### Force during DNA overstretching in the inelastic regime

Beyond the first peak, the elastic response ceased, and forces began to fluctuate significantly. As shown in Figure S7, substantial structural deformation occurred in the inelastic region, including partial melting of the double-helix structure. However, base pairs around the intercalation site remained paired throughout the overstretching process. This indicates that the intercalation sites are structurally stable and do not easily collapse during extension. This is consistent with the experiments showing that intercalated YOYO molecules exhibit neither static reorientation nor alteration in their rotational dynamics during the force-induced DNA melting [54], and that melting is confined to domains between adjacent YOYO molecules during the overstretching transition [54].

Figure S8 shows the base-pair step entropy, an indicator of local structural stability. As DNA extended, entropy increased markedly at AT steps (positions 3, 6, 10, and 14), indicating that overstretching preferentially occurred at these sites. This is consistent with the experimental observation that the thermal melting of the DNA predominantly initiates at an AT step-rich sequence [55]. The drop in force at the overstretch transition corresponded roughly to the changes in local base-pair entropy. Furthermore, Figure S7 reveals that bases can rearrange into ladder-stacking configurations (where bases from opposite strands stack parallel rather than paired), which maintain or even increase extension forces. Thus, force fluctuations arise from the dynamic interplay between base-pair melting (force drops) and formation of alternative stacking arrangements (force recovery).

The averaged forces in the inelastic regime were estimated to be (in pN): DOXO: 60.6 ± 1.8 (RISE-type) and 53.6 ± 2.3 (OPEN-type), SYBR: 55.8 ± 3.2 (RISE-type) and 57.8 ± 1.4 (OPEN-type), and YOYO: 86.3 ± 1.6 (one-YOYO intercalation) and 96.4 (two-YOYO intercalation). Compared with free dsDNA (55.7 pN for DOXO/SYBR sequence and 65.3 pN for YOYO sequence), forces for DOXO and SYBR intercalated DNA remained similar to their respective free dsDNA, while YOYO-intercalated DNA showed significantly higher forces. This indicates that the intercalated DNA with YOYO is resistant to DNA overstretching. The elevated forces (>80 pN) for YOYO likely arise from the flexible linker connecting the two YO moieties, which mechanically couples the intercalated sites and resists DNA extension. This is particularly pronounced for bis-intercalated YOYO, where base pairs between the two intercalation sites are effectively clamped, requiring additional force to induce melting or structural rearrangement. These results demonstrate that while mono-intercalators (DOXO, SYBR) stabilize local structure without significantly affecting global overstretching mechanics, the bis-intercalator YOYO fundamentally alters DNA mechanical response in the inelastic regime.

### Forces during DNA compression and bending

Figure 9 and Table SIV show the forces at the first peak during DNA contraction in the elastic regime (in pN): DOXO: –51.1 ± 8.9 at *d* = 58.5 ± 0.2 Å (RISE-type) and –48.6 ± 12.0 at *d* = 54.2 ± 1.2 Å (OPEN-type), SYBR: –36.9 ± 5.9 at *d* = 58.1 ± 0.4 Å (RISE-type) and –51.7 ± 0.2 at *d* = 53.6 ± 0.1 Å (OPEN-type), and YOYO: –34.4 ± 6.1 at *d* = 60.4 ± 0.1 Å (one-YOYO, RISE-type) and –36.4 at *d* = 68.7 Å (two-YOYO, RISE-type). For comparison, free dsDNA showed forces of –67.3 pN (DOXO/SYBR sequence) and –53.8 pN (YOYO sequence). These contraction forces are lower in magnitude than the corresponding extension forces (78-117 pN, see previous section), indicating asymmetric mechanical responses. Importantly, the decrease in end-to-end distance *d* during compression results primarily from DNA bending rather than uniform helical compression, as visualized in Figure S9.

In the case of the RISE-type at the major groove, the intercalator adopts a wedge-like geometry that facilitates bending away from the major groove, with the intercalator at the convex side of the bend. In the case of the RISE-type at the minor groove, DNA preferentially bends toward the minor groove. The intercalator can be accommodated within the compressed minor groove, effectively stabilizing the bent conformation with the intercalator on the concave side. For the OPEN-type, intercalators act as wedges that induce bending in the direction opposite to the intercalation site, regardless of groove location. This occurs because OPEN-type intercalation disrupts base pairing without significantly extending the helix, creating an asymmetric hinge point.

In summary, intercalation not only reduces the forces required for DNA bending but also introduces directional preferences that depend on both the intercalation mode (RISE vs OPEN) and groove location (major vs minor).

### Local DNA unwinding at intercalation sites

Intercalators adopt two distinct orientations relative to adjacent base pairs at intercalation sites (Table SIV): parallel to the long axis of flanking base pairs, or perpendicular to it. These orientations result in different local unwinding patterns.

For DOXO, Model 1 (parallel orientation) exhibited local DNA unwinding of –14.7° at the intercalated base step (Figure S10A(a)). In contrast, Models 2-4 (perpendicular orientation) showed a different pattern: unwinding occurred at base-pair steps adjacent to the intercalation site (–8.0 ± 2.2° per adjacent step), while the intercalated step itself was overwound by 3.2 ± 1.0° (Figure S10A(b-d)). In the perpendicular orientation, base pairs flanking the intercalator exhibit significant buckling to wrap around the intercalator, as shown in Figure 5. This adjacent-step unwinding pattern is consistent with the crystal structure of d(CGATCG)_2_ with two DOXO molecules intercalated at CG steps in perpendicular orientation (PDB code: 1d12 [24]). For OPEN-type models (5 and 6, parallel orientation), DNA unwinding was more pronounced at –30.4 ± 12.4° (Figure S7A(e-f)).

For SYBR, Models 1 and 2 (parallel orientation) showed local DNA unwinding of – 15.0 ± 1.5° at the intercalated base step (Table SIV, Figure S10B(a-b)), while Models 3 and 4 (perpendicular orientation) showed unwinding at adjacent base-pair steps (– 4.3 ± 1.4° per adjacent base-pair step, Figure S10B(c-d)). OPEN-type Models 5 and 6 (parallel orientation) showed unwinding of –27.1 ± 3.1° (Figure S10B(e-f)).

For YOYO, Models 1 and 2 (parallel and perpendicular orientations) exhibited DNA unwinding of –14.5 ± 5.5° per one moiety of YOYO at the intercalated base steps (Figure S10C(a-b)). For Model 3 (two-YOYO), unwinding was –13.9 ± 5.8° per YO moiety (Figure S10C(c)), indicating additive contributions from each YOYO. This unwinding pattern is consistent with the crystal structure of d(5’-CGCTAGCG-3’)_2_ with TOTO (a YOYO analogue) at the minor groove in the parallel orientation. Notably, the orientation of the YO moiety at the minor groove in Models 2 and 3 is inverted relative to the TOTO crystal structure, likely due to differences in DNA sequence context.

### Global DNA structural parameters during extension

We calculated average *Rise* values as a function of *d* (Figure 10, for clarity only models 1, 3, 5 are shown for DOXO and SYBR; individual models are shown in Figure S11.) At the free energy minima, RISE-type intercalation yielded *Rise* values of 3.62 ± 0.02 Å (DOXO, Figure 10(a)), 3.62 ± 0.02 Å (SYBR, Figure 10(b)), 3.79 ± 0.01 Å (one-YOYO), and 4.21 Å (two-YOYO, Figure 10(c)), compared to 3.39 Å and 3.43 Å for free dsDNA (DOXO/SYBR and YOYO sequences, respectively, Figure 10(d)). Summed across all 17 base-pair steps, these per-step increases correspond to total *Rise* changes of 3.91 Å, 3.91 Å, 6.12 Å, and 13.3 Å for DOXO, SYBR, one-YOYO, and two-YOYO, respectively. For OPEN-type, *Rise* values were 3.39 ± 0.01 Å (DOXO) and 3.35 ± 0.01 Å (SYBR), representing essentially no change (0.00 and –0.04 Å) relative to free DNA.

**Figure 10.**
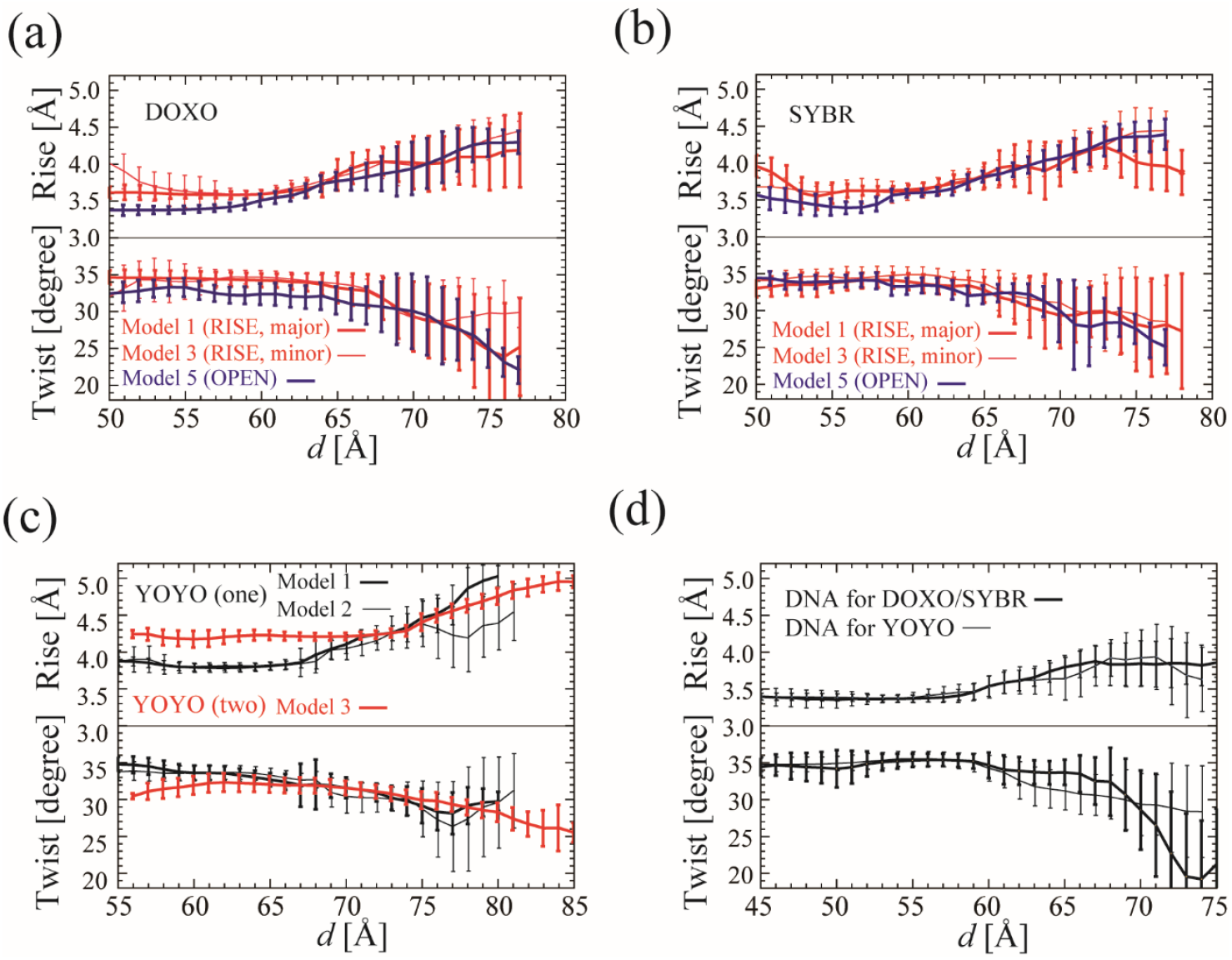
*Rise* and *Twist* of the intercalated DNA along *d*. *Rise* and *Twist* of the intercalated DNA along *d* for (a) DOXO, (b) SYBR, (c) YOYO, and (d) the free dsDNA, respectively. This value is averaged by 17 and 16 base-pair steps for RISE-type and Open-type, respectively. For clarity, only models, 1, 3 and 5 are shown for DOXO and SYBR. Individual values for all the models are shown in Figure S11.

Average *Twist* values were also calculated as a function of *d* (Figure 10). For RISE-type intercalation at the free energy minima, *Twist* values were, 34.5 ± 0.2°, 34.4 ± 0.4°, 33.5 ± 0.2°, and 31.6° for DOXO, SYBR, mono-YOYO, and bis-YOYO, respectively, compared with 35.4° and 35.5° for free dsDNA (DOXO/SYBR and YOYO sequences, respectively). These correspond to per-step *Twist* reductions of –0.9°, –1.0°, –2.0°, and –3.9° for the four intercalated systems. Summed across all 17 base-pair steps, the total *Twist* changes were –15.3°, –17.0°, –34.0°, and –66.3° for DOXO, SYBR, one-YOYO and two-YOYO, respectively. This demonstrates the additive effect of two YOYO molecules. For OPEN-type, *Twist* values were 33.2 ± 0.8° (DOXO) and 33.9 ± 0.1° (SYBR), showing per-step reductions of –2.2° and –1.5° relative to free DNA (35.4°). Across 16 base-pair steps, these correspond to total *Twist* changes of –35.2° and –24.0° for DOXO and SYBR, respectively.

Importantly, the relationship between *Twist* and end-to-end distance *d* was similar for free and intercalated DNA (Figure 10), indicating that DNA unwinding during extension is an inherent mechanical property of B-form DNA, independent of intercalator presence.

The total *Twist* changes for RISE-type intercalation were –15.3°, –17.0°, and –34.0° for DOXO, SYBR, and one-YOYO, respectively (ordered: DOXO < SYBR < YOYO). These values are in good agreement with single-molecule measurements: DOXO (–11° [7]) < SYBR (–19.1° [10]) < YOYO (–24° [12]), supporting our simulation approach. Notably, OPEN-type induced greater total unwinding (–24° to –35°) than RISE-type, despite causing minimal helical extension, reflecting the fundamentally different structural perturbations of these intercalation modes.

### Persistence length of intercalated DNA

In this study, the persistence length of free dsDNA was estimated to be 61.2 ± 3.2 nm for both sequences used (DOXO/SYBR and YOYO, Figure 11), which is in good agreement with previous MD simulations (∼30–60 nm [14] [45] [46]) and experimental measurements (∼45–55 nm [56]).

**Figure 11.**
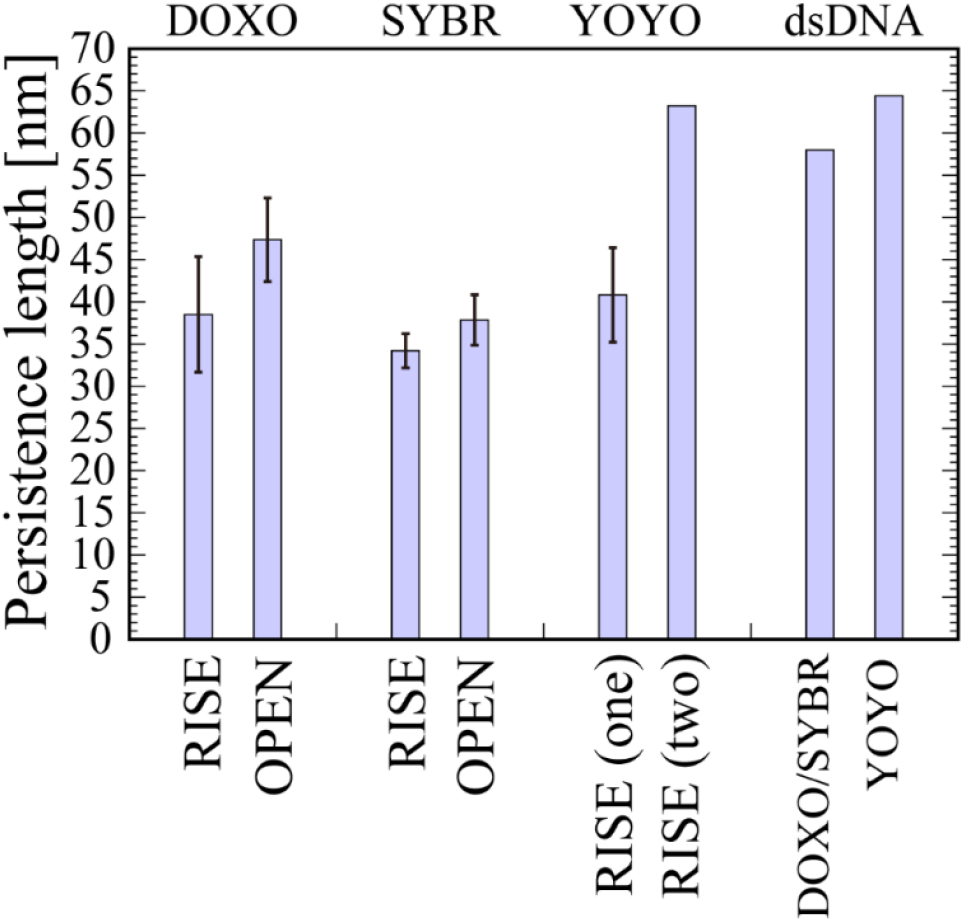
The persistence lengths of the intercalated DNA. The persistence lengths of the intercalated DNA for DOXO, SYBR, and YOYO, and the free dsDNA are shown. See also Table SIV. Individual values for all the models are shown in Figure S12.

The persistence length of intercalated DNA with DOXO and SYBR was significantly reduced to 38.5 ± 6.9 nm (DOXO, RISE-type), 47.4 ± 5.0 nm (DOXO, OPEN-type), 34.2 ± 2.0 nm (SYBR, RISE-type), and 37.9 ± 3.0 nm (SYBR, OPEN-type), representing 37– 77% of the free dsDNA value (individual model values in Figure S12). For YOYO, one-YOYO reduced the persistence length to 40.8 ± 5.6 nm (67% of free DNA), while two-YOYO restored it to 63.2 nm, comparable to free dsDNA (61.2 nm). These results indicate that mono-intercalators (DOXO, SYBR) and one-YOYO increase DNA flexibility, while two-YOYO rigidifies the DNA. This will be discussed in the following subsection.

### Energetics of DNA intercalation: MM-PBSA/MM-GBSA analysis

During intercalation, adjacent base pairs at the intercalation site must deform to accommodate the intercalator. This deformation requires energy to unstack base pairs and alter the sugar-phosphate backbone conformation for cavity formation [17] (see Method and Eq.(9)). These deformation energies are offset by favorable stacking interaction between the intercalator and adjacent base pairs, as well as electrostatic interactions between the intercalator’s positive charges and the phosphate’s negative charges.

Table II shows the contribution of the van der Waals interactions, electrostatic interactions, and solvation for intercalation, using the MM/PBSA and MM/GBSA methods. Here we discuss the MM/PBSA results. The MM/GBSA results show similar trends and are not discussed separately.

For DOXO RISE-type, the deformation energy components were: gas-phase van der Waals interaction energy (*E*_def_(VDW)): 23.4 ± 1.8 kcal/mol (unfavorable); gas-phase electrostatic interaction energy (*E*_def_(ELE)): –310.1 ± 44.6 kcal/mol (favorable); electrostatic polar solvation free energy (*G*_def_(PBCAL)): 293.8 ± 45.8 kcal/mol (unfavorable, opposing the electrostatic gain); net electrostatic contribution (*G*_def_(PBELE)): –16.4 ± 1.2 kcal/mol. These result in the positive total deformation energy *G*_def_(PBTOT): 14.8 ± 4.1 kcal/mol (Table II(a)), indicating that creating the intercalation cavity requires an energetic penalty. For OPEN-type, *E*_def_(VDW) (31.4 ± 2.0 kcal/mol) was larger than RISE-type because OPEN-type breaks two base-pair steps rather than one. Similarly, *E*_def_(ELE) (–28.3 ± 33.5 kcal/mol) was less favorable than RISE-type due to hydrogen bond breakage between the adjacent base pairs. The resulting *G*_def_(PBELE) (–13.6 ± 1.4 kcal/mol) and *G*_def_(PBTOT) (21.0 ± 0.8 kcal/mol) were both higher than RISE-type, indicating greater energetic cost for cavity formation.

**Table II(a).** The summary of MM-PBSA/GBSA for DOXO.

| Intercalator<br>DOXO | $E_{def}$<br>(VDW) | $E_{def}$<br>(ELE) | $G_{def}$<br>(PBCAL) | $G_{def}$<br>(PBELE) | $G_{def}$<br>(PBTOT) | $dE_{corr}$<br>(VDW) | $dG_{corr}$<br>(ELE) | $dG_{corr}$<br>(PBELE) | $dG_{corr}$<br>(PBTOT) | $d_{fix}$ [Å] |
| --- | --- | --- | --- | --- | --- | --- | --- | --- | --- | --- |
| Model1 | 21.0 | -316.6 | 300.2<br>(298.9) | -16.4<br>(-17.7) | 9.2<br>(7.4) | -27.5 | -940.7 | -1.3<br>(-8.0) | -28.9<br>(-33.6) | 60.0 (60.2) |
| Model2 | 26.0 | -364.8 | 350.3<br>(348.4) | -14.6<br>(-16.5) | 20.7<br>(18.1) | -27.2 | -985.0 | 0.29<br>(-19.7) | -22.7<br>(-40.7) | 61.0 (60.7) |
| Model3 | 23.9 | -318.6 | 302.1<br>(301.3) | -16.5<br>(-17.3) | 14.2<br>(12.9) | -38.7 | -1010.4 | 5.4<br>(-17.0) | -31.9<br>(-51.9) | 61.0 (60.7) |
| Model4 | 22.8 | -240.4 | 222.5<br>(225.7) | -17.9<br>(-14.7) | 15.2<br>(18.0) | -40.5 | -929.2 | 5.4<br>(-15.5) | -30.5<br>(-49.0) | 60.0 (60.3) |
| Model5 | 29.3 | 5.2 | -17.4<br>(-1.2) | -12.2<br>(4.1) | 20.2<br>(35.9) | -27.9 | -689.6 | 1.2<br>(-1.8) | -29.0<br>(-30.1) | 57.0 (57.2) |
| Model6 | 33.4 | -61.7 | 46.8<br>(57.7) | -14.9<br>(-4.0) | 21.9<br>(32.1) | -28.2 | -769.7 | -0.1<br>(-10.7) | -30.7<br>(-39.3) | 57.0 (57.1) |

The unit of the energies is kcal/mol. The values in parentheses for the columns of free energies are free energies calculated by MM-GBSA. The values in parentheses for the column of $d_{fix}$ are $d_{min}$ .

For SYBR, RISE-type showed *E*_def_(VDW) = 24.0 ± 1.3, *G*_def_(PBELE) = –15.0 ± 1.3, and *G*_def_(PBTOT) = 16.1 ± 2.1 kcal/mol, all lower than corresponding OPEN-type values (30.1 ± 0.1, –10.7 ± 0.4, and 24.5 ± 1.7 kcal/mol, respectively; Table II(b)), consistent with the DOXO pattern.

For one-YOYO, *E*_def_(VDW) (42.5 ± 1.8 kcal/mol; Table III(c)) was about double that of DOXO and SYBR, reflecting the larger intercalating moiety. *G*_def_(PBTOT) (21.4 ± 5.2 kcal/mol) was comparable with SYBR RISE-type. For two-YOYO, *E*_def_(VDW) (88.4 kcal/mol) approximately doubled again as expected for double of one-YOYO, while *G*_def_(PBTOT) (55.5 kcal/mol) more than doubled, indicating non-additive effects from accommodating both molecules.

**Table II(b).** The summary of MM-PBSA/GBSA for SYBR.

| Intercalator<br>SYBR | $E_{def}$<br>(VDW) | $E_{def}$<br>(ELE) | $G_{def}$<br>(PBCAL) | $G_{def}$<br>(PBELE) | $G_{def}$<br>(PBTOT) | $dE_{corr}$<br>(VDW) | $dG_{corr}$<br>(ELE) | $dG_{corr}$<br>(PBELE) | $dG_{corr}$<br>(PBTOT) | $d_{fix}$ [Å] |
| --- | --- | --- | --- | --- | --- | --- | --- | --- | --- | --- |
| Model 1 | 26.1 | -313.6 | 296.7<br>(299.5) | -16.9<br>(-14.1) | 13.5<br>(15.7) | -28.3 | -1563.8 | -7.8<br>(-11.1) | -37.3<br>(-38.3) | 60.0 (60.3) |
| Model 2 | 24.0 | -312.7 | 297.7<br>(298.6) | -15.0<br>(-14.1) | 16.8<br>(17.1) | -24.5 | -1562.8 | -10.8<br>(-16.5) | -32.4<br>(-36.0) | 60.0 (60.1) |
| Model 3 | 23.8 | -298.0 | 284.7<br>(282.9) | -13.3<br>(-15.2) | 15.1<br>(12.6) | -38.3 | -1649.5 | -0.9<br>(-19.8) | -40.6<br>(-52.3) | 60.0 (60.3) |
| Model 4 | 22.3 | -270.7 | 256.0<br>(265.1) | -14.8<br>(-5.7) | 19.0<br>(27.7) | -41.3 | -1648.8 | -1.1<br>(-14.4) | -37.1<br>(-48.1) | 60.0 (60.0) |
| Model 5 | 30.0 | 3.7 | -14.8<br>(0.2) | -11.1<br>(3.9) | 22.8<br>(37.3) | -36.0 | -1361.9 | -3.8<br>(0.5) | -41.9<br>(-35.3) | 56.0 (56.4) |
| Model 6 | 30.2 | -35.4 | 25.1<br>(44.8) | -10.2<br>(9.5) | 26.2<br>(45.4) | -36.0 | -1427.1 | -0.0<br>(8.4) | -36.1<br>(-25.0) | 56.0 (55.9) |

The unit of the energies is kcal/mol. The values in parentheses for the columns of free energies are free energies calculated by MM-GBSA. The values in parentheses for the column of $d_{fix}$ are $d_{min}$ .

**Table II(c).** The summary of MM-PBSA/GBSA for YOYO.

| Intercalator<br>YOYO | $E_{def}$<br>(VDW) | $E_{def}$<br>(ELE) | $G_{def}$<br>(PBCAL) | $G_{def}$<br>(PBELE) | $G_{def}$<br>(PBTOT) | $dE_{corr}$<br>(VDW) | $dG_{corr}$<br>(ELE) | $dG_{corr}$<br>(PBELE) | $dG_{corr}$<br>(PBTOT) | $d_{fix}$ [$\text{\AA}$] |
| --- | --- | --- | --- | --- | --- | --- | --- | --- | --- | --- |
| Model 1 (Major) | 40.7 | -527.8 | 495.9 (499.9) | -31.9 (-26.5) | 16.2 (19.5) | -56.6 | -3037.4 | -25.4 (-43.0) | -82.9 (-97.4) | 63.0 (63.0) |
| Model 2 (Minor) | 44.3 | -538.2 | 507.4 (508.7) | -30.8 (-29.5) | 26.5 (27.0) | -50.2 | -3115.3 | -20.4 (-53.7) | -65.8 (-96.3) | 63.0 (63.2) |
| Model 3 (Major<br>and Minor) | 88.4 | -968.9 | 910.0 (928.6) | -58.9 (-40.4) | 55.5 (72.3) | -102.4 | -5814.2 | -0.7 (77.7) | -95.0 (-9.7) | 70.0 (70.1) |
| Major at first | 88.4 | -968.9 | 910.0 (928.6) | -58.9 (-40.4) | 55.5 (72.3) | -7.7 | -3490.8 | -46.8 (-67.4) | -36.9 (-55.6) | 70.0 (70.1) |
| Minor at second | 0.0 | 0.0 | 0.0 | 0.0 | 0.0 | -94.8 | -2123.4 | 45.2 (145.2) | -58.0 (45.9) |  |
| Subtotal |  |  |  |  |  |  |  | -1.6 (77.8) | -94.9 (-9.6) |  |
| Minor at first | 88.4 | -968.9 | 910.0 (928.6) | -58.9 (-40.5) | 55.5 (72.3) | -6.4 | -3288.8 | -50.1 (-55.0) | -38.8 (-41.6) | 70.0 (70.1) |
| Major at second | 0.0 | 0.0 | 0.0 | 0.0 | 0.0 | -96.0 | -2325.4 | 48.5 (132.8) | -56.1 (32.0) |  |
| Subtotal |  |  |  |  |  |  |  | -1.6 (77.8) | -94.9 (-9.6) |  |

The unit of the energies is kcal/mol. The values in parentheses for the columns of free energies are free energies calculated by MM-GBSA. The values in parentheses for the column of $d_{fix}$ are $d_{min}$ .

Next, we calculated binding free energies (Δ*G*_corr_) by adding *G*_def_ to the raw binding energy (Δ*G*_bind_), which accounts for DNA deformation. For DOXO, Δ*G*_corr_(PBTOT) for RISE-type and OPEN-type were –28.5 ± 3.5 kcal/mol and –29.9 ± 0.85 kcal/mol, respectively. These negative values were large because Δ*G*_bind_(PBTOT), Δ*G*_corr_(PBTOT) – *G*_def_(PBTOT) = –43.3 ± 3.2 and –50.9 ± 1.7 kcal/mol, respectively, were much larger in magnitude than *G*_def_(PBTOT), indicating that the deformation energy was overcome by the favorable binding energy. These Δ*G*_corr_(PBTOT) values are close to the gas-phase van der Waals energies, Δ*E*_corr_(VDW) (–33.5 ± 6.1 kcal/mol for RISE-type and – 28.1 ± 0.2 kcal/mol for OPEN-type), indicating that the van der Waals interactions are the major driving force for intercalation. The same pattern holds for SYBR and YOYO. For SYBR, Δ*G*_corr_(PBTOT) for RISE-type and OPEN-type were –36.8 ± 2.9 and – 39.0 ± 2.9 kcal/mol, respectively and the corresponding Δ*E*_corr_(VDW) were –33.1 ± 6.9 and –36.0 kcal/mol, respectively. For YOYO, Δ*G*_corr_(PBTOT) were –74.4 ± 8.6 and – 94.9 kcal/mol for one-YOYO and two-YOYO, respectively and Δ*E*_corr_(VDW) were – 53.4 ± 3.2 and –102 kcal/mol, respectively.

### Intercalation preference: RISE-type at minor groove > at major groove ≈ OPEN-type

For DOXO and SYBR RISE-type intercalation, Δ*G*_corr_(PBTOT) values are less favorable at the major groove (DOXO/SYBR Models 1 and 2) compared with the minor groove (Models 3 and 4), indicating that the minor groove intercalation is thermodynamically preferred due to more favorable van der Waals interactions.

For OPEN-type intercalation, the deformation energy *G*_def_(PBTOT) was higher than RISE-type: 21.1 ± 0.8 kcal/mol (DOXO, OPEN-type) and 24.5 ± 1.7 kcal/mol (SYBR, OPEN-type) versus 14.8 ± 4.1 kcal/mol (DOXO, RISE-type) and 16.1 ± 2.1 kcal/mol (SYBR, RISE-type). However, Δ*G*_corr_(PBTOT) for OPEN-type were comparable to RISE-type at the major groove: –29.9 ± 0.9 kcal/mol (DOXO, OPEN-type) and – 39.0 ± 2.9 kcal/mol (SYBR, OPEN-type) versus –28.5 ± 3.5 kcal/mol (DOXO, RISE-type) and –36.8 ± 2.9 kcal/mol (SYBR, RISE-type). This is because the free energy of the DNA – intercalator complex, *G*_complex_(PBTOT) (in Eq.(8)), is more favorable for OPEN-type, likely due to stabilizing interactions between the intercalator and the disrupted (flipped-out) bases, which offset the higher deformation cost. These results suggest that OPEN-type intercalation is thermodynamically competitive with RISE-type intercalation at the major groove, despite requiring greater DNA deformation.

In summary, intercalation follows the thermodynamic preference order: RISE-type at minor groove (most favorable, lowest Δ*G*_corr_) > RISE-type at major groove ≈ OPEN-type, consistent with the predominance of minor groove RISE-type conformations observed in our simulations.

### Non-cooperative binding in two-YOYO intercalation

Although the van der Waals contribution for two-YOYO (Δ*E*(VDW) = 88.4 kcal/mol) is about double that of one-YOYO (42.5 ± 1.8 kcal/mol), the electrostatic contribution shows a striking deviation: Δ*G*(PBELE) = –1.7 kcal/mol for two-YOYO is far less favorable than double the ono-YOYO value (–22.9± 2.5 kcal/mol). This indicates that the additional intercalation of YOYO is hindered not by stacking interactions, but rather by unfavorable electrostatic repulsion between the two positively charged YOYO molecules. Consequently, Δ*G*_corr_(PBTOT) for two-YOYO (–94.9 kcal/mol) is substantially less favorable than twice the one-YOYO value (2 × –74.4 ± 8.6 kcal/mol = –148.8 kcal/mol), demonstrating that two-YOYO intercalation is thermodynamically suppressed by one-YOYO. This non-cooperative binding contrasts sharply with previous MD simulations of bis-intercalating doxorubicin and daunomycin[17] [19], where Δ*G*_corr_ for the second intercalation was more favorable than expected, indicating cooperative binding effects[17] [19].

To confirm that non-cooperativity arises from electrostatic repulsion between YOYO molecules, we analyzed Δ*G*_corr_ for sequential intercalation steps: (1) first intercalation at the major groove and (2) the second intercalation at the minor groove and vice versa. The former yielded Δ*G*_corr_first_major_(PBTOT) = –36.9 kcal/mol for the first step and – 58.0 kcal/mol for the second, totaling Δ*G*_corr_(PBTOT) = –94.9 kcal/mol. The latter yielded –38.8 kcal/mol for the first and –56.1 kcal/mol for the second, again totaling – 94.9 kcal/mol. These results demonstrate that while the first YOYO intercalation stabilizes DNA electrostatically, the second intercalation experiences reduced binding affinity due to electrostatic repulsion between the two positively charged intercalators. The asymmetry in Δ*G*_corr_(PBELE) between first and second intercalation steps directly reflects this intermolecular repulsion (Table II(c)).

The non-cooperative effect is further evidenced by DNA extension patterns: one-YOYO (Model 1) extends free DNA by 6.3 Å (from d_min_ = 56.7 Å to 63.0 Å), while two-YOYO (Model 4) extends the already-intercalated DNA by 7.1 Å (from 63.0 Å to 70.1 Å). The greater extension required for the second intercalation (7.1 Å vs 6.3 Å) suggests increased structural strain, consistent with electrostatic repulsion forcing the two intercalators further apart.

In summary, electrostatic repulsion between positively charged YOYO molecules produces non-cooperative bis-intercalation, where the second binding event is thermodynamically less favorable than the first. This distinguishes YOYO from smaller intercalators like doxorubicin that show cooperative binding, likely due to YOYO’s higher charge.

## Discussion

### DNA sequence preference of intercalators:DOXO, SYBR, and YOYO

DOXO has been reported to prefer CG steps in solution [57] [58], yet experimental studies show comparable affinity for poly AT and poly CG sequences [26] [59]. In the crystal structure of d(CGATCG)_2_ complexed with DOXO (PDB: 1d12 [60]), DOXO intercalates at the CG step in the minor groove—a configuration rarely observed in our simulations. This discrepancy likely reflects differences between crystallization conditions and the solution environment modeled in our MD simulations. Importantly, intercalation can be affected not only by the adjacent base pairs, but also by those located downstream base pairs with which the DOXO amino group interacts [18]. Kellogg et al. systematically investigated DOXO binding across 64 base-pair quartet combinations (CAxy, CGxy, TAxy, TGxy, where x and y are A, C, G or T) and demonstrated highly complex sequence specificity [61]. These findings support our observation that DOXO exhibits moderate rather than strong sequence preferences.

In our analysis of ABMD-generated structures, we focused on base pairs directly at the intercalation site rather than including downstream quartets, as the latter would exponentially increase the parameter space. The quartet sequences in our Models 1-4 (5’-G9A10A11C12-3’, 5’-A11C12G13A14-3’, 5’-A7C8G9A10-3’, and 5’-T23C24G25T26-3’, respectively) fall outside Kellogg et al.’s analyzed set. Nevertheless, Models 3 and 4 (RISE-type at the minor groove) exhibited the lowest MM-PBSA/GBSA free energies, strongly suggesting these represent thermodynamically favorable intercalation modes.

To the best of our knowledge, no crystal structure of DNA–SYBR has been reported, nor have specific DNA sequence preferences been experimentally identified. This absence of strong sequence specificity in the literature is consistent with our simulation results, which show SYBR intercalating at multiple sequence contexts with comparable binding energies. The lack of strong sequence discrimination may contribute to SYBR’s utility as a general nucleic acid stain.

For YOYO, the crystal structure of d(5’-CGCTAGCG-3’)_2_ with TOTO (PDB: 108d [25]), a YOYO analogue, shows bis-intercalation at CT and AG steps in the minor groove. While we did not observe it at the C8T9/A10G11 steps, we did observe it at three alternative sites: C6G7/C8T9, A10G11/C12G13, and C4G5/C6G7. Notably, the AG (=CT) and CG steps where YOYO intercalates exhibit lower intrinsic *Twist* values in free B-DNA compared with GC steps (Figure S10C(d)). This correlation suggests that YOYO preferentially intercalates at base-pair steps with inherently low Twist angles, which may require less energetic penalty for DNA unwinding during intercalation. The flexible linker connecting YOYO’s two YO moieties may allow adaptation to multiple sequence contexts while maintaining this preference for low-Twist steps.

### Intercalation kinetics and comparison with experimental data

Our calculated *k*_on_ for YOYO bis-intercalation was substantially lower than for mono-intercalation by DOXO and SYBR, consistent with experimental measurements: YOYO (3.3×10^5^ M^-1^s^-1^ [62]) shows relatively slow association kinetics as compared with SYBR (2.1×10^6^ – 1.2×10^7^ M^-1^s^-1^ [9]) and DOXO (7.0×10^6^ M^-1^s^-1^ [43]). The experimental *k*_on_ for YO (3.2×10^7^ M^-1^s^-1^) is about 100-fold faster than for YOYO (3.3×10^5^ M^-1^s^-1^ [9]), indicating that the first YO moiety’s intercalation does not facilitate the second moiety’s binding. The linker connecting the two YO moieties likely restricts conformational freedom for achieving optimal positioning of the second moiety for intercalation. This geometric constraint is compounded by electrostatic repulsion between the positively charged moieties. This analysis of the non-cooperative YO moiety’s intercalation is similar to our energetic analysis, which showed non-cooperative binding in YOYO intercalation. Our simulations reproduce the experimental kinetic order: mono-YO > SYBR > DOXO > bis-YOYO, supporting the validity of our ABMD approach for capturing relative intercalation dynamics.

While our simulations correctly reproduce the qualitative kinetic order, the calculated *k*_on_ values are 1–3 orders of magnitude faster than experimental measurements (Table SIII). Several factors contribute to this discrepancy. First, the small simulation box with periodic boundary conditions creates effective intercalator concentrations (2,970 mM) 3 to 6 orders of magnitude exceeding experimental conditions (typically µM–mM range), artificially accelerating encounter rates. Second, measured kinetics are highly sensitive to DNA length, intercalator concentration, salt type and concentration, pH, and solvent conditions [9] [13] [62] [43], making direct quantitative comparison challenging. Finally, our operational definition of the intercalated state (see Methods) necessarily differs from experimental criteria. Rigorous comparison would require free energy barrier calculations and transition state analysis between groove-bound and intercalated states [15] [16], which lies beyond the scope of this study. Despite these quantitative discrepancies, the consistent order of relative rates across all three intercalators—and the correct prediction that YOYO bis-intercalation is slowest—demonstrates that our ABMD approach successfully captures the fundamental physical factors governing intercalation kinetics. The simulations provide mechanistic insights into why these kinetic differences arise, particularly the role of molecular size, charge distribution, and linker constraints in determining association rates.

### Unwinding of the DNA by intercalation

Intercalation induces DNA unwinding through two distinct mechanisms: unwinding at the intercalation site itself or unwinding at adjacent base-pair steps. The mechanism depends on intercalator orientation. When DOXO and SYBR orient parallel to the long axis of flanking base pairs, unwinding occurs primarily at the intercalation site. When oriented perpendicular, unwinding occurs at adjacent base-pair steps instead. (For YOYO, unwinding at the intercalation site predominates regardless of orientation.)

Parallel orientation optimizes π-π stacking interactions between the intercalator and adjacent base pairs, concentrating the structural perturbation at the intercalation site. Perpendicular orientation, in contrast, distributes the structural strain to adjacent steps, possibly to minimize backbone distortion at the intercalation site itself. The observed orientation and unwinding pattern likely reflect an energetic balance between maximizing base-intercalator stacking and minimizing backbone strain.

Increased fluctuations in base-pair step parameters at intercalation sites (Figure S10) indicate enhanced local structural flexibility, which expands the elastic range of *d*. When buckling occurs, this structural perturbation propagates to induce conformational changes in neighboring base pairs.

### Possible interconversion between RISE-type and OPEN-type conformations

Our MM-PBSA/GBSA analysis revealed comparable binding free energies for RISE-type at the major groove and OPEN-type conformations, suggesting thermodynamic accessibility of both states. Direct evidence for this interconversion appeared when DOXO model 1 and SYBR model 1 for the windows at *d*_fix_ = 61.0 Å and 51.0 Å, respectively, spontaneously transitioned from RISE-type to OPEN-type during umbrella sampling, demonstrating that the energy barrier between these states is surmountable at physiological temperatures. (These data were excluded for the umbrella sampling analysis.) OPEN-type conformations have been observed in other MD simulations [4] [5], and the OPEN-type has been suggested as a transition state to the RISE-type in the intercalation [5]. Together, these observations suggest a dynamic intercalation model in which OPEN-type and RISE-type conformations represent interconvertible states rather than distinct endpoints, with transitions governed by local energetic and structural factors.

### Persistence length compared with experiments

Recent single-molecule force-extension experiments on intercalated DNA employ low forces (<10 pN) to maintain constant intercalator occupancy during measurements. These experiments reveal that persistence length for SYBR [10] and YOYO [12] initially decreases at low fractional elongation (defined as the end-to-end extension normalized by the contour length) around 0.1, then recovers toward free dsDNA values at higher elongations.

For SYBR, experimental measurements show that persistence length decreases at concentrations up to 0.16 µM, then increases at higher concentrations [10]. In the low-concentration regime (0–0.16 µM), the contour length of 7.9-kb DNA increases from 2.4 to 2.9 µm, corresponding to a fractional elongation of (2.9 – 2.4)/2.4 = 0.208 [10]. In our simulation of 18-mer DNA with one SYBR molecule, the fractional elongation is estimated to be 1/17 = 0.059 (where 1 represents the extension from one intercalation event and 17 is the number of base-pair steps). As this value falls within the range of decreased persistence length in our simulation, our observation of reduced persistence length is consistent with experimental trends.

For YOYO, experiments similarly show decreased persistence length at low concentrations, followed by recovery to free DNA levels at higher concentrations[11] [12]. Specifically, persistence length is minimized at a staining ratio (dye/bp) of ∼0.1 (fractional elongation ∼0.06) and recovers at a ratio of ∼0.2 (fractional elongation ∼0.35) [12]. Our simulations of 18mer-DNA yield fractional elongations of approximately 2 / 17 = 0.12 for one-YOYO and 4/17 = 0.24 for two-YOYO. These values bracket the experimental transition point, and our results, which show decreased persistence length for one-YOYO and recovery for two-YOYO, agree well with experimental data.

The non-monotonic persistence length behavior observed for SYBR (+2 charge) and YOYO (+4 charge) likely arises from competing effects. At low intercalator concentrations, local structural disruption at intercalation sites introduces flexibility, decreasing persistence length. At high concentrations, electrostatic repulsion between cationic intercalators distributed along the DNA produces a self-stiffening effect, mechanically restraining bending fluctuations and recovering rigidity. However, the observation that the persistence length plateaus beyond a staining ratio of 0.4 despite stronger repulsion suggests that additional factors may operate at high concentrations, such as changes in binding mode or saturation effects that limit further intercalation.

Unlike SYBR and YOYO, DOXO (+1 charge) shows monotonically increasing persistence length until fractional elongation of ∼0.23 in optical tweezer experiments [8]. Previous MD simulations of DNA–daunomycin [17] and DNA–DOXO [19] complexes show that the second intercalation is more favorable than the first (binding free energy for two molecules is less than double that for one). This cooperativity promotes binding of multiple intercalators to DNA, resulting in uniform distribution along the helix. This accumulation progressively straightens the DNA helix and increases its persistence length. Testing this hypothesis would require extended simulations with longer DNA and multiple DOXO molecules to directly observe cooperative binding effects on global DNA structure.

### Role of linker structure in bis-intercalators

It is helpful to understand YOYO’s mechanical behavior by comparing it with other bis-intercalators with different linker structures. Ditercalinium, another bis-intercalator with +4 charge, exhibits monotonically increasing persistence length [63], contrasting with YOYO’s plateau at high concentrations. The key structural difference lies in their linkers: YOYO possesses a long, highly flexible alkyl linker with quaternary amines and multiple rotatable bonds, whereas ditercalinium has a rigid linker comprising two piperidine rings [63]. The rigid linker of ditercalinium, combined with its highly positive charge, likely prevents the conformational flexibility that would allow DNA bending, resulting in monotonic stiffening with increasing concentration.

Thiocoraline, an uncharged bis-intercalator, provides further evidence for the importance of linker rigidity: it increases stretch modulus while maintaining constant persistence length at high concentrations [64]. Since thiocoraline is uncharged, its mechanical effects cannot arise from electrostatic repulsion. Like ditercalinium, thiocoraline possesses a rigid linker—a large macrocycle with a disulfide bridge. This rigid linker likely constrains DNA conformational freedom, increasing axial stiffness (stretch modulus) while preventing the bending flexibility changes observed with YOYO’s flexible linker.

These comparisons demonstrate that linker structure is a critical determinant of bis-intercalator effects on DNA mechanics. Our simulations directly support this conclusion: we observed YOYO bis-intercalation with variable spacing between moieties—either one or three base pairs separating the two intercalation sites (Figure S13). This variability demonstrates that YOYO’s flexible linker can accommodate multiple binding configurations, enabling DNA bending motions that rigid linkers such as those in ditercalinium and thiocoraline would prohibit. Thus, YOYO’s flexible linker enables the non-monotonic persistence length behavior we observed: at low concentrations, local disruption dominates; at high concentrations, electrostatic repulsion provides some stiffening, but the flexible linker cannot enforce the rigid DNA conformations imposed by ditercalinium or thiocoraline. This explains why YOYO’s persistence length plateaus rather than continuing to increase at high concentrations.

### Additional factors affecting DNA mechanical properties

We examined the stretch modulus and persistence length of DNA. We then considered factors such as base stacking, electrostatic interactions, and linker properties, which are thought to influence persistence length or stretch modulus. However, several additional factors should be considered for a complete understanding of intercalated DNA mechanics.

Our simulations employed physiological salt concentrations (150 mM NaCl), but experimental conditions often vary. Ionic strength has complex, opposing effects on DNA mechanics: increasing ionic strength increases stretch modulus while decreasing persistence length [56]. MD simulations of free dsDNA reproduce these trends [46], suggesting that intercalated DNA likely shows a similar ionic-strength dependence. The balance between local disruption and electrostatic stiffening effects observed for YOYO and SYBR may shift significantly at different salt concentrations.

The binding mode may change with intercalator concentration. The plateau in persistence length observed for YOYO [11] [12] and SYBR [10] at high concentrations may reflect changes in binding mode rather than simple saturation. For YOYO, proposals include a shift to major groove binding at high concentrations [65] or preferential mono-intercalation over bis-intercalation [66]. Similarly, DOXO may adopt alternative binding modes at high concentrations [26].

Our identification of OPEN-type conformations adds another layer of complexity: OPEN-type intercalation disrupts base pairing without substantial DNA extension, potentially contributing to the mechanical properties observed at high intercalator concentrations. Future experimental interpretation should consider the possibility of mixed populations, including RISE-type, OPEN-type, and groove-bound states, each contributing differently to overall DNA mechanics.

## Conclusion

The elastic properties of DNA, such as stretch modulus and persistence length, play crucial roles in biological processes including DNA packaging, replication, and transcription. Understanding how intercalators, including anticancer drugs, modulate these mechanical properties is essential for rational drug design and interpreting single-molecule experiments.

Our MM-PBSA/GBSA analysis demonstrates that van der Waals (stacking) interactions drive intercalation. We identified two distinct modes: RISE-type, which extends the DNA contour length (∼3.4 Å per intercalator), and OPEN-type, which disrupts base pairing without extending the DNA contour length. For DOXO and SYBR, minor groove binding is preferred over major groove, with the stability order: RISE-type (minor groove) > RISE-type (major groove) ≈ OPEN-type. Our kinetic predictions (mono-YO > SYBR > DOXO > bis-YOYO) match experimental trends, validating the ABMD approach.

RISE-type intercalation increased DNA helical rise compared with free dsDNA, while OPEN-type intercalation showed essentially no change in rise relative to free DNA. Both intercalation modes induced DNA unwinding, with OPEN-type causing substantially greater unwinding than RISE-type despite minimal helical extension. This reflects the distinct mechanisms of OPEN-type: disruption of base pairing and stacking without increasing helical rise. Our calculated unwinding angles for RISE-type followed the order DOXO < SYBR < YOYO, in good agreement with experimental measurements.

Intercalation affects persistence length through competing mechanisms: local structural disruption decreases stiffness at low concentrations, while electrostatic repulsion between cationic intercalators increases stiffness at high concentrations. This balance explains the non-monotonic concentration dependence observed for SYBR (+2) and YOYO (+4). For YOYO, we observed non-cooperative binding with the second intercalation electrostatically disfavored, unlike smaller intercalators. YOYO’s flexible linker enables conformational adaptability, unlike the rigid linkers of ditercalinium and thiocoraline.

All three intercalators exhibit moderate, context-dependent sequence preferences. DOXO shows preferences based on base-pair composition, YOYO favors low-Twist steps, while SYBR displays the broadest sequence tolerance. These preferences align well with their applications: DOXO as a targeted chemotherapeutic agent, YOYO as a fluorescent visualization tool, and SYBR as a general nucleic acid stain.

This study provides a foundation for understanding DNA-intercalator mechanics by identifying distinct binding modes, elucidating the effects of linker flexibility, and demonstrating non-cooperative binding in highly charged systems—insights essential for future drug design and nanotechnology applications.

## Data Availability

Relevant conformations and trajectory files have been submitted to the Biological Structure Model Archive (BSM-Arc) under BSM-ID BSM00093 (https://bsma.pdbj.org/entry/93).

## Author Contribution

H.I. and H.K. designed the research. H.I. carried out all simulations, analyzed the data, and wrote the article. H.K. assisted in analyzing the data and writing the article.

## Acknowledgements

We would like to thank Yvonne Ishida for reading this manuscript carefully. This work was supported by the HPCI system provided by Hokkaido University (Project ID: hp220012 and hp230008) and JSPS KAKENHI (JP23K05726 to HI)

## REFERENCES

[1] Gago F. Stacking interactions and intercalative DNA binding. Methods (San Diego, Calif). 1998;14:277–92.

[2] Mattioli R, Ilari A, Colotti B, Mosca L, Fazi F, Colotti G. Doxorubicin and other anthracyclines in cancers: Activity, chemoresistance and its overcoming. Mol Aspects Med. 2023;93:101205.

[3] van der Zanden SY, Qiao X, Neefjes J. New insights into the activities and toxicities of the old anticancer drug doxorubicin. Febs j. 2021;288:6095–111.

[4] Galindo-Murillo R, García-Ramos JC, Ruiz-Azuara L, Cheatham TE, 3rd, Cortés-Guzmán F. Intercalation processes of copper complexes in DNA. Nucleic acids research. 2015;43:5364–76.

[5] Lei H, Wang X, Wu C. Early stage intercalation of doxorubicin to DNA fragments observed in molecular dynamics binding simulations. J Mol Graph Model. 2012;38:279–89.

[6] Cordier C, Pierre VC, Barton JK. Insertion of a bulky rhodium complex into a DNA cytosine-cytosine mismatch: an NMR solution study. Journal of the American Chemical Society. 2007;129:12287–95.

[7] Salerno D, Brogioli D, Cassina V, Turchi D, Beretta GL, Seruggia D, et al. Magnetic tweezers measurements of the nanomechanical properties of DNA in the presence of drugs. Nucleic acids research. 2010;38:7089–99.

[8] Silva EF, Bazoni RF, Ramos EB, Rocha MS. DNA-doxorubicin interaction: New insights and peculiarities. Biopolymers. 2017;107.

[9] Biebricher AS, Heller I, Roijmans RF, Hoekstra TP, Peterman EJ, Wuite GJ. The impact of DNA intercalators on DNA and DNA-processing enzymes elucidated through force-dependent binding kinetics. Nat Commun. 2015;6:7304.

[10] Kolbeck PJ, Vanderlinden W, Gemmecker G, Gebhardt C, Lehmann M, Lak A, et al. Molecular structure, DNA binding mode, photophysical properties and recommendations for use of SYBR Gold. Nucleic acids research. 2021;49:5143–58.

[11] Kundukad B, Yan J, Doyle PS. Effect of YOYO-1 on the mechanical properties of DNA. Soft Matter. 2014;10:9721–8.

[12] Günther K, Mertig M, Seidel R. Mechanical and structural properties of YOYO-1 complexed DNA. Nucleic acids research. 2010;38:6526–32.

[13] Galindo-Murillo R, Cheatham TE. Ethidium bromide interactions with DNA: an exploration of a classic DNA-ligand complex with unbiased molecular dynamics simulations. Nucleic acids research. 2021;49:3735–47.

[14] Sahoo AK, Bagchi B, Maiti PK. Understanding enhanced mechanical stability of DNA in the presence of intercalated anticancer drug: Implications for DNA associated processes. The Journal of Chemical Physics. 2019;151.

[15] Mukherjee A, Lavery R, Bagchi B, Hynes JT. On the molecular mechanism of drug intercalation into DNA: a simulation study of the intercalation pathway, free energy, and DNA structural changes. Journal of the American Chemical Society. 2008;130:9747–55.

[16] Wilhelm M, Mukherjee A, Bouvier B, Zakrzewska K, Hynes JT, Lavery R. Multistep drug intercalation: molecular dynamics and free energy studies of the binding of daunomycin to DNA. Journal of the American Chemical Society. 2012;134:8588–96.

[17] Trieb M, Rauch C, Wibowo FR, Wellenzohn B, Liedl KR. Cooperative effects on the formation of intercalation sites. Nucleic acids research. 2004;32:4696–703.

[18] Jawad B, Poudel L, Podgornik R, Ching WY. Thermodynamic Dissection of the Intercalation Binding Process of Doxorubicin to dsDNA with Implications of Ionic and Solvent Effects. The journal of physical chemistry B. 2020;124:7803–18.

[19] Jawad B, Poudel L, Podgornik R, Steinmetz NF, Ching WY. Molecular mechanism and binding free energy of doxorubicin intercalation in DNA. Phys Chem Chem Phys. 2019;21:3877–93.

[20] Wang J, Wang W, Kollman PA, Case DA. Automatic atom type and bond type perception in molecular mechanical calculations. J Mol Graph Model. 2006;25:247–60.

[21] Wang J, Wolf RM, Caldwell JW, Kollman PA, Case DA. Development and testing of a general amber force field. Journal of computational chemistry. 2004;25:1157–74.

[22] Frisch MJ, Trucks GW, Schlegel HB, Scuseria GE, Robb MA, Cheeseman JR, et al. Gaussian 16 Rev. C.01. Wallingford, CT 2016.

[23] Bayly CI, Cieplak P, Cornell W, Kollman PA. A well-behaved electrostatic potential based method using charge restraints for deriving atomic charges: the RESP model. The Journal of Physical Chemistry. 1993;97:10269–80.

[24] Case DA, Aktulga HM, Belfon K, Cerutti DS, Cisneros GA, Cruzeiro VWD, et al. AmberTools. Journal of Chemical Information and Modeling. 2023;63:6183–91.

[25] Spielmann HP, Wemmer DE, Jacobsen JP. Solution structure of a DNA complex with the fluorescent bis-intercalator TOTO determined by NMR spectroscopy. Biochemistry. 1995;34:8542–53.

[26] Pérez-Arnaiz C, Busto N, Leal JM, García B. New insights into the mechanism of the DNA/doxorubicin interaction. The journal of physical chemistry B. 2014;118:1288–95.

[27] Jung J, Mori T, Kobayashi C, Matsunaga Y, Yoda T, Feig M, et al. GENESIS: a hybrid-parallel and multi-scale molecular dynamics simulator with enhanced sampling algorithms for biomolecular and cellular simulations. WIREs Computational Molecular Science. 2015;5:310–23.

[28] Kobayashi C, Jung J, Matsunaga Y, Mori T, Ando T, Tamura K, et al. GENESIS 1.1: A hybrid-parallel molecular dynamics simulator with enhanced sampling algorithms on multiple computational platforms. Journal of computational chemistry. 2017;38:2193–206.

[29] Ivani I, Dans PD, Noy A, Pérez A, Faustino I, Hospital A, et al. Parmbsc1: a refined force field for DNA simulations. Nat Methods. 2016;13:55–8.

[30] Joung IS, Cheatham TE. Determination of alkali and halide monovalent ion parameters for use in explicitly solvated biomolecular simulations. J Phys Chem B. 2008;112:9020–41.

[31] Jorgensen WL, Chandrasekhar J, Madura JD, Impey RW, Klein ML. Comparison of simple potential functions for simulating liquid water. J Chem Phys. 1983;79:926–35.

[32] Darden T, York D, Pedersen L. Particle mesh Ewald: An N°log(N) method for Ewald sums in large systems. The Journal of Chemical Physics. 1993;98:10089–92.

[33] Essmann U, Perera L, Berkowitz ML, Darden T, Lee H, Pedersen LG. A smooth particle mesh Ewald method. The Journal of Chemical Physics. 1995;103:8577–93.

[34] Ryckaert J-P, Ciccotti G, Berendsen H. Numerical-Integration of Cartesian Equations of Motion of a System with Constraints – Molecular-Dynamics of N-Alkanes. Journal of Computational Physics. 1977;23:327–41.

[35] Gunsteren WFv, Berendsen HJC. Algorithms for macromolecular dynamics and constraint dynamics. Molecular Physics. 1977;34:1311--27.

[36] Babin V, Roland C, Sagui C. Adaptively biased molecular dynamics of free energy calculations. J Chem Phys. 2008;128:134101.

[37] Ishida H, Kono H. Free Energy Landscape of H2A-H2B Displacement From Nucleosome. Journal of molecular biology. 2022;434:167707.

[38] Kumar S, Bouzida D, Swendsen RH, Kollman PA, Rosenberg JM. The weighted histogram analysis method for free energy calculations on biomolecules. I. The method. J Comput Chem. 1992;13:1011–21.

[39] Schlitter J. Estimation of Absolute and Relative Entropies of Macromolecules Using the Covariance Matrix. Chem Phys Lett. 1993;215:617–21.

[40] Ishida H. Essential function of the N-termini tails of the proteasome for the gating mechanism revealed by molecular dynamics simulations. Proteins. 2014;82:1985–99.

[41] Ishida H, Kono H. Torsional stress can regulate the unwrapping of two outer half superhelical turns of nucleosomal DNA. Proceedings of the National Academy of Sciences of the United States of America. 2021;118.

[42] Lu XJ, Olson WK. 3DNA: a software package for the analysis, rebuilding and visualization of three-dimensional nucleic acid structures. Nucleic acids research. 2003;31:5108–21.

[43] Rizzo V, Sacchi N, Menozzi M. Kinetic studies of anthracycline-DNA interaction by fluorescence stopped flow confirm a complex association mechanism. Biochemistry. 1989;28:274–82.

[44] Odijk T. Stiff Chains and Filaments under Tension. Macromolecules. 1995;28:7016–8.

[45] Mogurampelly S, Nandy B, Netz RR, Maiti PK. Elasticity of DNA and the effect of dendrimer binding. Eur Phys J E Soft Matter. 2013;36:68.

[46] Garai A, Saurabh S, Lansac Y, Maiti PK. DNA Elasticity from Short DNA to Nucleosomal DNA. The journal of physical chemistry B. 2015;119:11146–56.

[47] Qiu D, Shenkin PS, Hollinger FP, Still WC. The GB/SA Continuum Model for Solvation. A Fast Analytical Method for the Calculation of Approximate Born Radii. The Journal of Physical Chemistry A. 1997;101:3005–14.

[48] Miller BR, 3rd, McGee TD, Jr., Swails JM, Homeyer N, Gohlke H, Roitberg AE. MMPBSA.py: An Efficient Program for End-State Free Energy Calculations. Journal of chemical theory and computation. 2012;8:3314–21.

[49] Su PC, Tsai CC, Mehboob S, Hevener KE, Johnson ME. Comparison of radii sets, entropy, QM methods, and sampling on MM-PBSA, MM-GBSA, and QM/MM-GBSA ligand binding energies of F. tularensis enoyl-ACP reductase (FabI). Journal of computational chemistry. 2015;36:1859–73.

[50] Sanner MF, Olson AJ, Spehner JC. Reduced surface: an efficient way to compute molecular surfaces. Biopolymers. 1996;38:305–20.

[51] Still WC, Tempczyk A, Hawley RC, Hendrickson T. Semianalytical treatment of solvation for molecular mechanics and dynamics. J Am Chem Soc. 1990;112:6127–9.

[52] Ishida H, Matsumoto A. Mechanism for verification of mismatched and homoduplex DNAs by nucleotides-bound MutS analyzed by molecular dynamics simulations. Proteins. 2016;84:1287–303.

[53] Nozaki T, Onoda M, Habazaki M, Takeuchi Y, Ishida H, Sato Y, et al. Designer Catalyst-Enabled Regiodivergent Histone Acetylation. Journal of the American Chemical Society. 2025;147:13732–43.

[54] van Mameren J, Vermeulen K, Wuite GJL, Peterman EJG. A polarized view on DNA under tension. J Chem Phys. 2018;148:123306.

[55] Zeng Y, Montrichok A, Zocchi G. Bubble nucleation and cooperativity in DNA melting. Journal of molecular biology. 2004;339:67–75.

[56] Baumann CG, Smith SB, Bloomfield VA, Bustamante C. Ionic effects on the elasticity of single DNA molecules. Proceedings of the National Academy of Sciences of the United States of America. 1997;94:6185–90.

[57] Cullinane C, Phillips DR. Induction of stable transcriptional blockage sites by adriamycin: GpC specificity of apparent adriamycin-DNA adducts and dependence on iron(III) ions. Biochemistry. 1990;29:5638–46.

[58] DuVernay VH, Jr., Pachter JA, Crooke ST. Deoxyribonucleic acid binding studies on several new anthracycline antitumor antibiotics. Sequence preference and structure--activity relationships of marcellomycin and its analogues as compared to adriamycin. Biochemistry. 1979;18:4024–30.

[59] Airoldi M, Barone G, Gennaro G, Giuliani AM, Giustini M. Interaction of doxorubicin with polynucleotides. A spectroscopic study. Biochemistry. 2014;53:2197–207.

[60] Frederick CA, Williams LD, Ughetto G, van der Marel GA, van Boom JH, Rich A, et al. Structural comparison of anticancer drug-DNA complexes: adriamycin and daunomycin. Biochemistry. 1990;29:2538–49.

[61] Kellogg GE, Scarsdale JN, Fornari FA, Jr. Identification and hydropathic characterization of structural features affecting sequence specificity for doxorubicin intercalation into DNA double-stranded polynucleotides. Nucleic acids research. 1998;26:4721–32.

[62] Reuter M, Dryden DT. The kinetics of YOYO-1 intercalation into single molecules of double-stranded DNA. Biochem Biophys Res Commun. 2010;403:225–9.

[63] Berge T, Jenkins NS, Hopkirk RB, Waring MJ, Edwardson JM, Henderson RM. Structural perturbations in DNA caused by bis-intercalation of ditercalinium visualised by atomic force microscopy. Nucleic acids research. 2002;30:2980–6.

[64] Camunas-Soler J, Manosas M, Frutos S, Tulla-Puche J, Albericio F, Ritort F. Single-molecule kinetics and footprinting of DNA bis-intercalation: the paradigmatic case of Thiocoraline. Nucleic acids research. 2015;43:2767–79.

[65] Larsson A, Carlsson C, Jonsson M, Albinsson B. Characterization of the Binding of the Fluorescent Dyes YO and YOYO to DNA by Polarized Light Spectroscopy. Journal of the American Chemical Society. 1994;116:8459–65.

[66] Kucharska K, Pilz M, Bielec K, Kalwarczyk T, Kuzma P, Holyst R. Two Intercalation Mechanisms of Oxazole Yellow Dimer (YOYO-1) into DNA. Molecules (Basel, Switzerland). 2021;26.

