## Supplementary Information for "Impact of intercalators on the properties of DNA analyzed by molecular dynamics simulations"

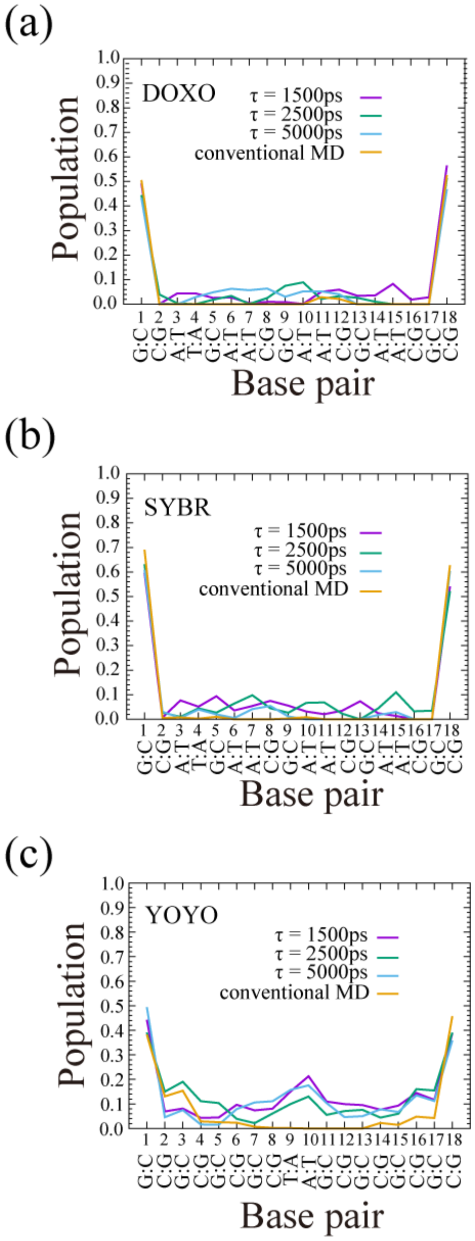

Figure S1: Distribution of (a) DOXO, (b) SYBR, and (c) YOYO on the DNA

The trajectories of 24 replicas for 700 ns (24 × 700 = 16,800 data) were used to calculate the distance between the COM of the intercalative part of the intercalator and the COM of the base pair. The number of four intercalator molecules with a distance of less than 4.0 Å was counted and normalized by the number of the data (16,800), which was defined as population. Therefore, if the population was four at the 1$^{st}$ base pair, then four DOXO/SYBR molecules would always be near the 1st base pair. Similarly, if the population was eight at the 1$^{st}$ base pair, then eight YO-moieties of YOYO were always near the 1st base pair.

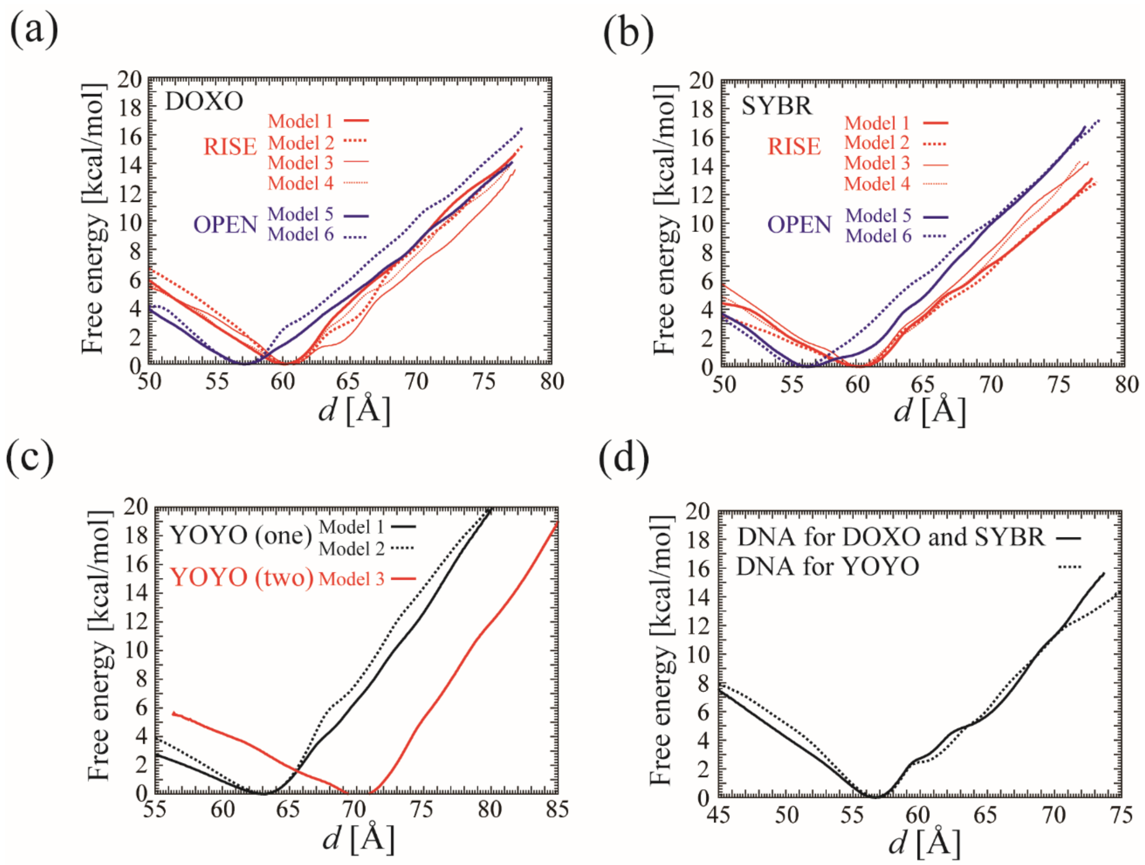

Figure S2: Free energy curve for the extension and contraction of the intercalated DNA along *d*.

The FECs of the DNA intercalation against *d* (in units of Å) for (a) DOXO, (b) SYBR, (c) YOYO, and (d) the free dsDNA, respectively. The values of the free energies were set at zero at $d = d_{\mathrm{min}}$.

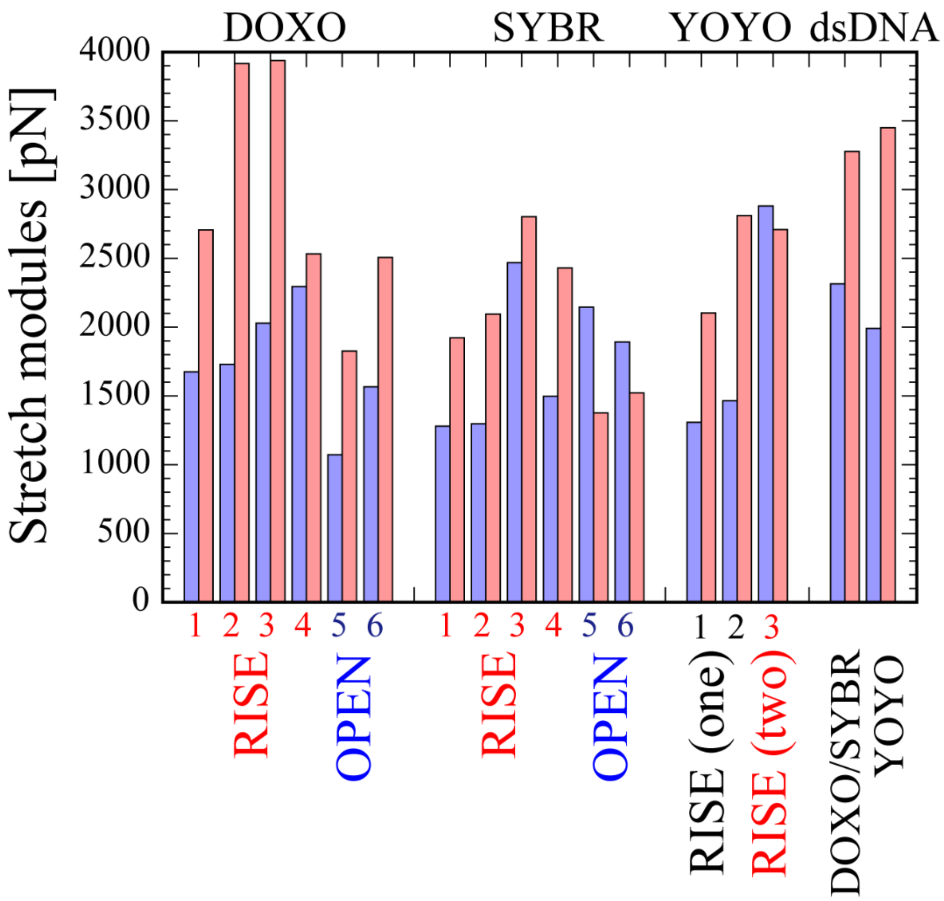

Figure S3: The stretch modulus for the intercalated DNA.

The stretch module for extension (in red) and contraction (in blue) of the intercalated DNA for DOXO, SYBR, YOYO, and the free dsDNA are shown.

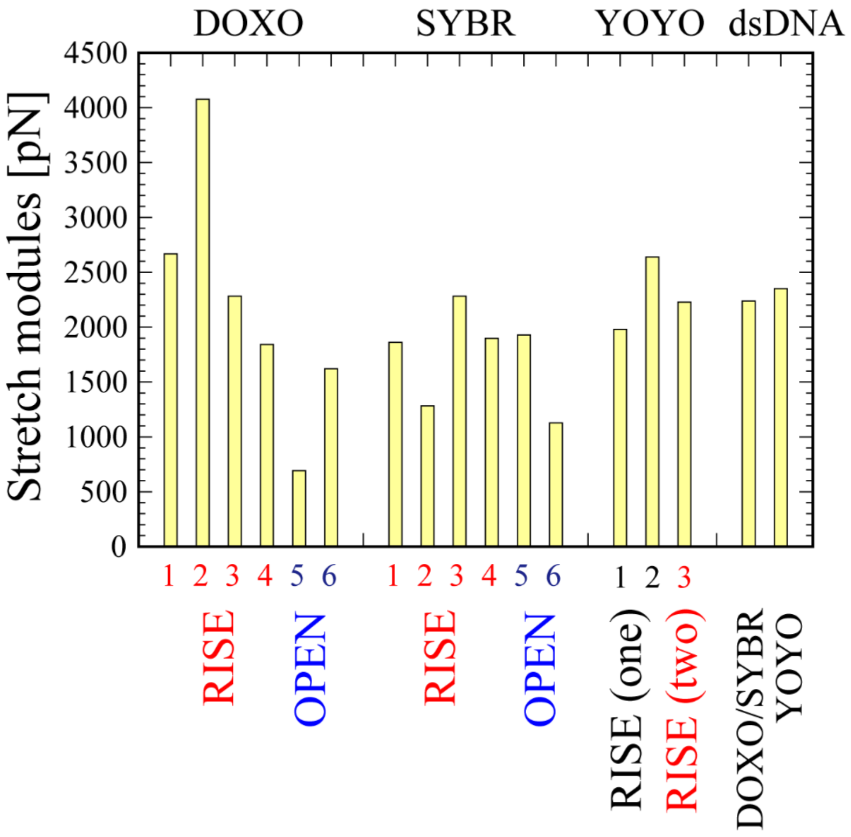

Figure S4: The stretch modules for the DNA calculated from the contour length.

For the fitting of contour length in Eq. (6), the range of the contour length which falls within the elastic region of *d* was used (see also Table SIV).

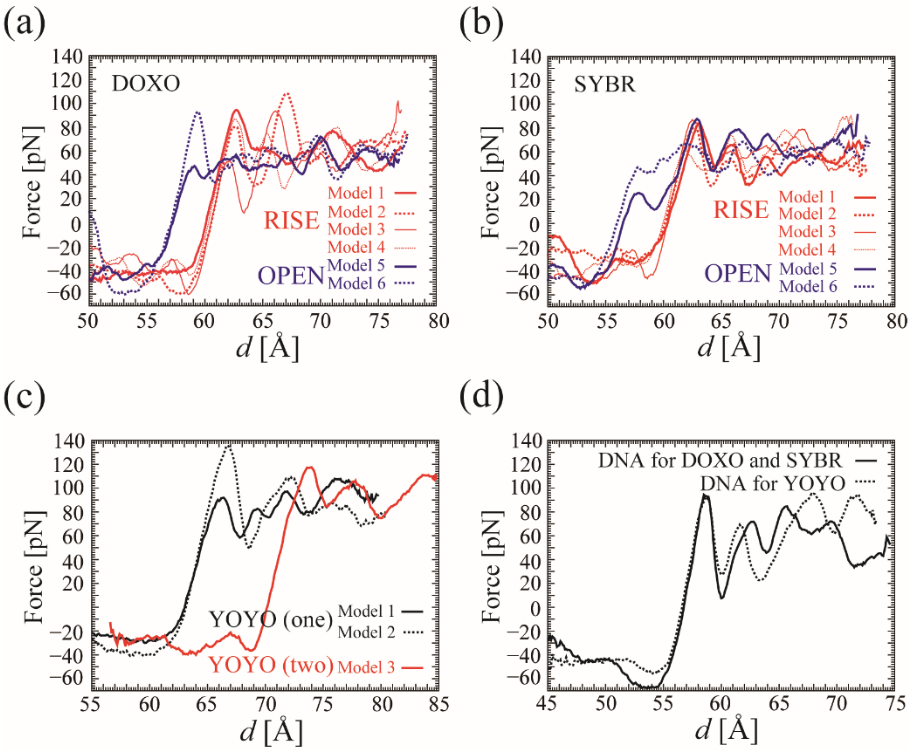

Figure S5: Force for the extension and contraction of the intercalated DNA

The force was calculated as the derivative of the free energy curve, obtained by the lest-square fitting of seven data points (the current points with three preceding and three subsequent points) at the interval of 0.1 Å.

(a) For DOXO, the forces at the first peak of modes 1-6 were 93.7 pN at $d$ = 62.8 Å, 80.4 pN at $d$ = 62.6 Å, 51.6 pN at $d$ = 62.0 Å, 85.8 pN at $d$ = 62.6 Å, 46.5 pN at $d$ = 59.0 Å, 91.6 pN at $d$ = 59.4 Å, respectively. The averaged forces after the first peak of models 1-6 were 60.4, 63.4, 58.6, 59.9, 51.3, 55.9 pN, respectively.

(b) For SYBR, the forces at the first peak of models 1-6 were 82.8 pN at $d$ = 63.1 Å, 74.3 pN at $d$ = 63.8 Å, 80.4 pN at $d$ = 62.6 Å, 86.1 pN at $d$ = 62.5 Å, 25.3 pN at $d$ = 57.8 Å, 46.2 pN at $d$ = 58.0 Å, respectively. The averaged forces after the first peak of models 1-6 were 54.7, 57.6, 51.2, 59.8, 59.2, 56.5 pN, respectively.

(c) For YOYO, the forces at the first peak of models 1-3 were 90.1 pN at $d$ = 66.0 Å, 134.5 pN at $d$ = 66.8 Å, 117.2 pN at $d$ = 73.9 Å. The averaged forces after the first peak of models 1-3 were 87.9, 84.7, and 96.4 pN, respectively.

(d) For the free dsDNA, the forces at the first peak for the free dsDNA with DNA sequence of DOXO/SYBR and YOYO were 93.2 pN at $d$ = 58.6 Å, and 91.7 pN at $d$ = 58.7 Å, respectively. The averaged forces after the first peak for the free dsDNA with DNA sequence of DOXO/SYBR and YOYO were 55.7 and 65.3 pN, respectively.

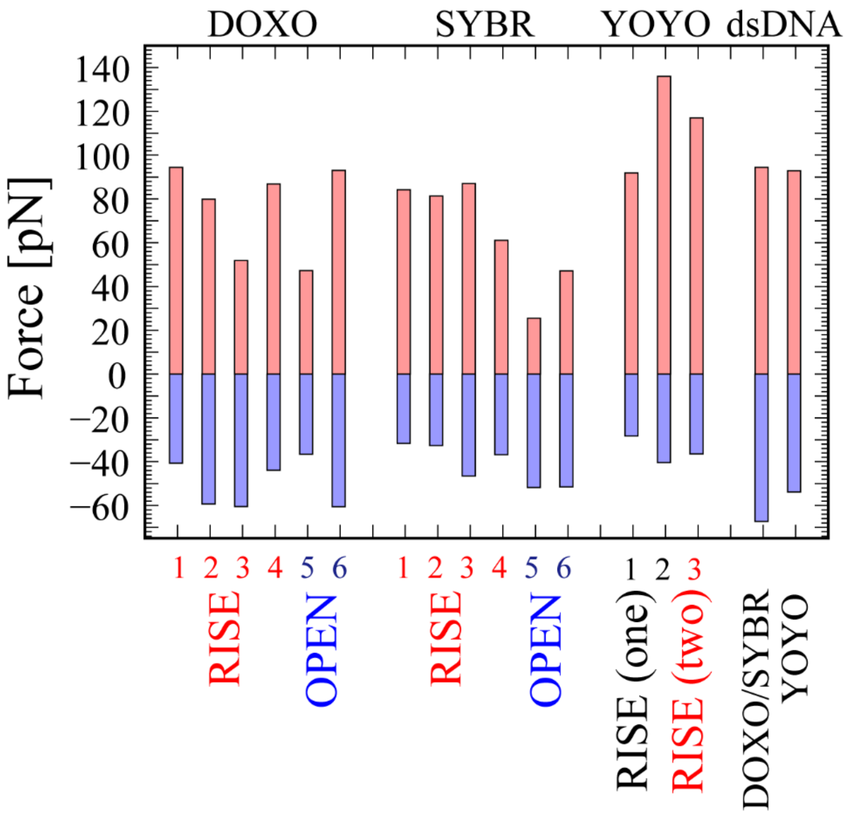

Figure S6: Force at the first peak for the extension and contraction of the intercalated DNA

The force at the first peak for the extension (in red) and contraction (in blue) of the intercalated DNA for DOXO, SYBR, and YOYO, and the free dsDNA are shown.

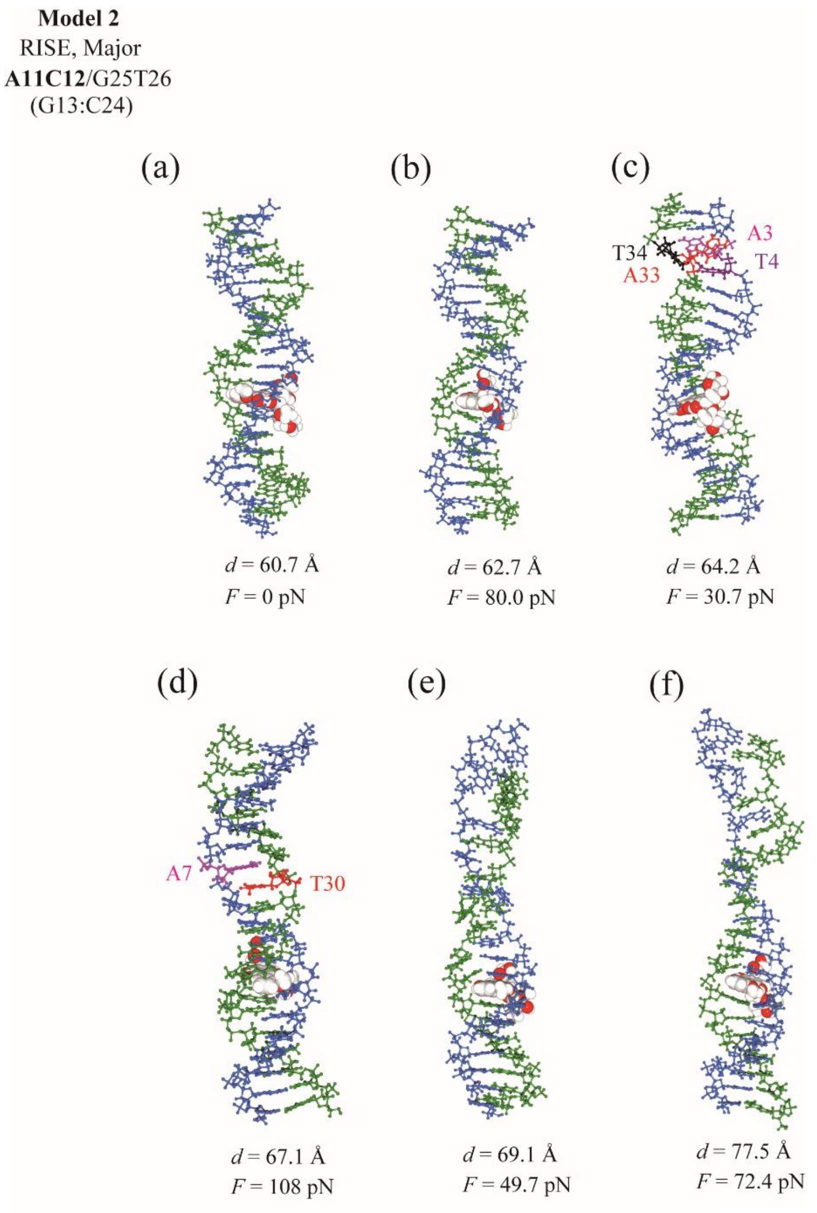

Figure S7A

Figure S7A: Overstretch of the DNA with partial melting of the DNA (for DOXO)

For DOXO model2, (a) Force = 0 pN at $d_{\mathrm{min}}$ = 60.7 Å, (b) Force = 80.0 pN at *d* = 62.7 Å, (c) Force = 30.7 pN at *d* = 64.2 Å, where base pairs A3:T34 and T4:A34 were broken. (d) Force = 108.2 pN at *d* = 67.1 Å, where A7 and T30 formed a ladder structure. (e) Force = 49.7 pN at *d* = 69.1 Å, where T4 and G5 were separated from each other. (f) Force=72.4 pN at *d* = 77.5 Å

It should be noted that the series along *d* is not a time series in the umbrella sampling simulation.

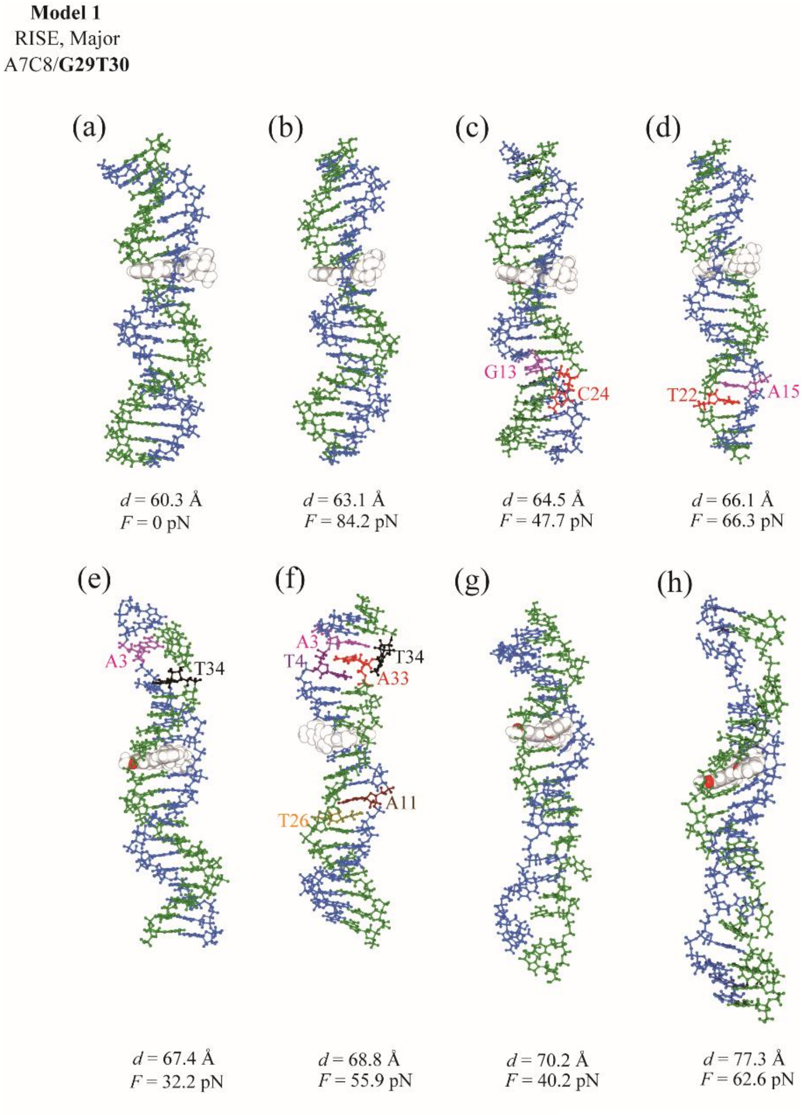

Figure S7B

Figure S7B: Overstretch of the DNA with partial melting of the DNA (for SYBR)

For SYBR model 1, (a) Force = 0 pN at $d_{\text{min}}$ = 60.3 Å, (b) Force = 84.2 pN at *d* = 63.1 Å, (c) Force = 47.7 pN at *d* = 64.5 Å, where a base pair G13:C24 was broken. (d) Force = 66.3 pN at *d* = 66.1 Å, where A15 and T22 formed a ladder structure. (e) Force = 32.2 pN at *d* = 67.4 Å, where base pairs A3:T34 and T4:A33 are broken. (f) Force = 55.9 pN at *d* = 68.8 Å, where A3, A33, and T4 formed a ladder structure, A11 and T26 formed another ladder structure, and T34 was flipped out. (g) Force = 40.2 pN at *d* = 70.2 Å, where a base pair C16:G21 was broken. (h) Force = 62.6 pN at *d* = 77.3 Å.

It should be noted that the series along *d* is not a time series in the umbrella sampling simulation.

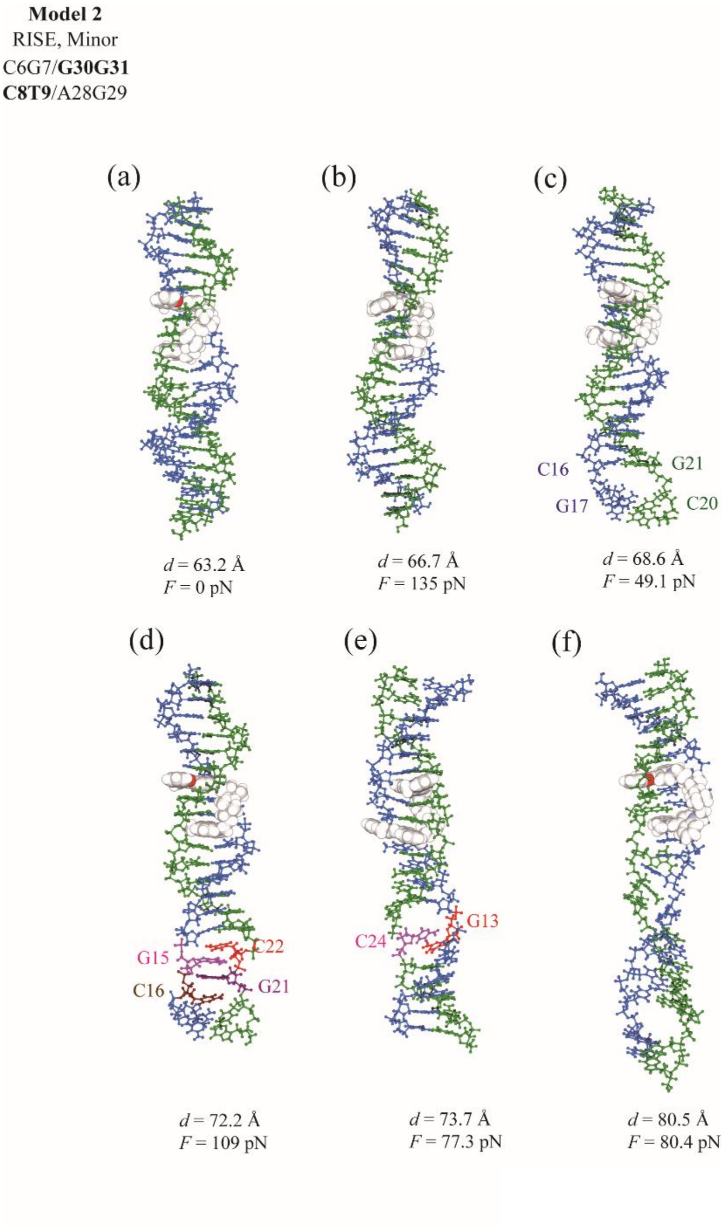

Figure S7C

Figure S7C: Overstretch of the DNA with partial melting of the DNA (for YOYO)

For YOYO model 2, (a) Force = 0 pN at $d_{\min}$ = 63.2 Å, (b) Force = 135.7 pN at $d$ =66.7 Å (c) Force = 49.1 pN at $d$ = 68.6 Å, where C16 and G17 were separated from each other. (d) Force = 109.4 pN at $d$ = 72.2 Å, where C16 and G21, G15 and C22 formed a ladder structure. (e) Force = 77.3 pN at $d$ = 73.7 Å, where C12 and G13 were separated from each other. (f) Force = 80.4 pN at $d$ = 80.5 Å

It should be noted that the series along $d$ is not a time series in the umbrella sampling simulation.

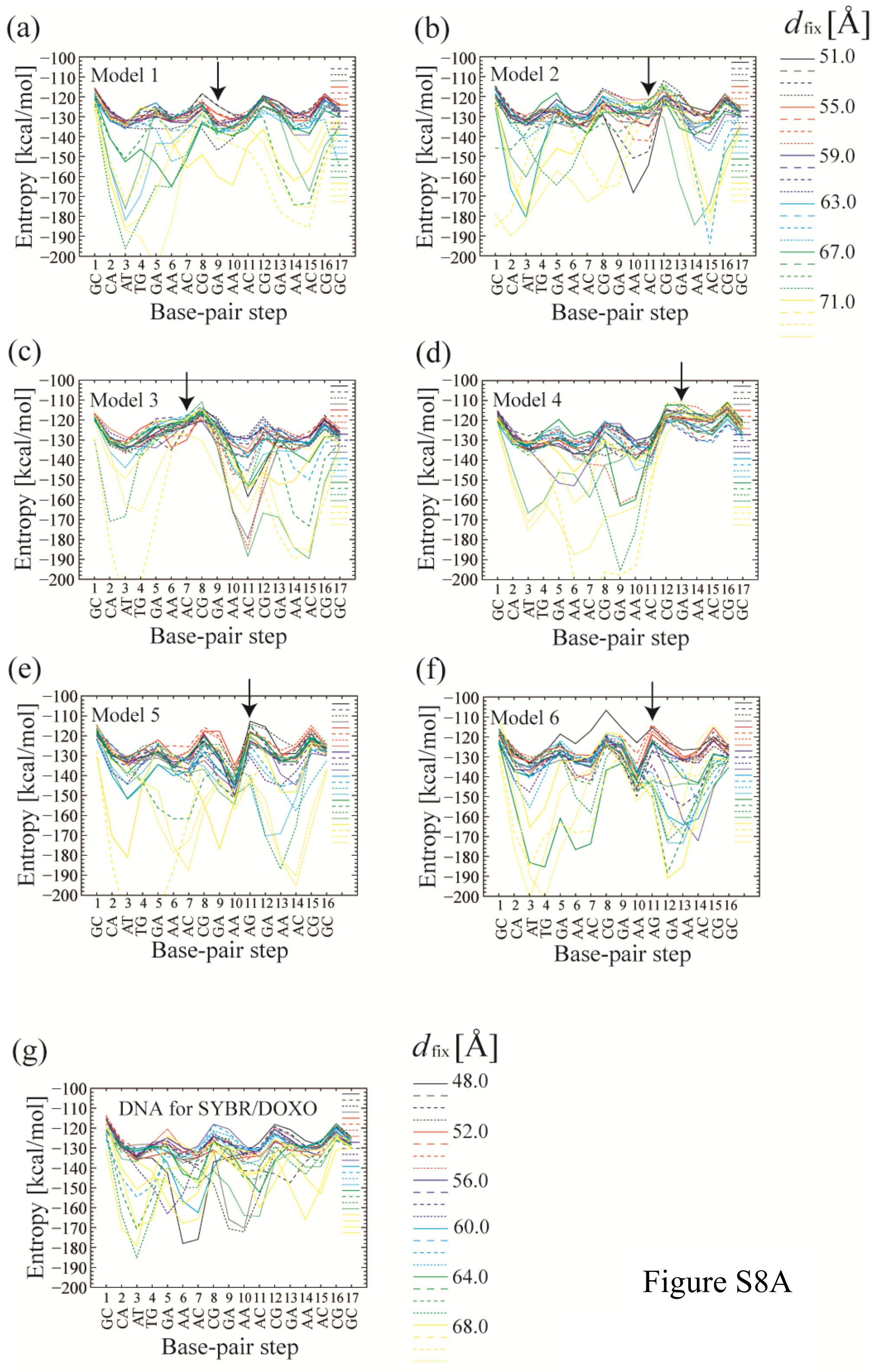
(a)
Model 1
(b)
Model 2
(c)
Model 3
(d)
Model 4
(e)
Model 5
(f)
Model 6
(g)
DNA for SYBR/DOXO
Entropy [kcal/mol]
Base-pair step
GC CA AT TG GA AA AC CG GA AA AC CG GA AA AC CG GC
GC CA AT TG GA AA AC CG GA AA AG GA AA AC CG GC
$d_{fix}$ [Å]
51.0
55.0
59.0
63.0
67.0
71.0
$d_{fix}$ [Å]
48.0
52.0
56.0
60.0
64.0
68.0

Figure S8A

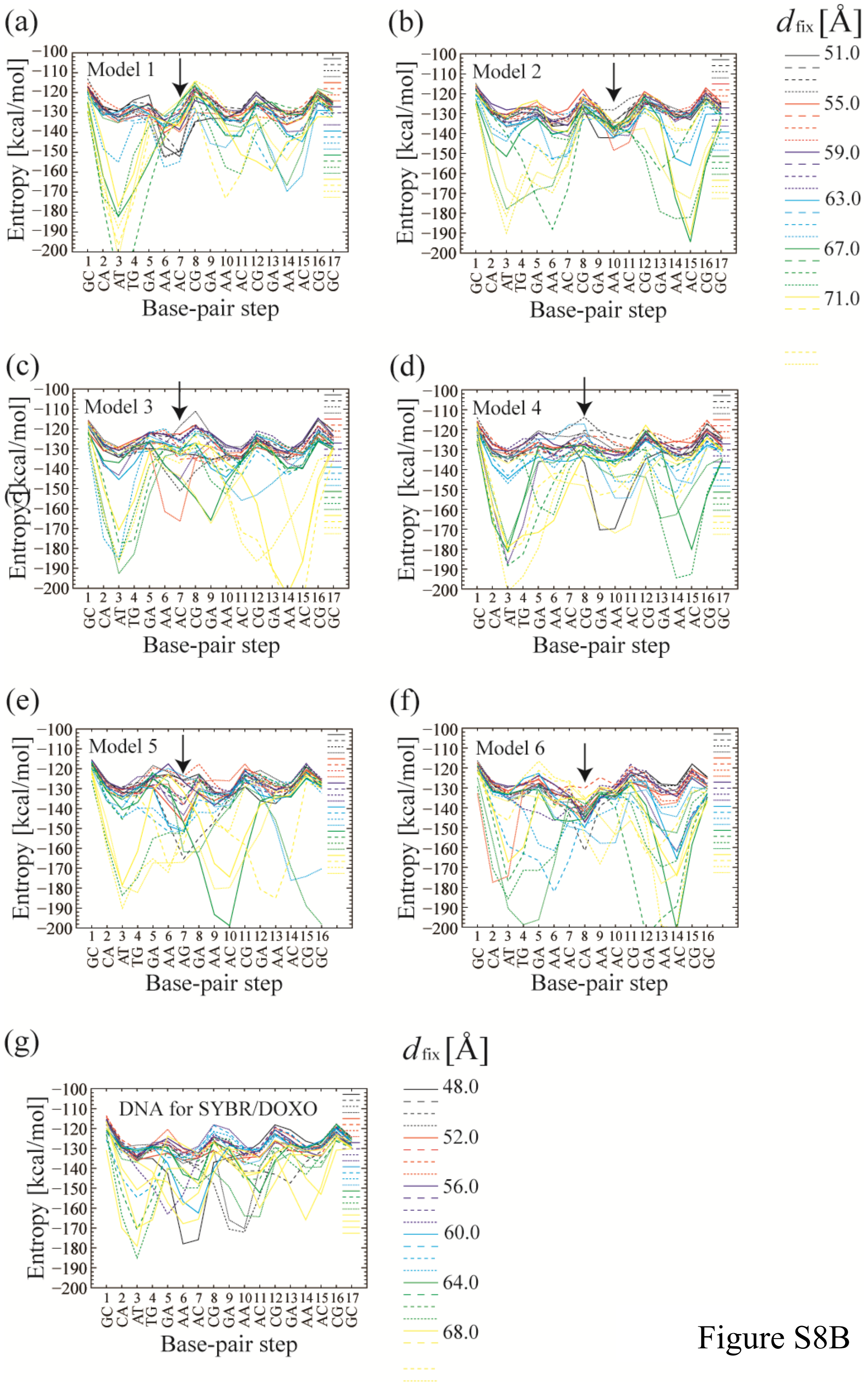

(a)
Model 1
(b)
Model 2
(c)
Model 3
(d)
Model 4
(e)
Model 5
(f)
Model 6
(g)
DNA for SYBR/DOXO
Entropy [kcal/mol]
Base-pair step
d fix [Å]
51.0
55.0
59.0
63.0
67.0
71.0
48.0
52.0
56.0
60.0
64.0
68.0

Figure S8B

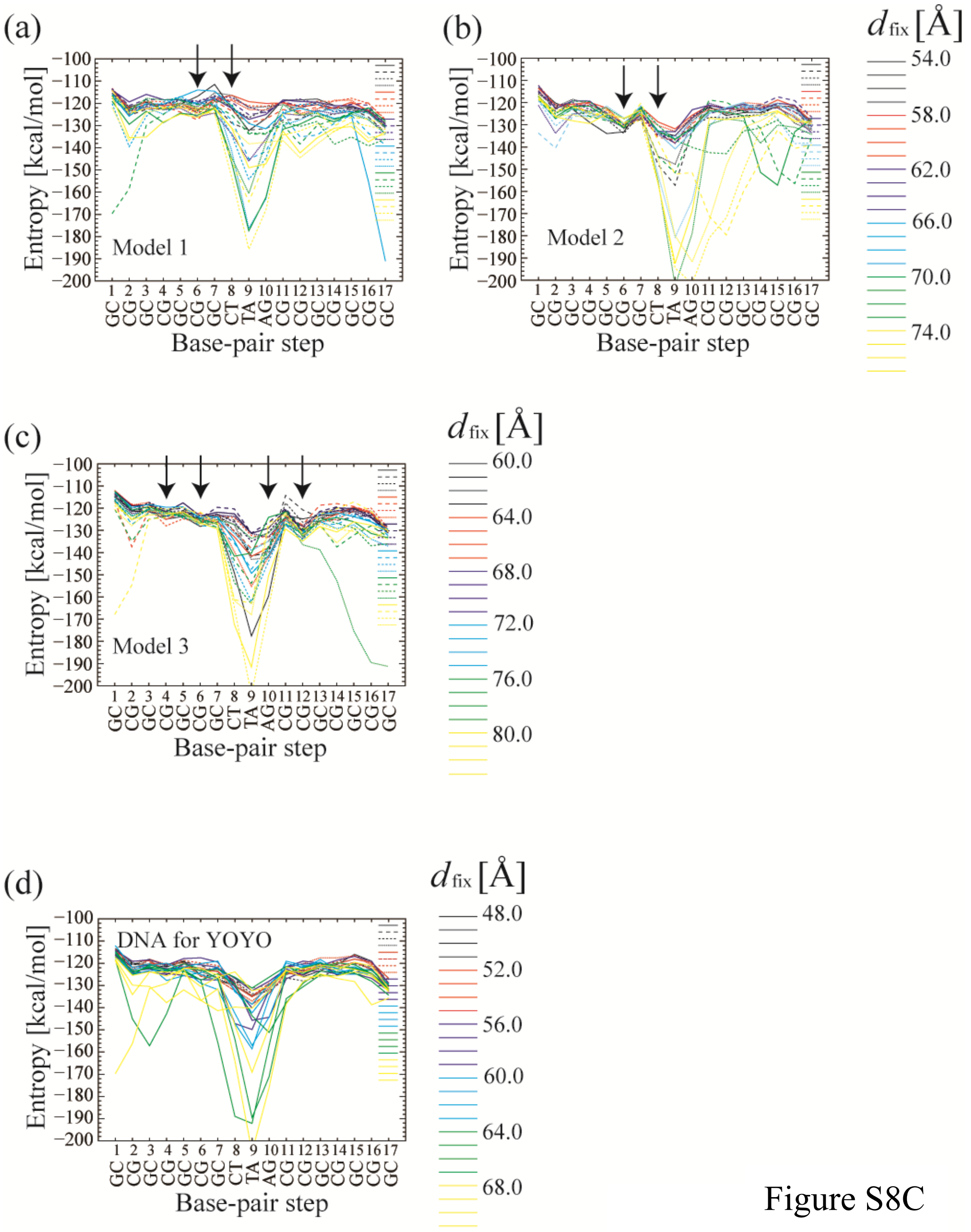

Figure S8C

Figure S8: Entropies for base-pair steps

The entropies for base-pair steps for Models 1–6 (a–f) of (A) DOXO, (B) SYBR, and (C) YOYO are shown, respectively (see Methods). The data for the window at $d_{fix}$ = 51.0 Å for DOXO Model 1, and the window at $d_{fix}$ = 61.0 Å for SYBR Model 1 were excluded for the analysis because these states changed from RISE-type to OPEN-type during the umbrella sampling simulations.

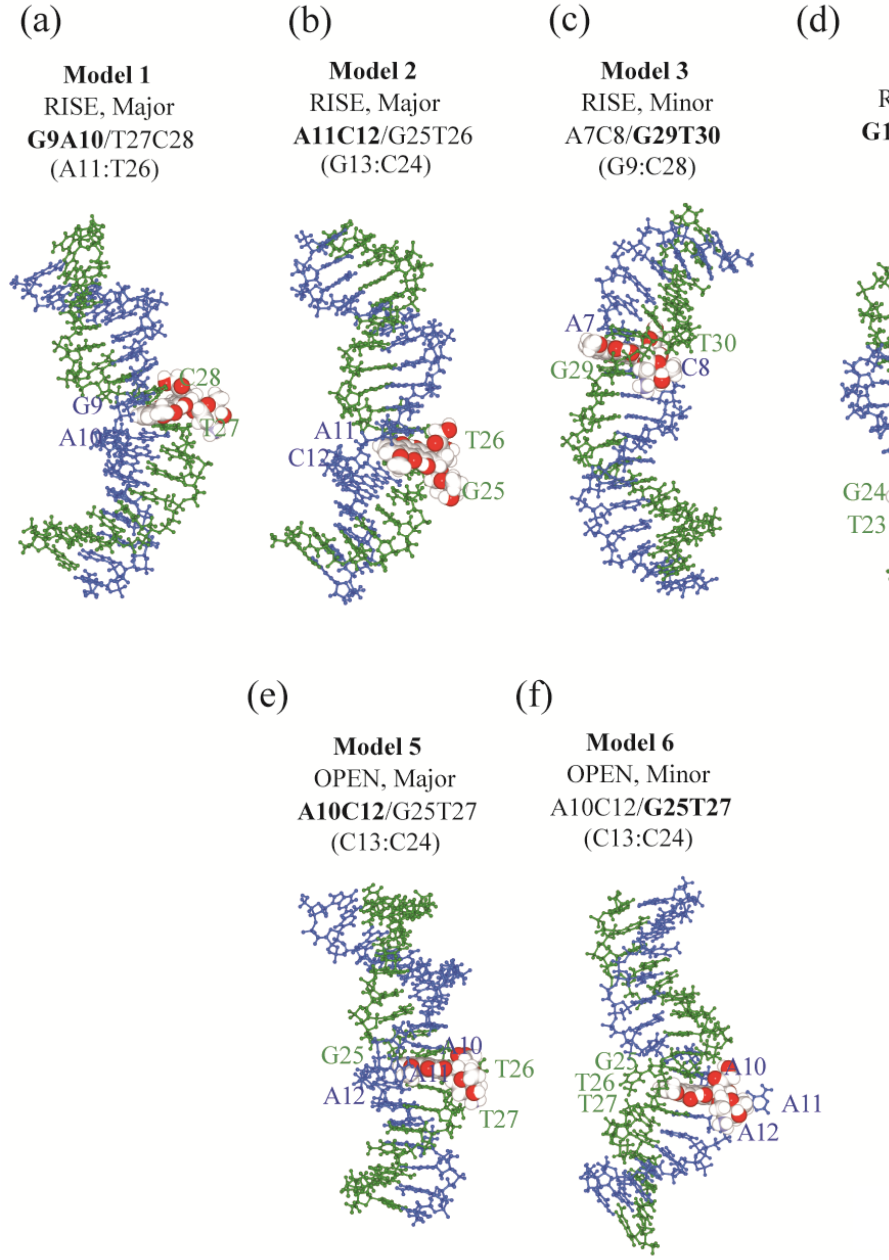

Figure S9A

Figure S9A: bent DNA at $d$ < 54.0 Å. (for DOXO)

For DOXO, snapshots for (a) model 1 at $d_{fix}$ = 52.0 Å, (b) model 2 at $d_{fix}$ = 51.0 Å, (c) model 3 at $d_{fix}$ = 52.0 Å, (d) model 4 at $d_{fix}$ = 51.0 Å, (e) model 5 at $d_{fix}$ = 51.0 Å, and (f) model 6 at $d_{fix}$ = 52.0 Å are shown.

For model 1 (RISE-type at the major groove), the DNA bent towards the opposite side of the major groove by 96.0 ± 3.7% (99.2, 89.9, 98.7, 96.0% for the window at $d_{fix}$ = 52.0, 53.0, 54.0, and 55.0 Å, respectively). For model 2 (RISE-type at the major groove), the DNA bent towards the opposite side of the major groove by 88.7 ± 8.2% (90.0, 89.3, 98.3, 73.6, 92.1% for the window for $d_{fix}$ = 51.0, 52.0, 53.0, 54.0, and 55.0 Å, respectively).

For model 3 (RISE-type at the minor groove), the DNA bent towards the opposite side of the minor groove by 39.6 ± 32.0% (98.6, 38.1, 8.5, 39.2, 13.4% for the window for $d_{fix}$ = 51.0, 52.0, 53.0, 54.0, and 55.0 Å, respectively) (i.e., the DNA bent towards the same side of the minor groove by 60.4 ± 32.0%). For model 4 (RISE-type in the minor groove), the DNA bent towards the opposite side of the minor groove by 52.4 ± 38.5% (7.5, 94.6, 100, 42.3, 17.3% for the window for $d_{fix}$ = 51.0, 52.0, 53.0, 54.0, and 55.0 Å, respectively) (i.e., the DNA bent the same side of the minor groove by 47.6 ± 38.5%). For model 5 (OPEN-type at the major groove), the DNA bent towards the opposite side of the major groove by 57.1 ± 29.9% (84.3, 2.4, 72.8, 48.8, 77.4% for the window for $d_{fix}$ = 51.0, 52.0, 53.0, 54.0, and 55.0 Å, respectively). For model 6 (OPEN-type in the minor groove) , the DNA bent towards the opposite side of the minor groove by 76.1 ± 21.6% (34.3, 93.0, 90.7, 85.4, 77.0% for the window for $d_{fix}$ = 51.0, 52.0, 53.0, 54.0, and 55.0 Å, respectively)

Consequently, RISE-type DNA with DOXO at the major groove showed that the DNA bent towards the opposite side of the intercalator by 96.0 ± 3.7% and 88.7 ± 8.2% for models 1 and 2, while RISE-type DNA with DOXO at the minor groove showed the DNA bent towards the opposite side of the intercalator by 39.6 ± 32.0% and 52.4 ± 38.5% for models 3 and 4. This indicates that the DNA tends to bend towards the opposite side for DOXO at the major groove, and towards the same side for DOXO at the minor groove. OPEN-type DNA with DOXO at the major and minor grooves bent towards the opposite side by 57.1 ± 29.9% and 76.1 ± 21.6%, showing no clear direction.

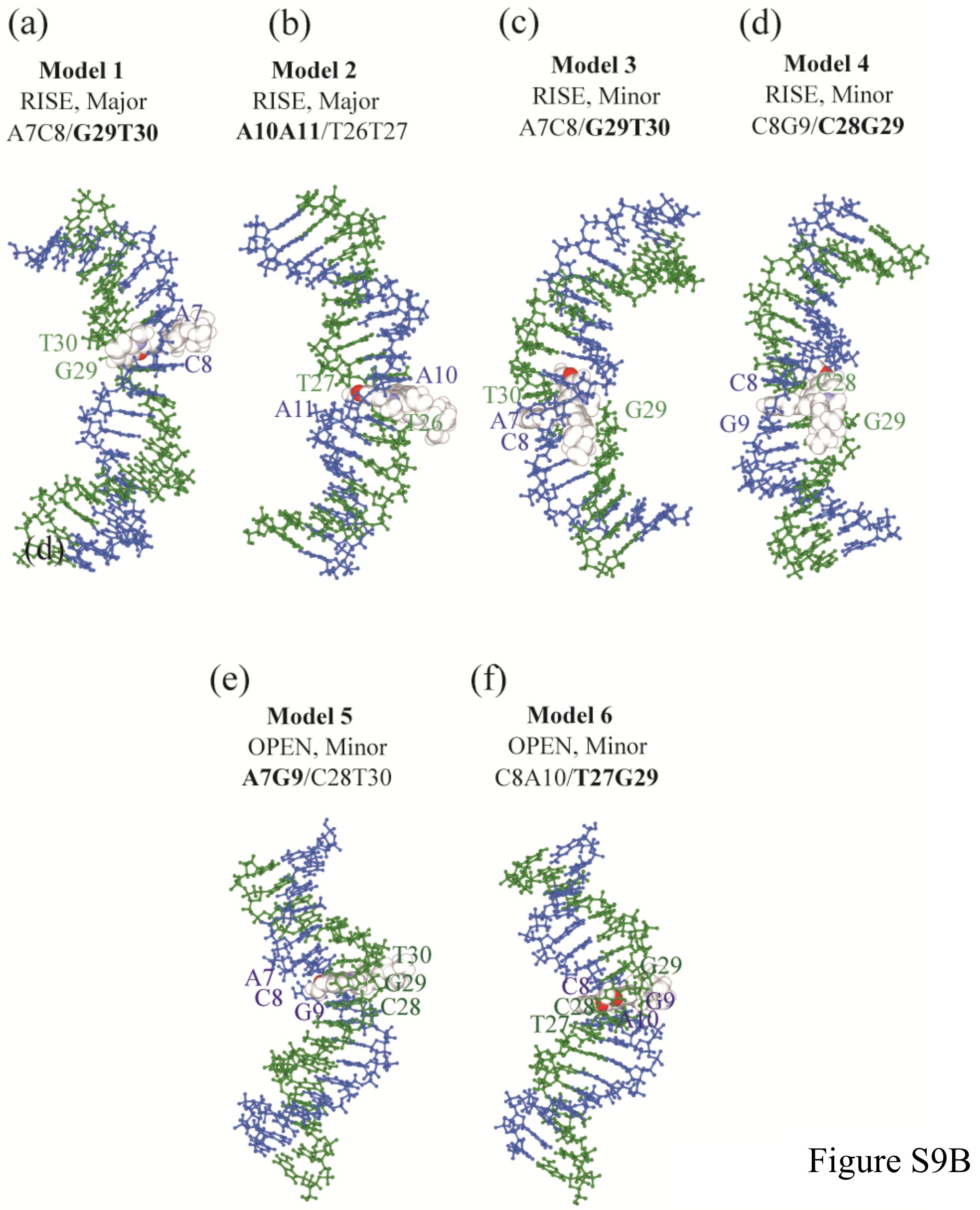

Figure S9B: bent DNA at $d$ < 54.0 Å. (for SYBR)

For SYBR, snapshots for (a) model 1 at $d_{fix}$ = 52.0 Å, (b) model 2 at $d_{fix}$ = 51.0 Å, (c) model 3 at $d_{fix}$ = 52.0 Å, (d) model 4 at $d_{fix}$ = 52.0 Å, (e) model 5 at $d_{fix}$ = 51.0 Å, and (f) model 6 at $d_{fix}$ = 51.0 Å are shown.

For model 1 (RISE-type at the major groove), the DNA bent towards the opposite side of the major groove by 68.5 ± 31.9% (10.4, 77.1, 95.8, 61.6, and 97.4% for the window at $d_{fix}$ = 51.0, 52.0, 53.0, 54.0, and 55.0 Å, respectively). For model 3 (RISE-type at the major groove), the DNA bent towards the opposite side of the major groove by

88.5 ± 16.8 % (97.5, 97.0, 96.7, 96.3, and 54.8% for the window at $d_{fix}$ = 51.0, 52.0, 53.0, 54.0, and 55.0 Å, respectively). For model 3 (RISE-type at the minor groove), the DNA bent towards the opposite side of the minor groove by 2.6 ± 1.7 % (0.8, 2.7, 5,4, 1.5, and 2.6 % for the window at $d_{fix}$ = 51.0, 52.0, 53.0, 54.0, and 55.0 Å, respectively) (i.e., the DNA bent the same side of the minor groove by 97.4 ± 1.7 %). For model 4 (RISE-type at the minor groove), DNA bent towards the opposite side of the minor groove by 10.5 ± 10.8 % (16.2, 29.0, 0.1, 4.5, and 2.7 % for the window at $d_{fix}$ = 51.0, 52.0, 53.0, 54.0, and 55.0 Å, respectively) (i.e., the DNA bent the same side of the minor groove by 88.5 ± 10.8 %). For model 5 (OPEN-type at the minor groove), the DNA bent towards the opposite side of the minor groove by 58.8 ± 33.3 % (91.4, 13.0, 83.4, 82.3, and 23.9 % for the window at $d_{fix}$ = 51.0, 52.0, 53.0, 54.0, and 55.0 Å, respectively) (i.e., the DNA bent the same side of the minor groove by 41.2 ± 33.3%). For model 6, (OPEN-type at the minor groove), DNA bent towards the opposite side of the minor groove by 74.6 ± 12.8 % (94.3, 60.2, 82.8, 74.0, and 62.0 % for the window at $d_{fix}$ = 51.0, 52.0, 53.0, 54.0, and 55.0 Å, respectively) (i.e., the DNA bent the same side of the minor groove by 25.4 ± 12.8 %)

Consequently, RISE-type DNA with SYBR at the major groove showed that the DNA bent towards the opposite side of the intercalator by 68.5 ± 31.9% and 88.5 ± 16.8 % for models 1 and 3, while RISE-type DNA with SYBR at the minor groove showed the DNA bent towards the opposite side of the intercalator by 2.6 ± 1.7 % and 10.5 ± 10.8 % for models 4 and 4B. This indicates that the DNA tends to bend towards the opposite side for SYBR at the major groove, and towards the same side for SYBR at the minor groove. OPEN-type DNA with SYBR at the major and minor grooves bent towards the opposite side by 58.8 ± 33.3 % and 74.68 ± 12.8 %, showing no clear direction.

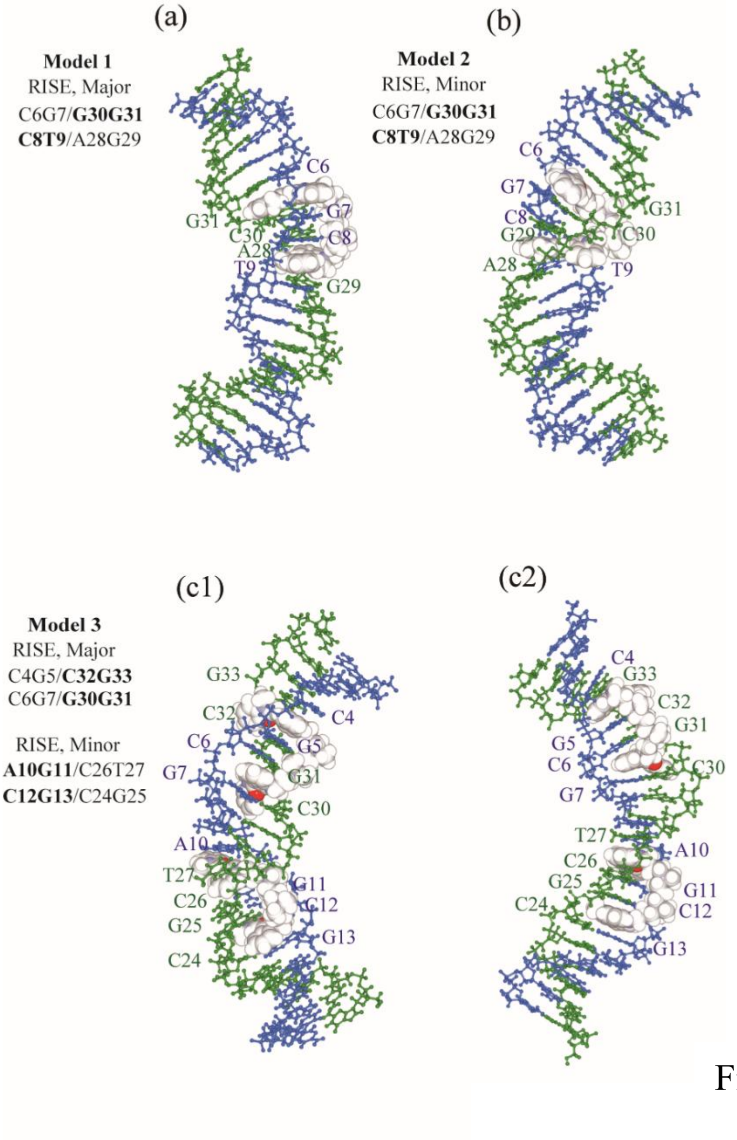

Figure S9C

Figure S9C: bent DNA at $d < 54.0$ Å. (for YOYO)

For YOYO, snapshots for (a) model 1 at $d_{\text{fix}} = 55.0$ Å, (b) model 2 at $d_{\text{fix}} = 55.0$ Å, and (c) model 3 at $d_{\text{fix}} = 60.0$ Å, and (d) model 4 at $d_{\text{fix}} = 63.0$ Å are shown.

For model 1 (RISE-type at the major groove), the DNA bent towards the opposite side of the major groove by 99.9 ± 0.1 % (100, 100, 100, 99.9, and 99.6 % for the window at $d_{\text{fix}}$ = 54.0, 55.0, 56.0, 57.0, and 58.0 Å, respectively). For model 2 (RISE-type at the minor groove), the DNA bent towards the opposite side of the minor groove by 10.7 ± 13.1 % (1.3, 36.5, 6.0, 7.5, and 2.0 % for the window at $d_{\text{fix}}$ = 54.0, 55.0, 56.0, 57.0, and 58.0 Å, respectively) (i.e., the DNA bent towards the same side of the minor groove by 89.3 ± 13.1 %). For model 3 (RISE-types at the major and minor grooves), the DNA bent towards the opposite side of the major groove (for one YOYO at the major groove) by 83.3 ± 11.7 % (60.8, 90.5, 86.8, 84.1, 94.1 % for the window at dfix $d_{fix}$ = 60.0, 61.0, 62.0, 63.0, and 64.0 Å, respectively), and the DNA bent towards the opposite side of the minor groove (for the other YOYO at the minor groove) by 67.4 ± 20.4 % (41.2, 99.5, 50.5, 69.9, and 76.0 % for the window at dfix $d_{fix}$ = 60.0, 61.0, 62.0, 63.0, and 64.0 Å, respectively).

Consequently, RISE-type DNA with one YOYO molecule at the major and minor groove showed that the DNA bend opposite side of the intercalator by 99.9 ± 0.1 % and 10.7 ± 13.1 %, respectively, while RISE-type DNA with two YOYO molecules at the major and minor groove showed the DNA bend opposite side of the intercalator by 83.3 ± 11.7 % and 67.4 ± 20.4 % for model 4. This indicates that the DNA tends to bend towards the opposite side for YOYO at the major groove, and towards the same side for YOYO at the minor groove.

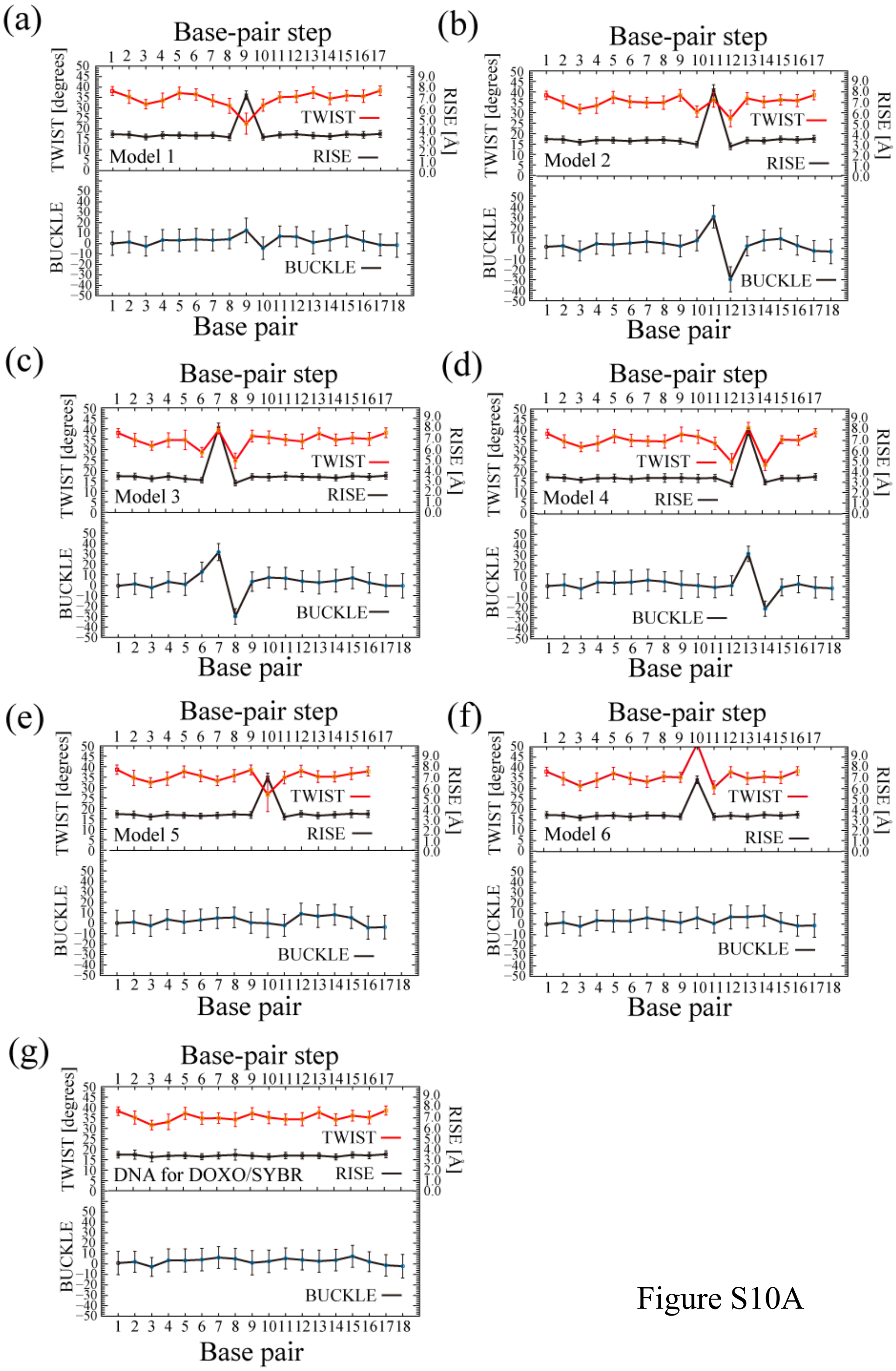
(a)
Base-pair step
TWIST [degrees]
RISE [Å]
TWIST
RISE
Model 1
BUCKLE
Base pair
(b)
Base-pair step
TWIST [degrees]
RISE [Å]
TWIST
RISE
Model 2
BUCKLE
Base pair
(c)
Base-pair step
TWIST [degrees]
RISE [Å]
TWIST
RISE
Model 3
BUCKLE
Base pair
(d)
Base-pair step
TWIST [degrees]
RISE [Å]
TWIST
RISE
Model 4
BUCKLE
Base pair
(e)
Base-pair step
TWIST [degrees]
RISE [Å]
TWIST
RISE
Model 5
BUCKLE
Base pair
(f)
Base-pair step
TWIST [degrees]
RISE [Å]
TWIST
RISE
Model 6
BUCKLE
Base pair
(g)
Base-pair step
TWIST [degrees]
RISE [Å]
TWIST
RISE
DNA for DOXO/SYBR
BUCKLE
Base pair

Figure S10A

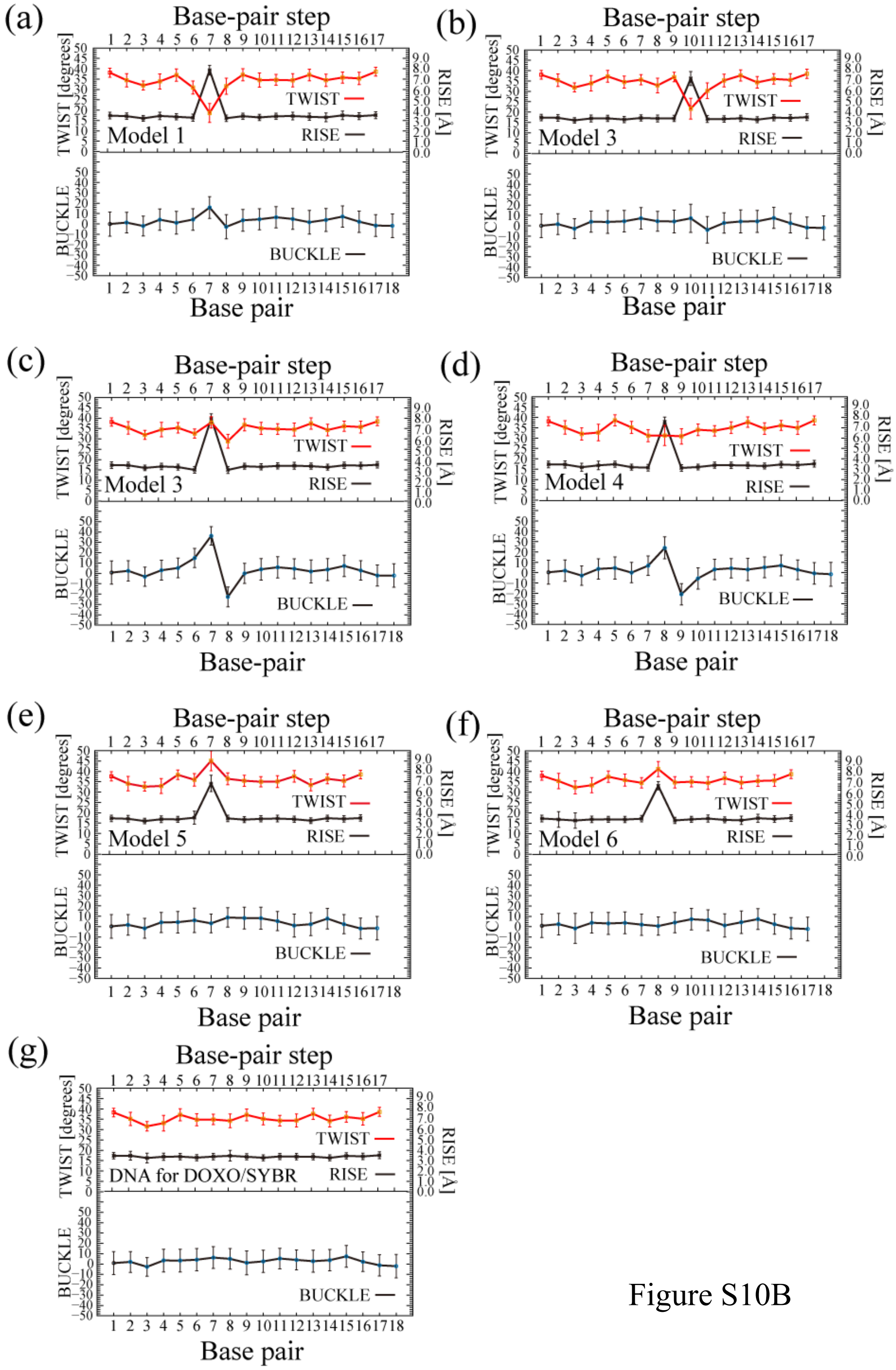
(a)
Base-pair step
TWIST [degrees]
RISE [Å]
TWIST
RISE
Model 1
BUCKLE
Base pair
(b)
Base-pair step
TWIST [degrees]
RISE [Å]
TWIST
RISE
Model 3
BUCKLE
Base pair
(c)
Base-pair step
TWIST [degrees]
RISE [Å]
TWIST
RISE
Model 3
BUCKLE
Base-pair
(d)
Base-pair step
TWIST [degrees]
RISE [Å]
TWIST
RISE
Model 4
BUCKLE
Base pair
(e)
Base-pair step
TWIST [degrees]
RISE [Å]
TWIST
RISE
Model 5
BUCKLE
Base pair
(f)
Base-pair step
TWIST [degrees]
RISE [Å]
TWIST
RISE
Model 6
BUCKLE
Base pair
(g)
Base-pair step
TWIST [degrees]
RISE [Å]
TWIST
RISE
DNA for DOXO/SYBR
BUCKLE
Base pair

Figure S10B

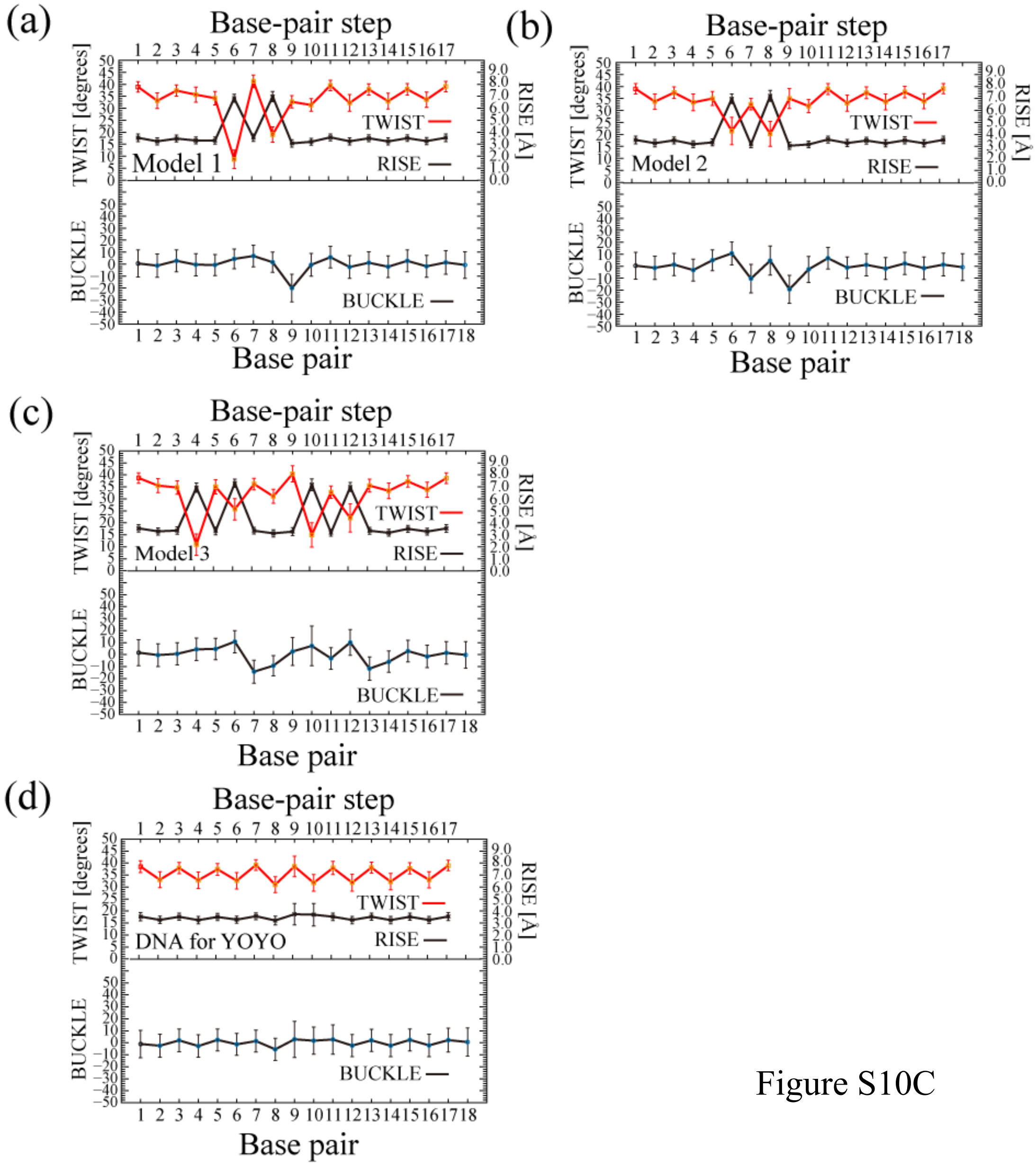

Figure S10C

Figure S10: *Rise* and *Twist* along the base-pair step, and *Buckle* along the base pair

(A) DOXO for models 1–6 (a–f). For comparison, the values for the DOXO/SYBR sequence DNA are shown in (g)

(B) SYBR for models 1–6 (a–f). For comparison, the values for the DOXO/SYRB equence DNA are shown in (g).

(C) YOYO for models 1–3 (a–c). For comparison, the values for the YOYO sequence DNA are shown in (d).

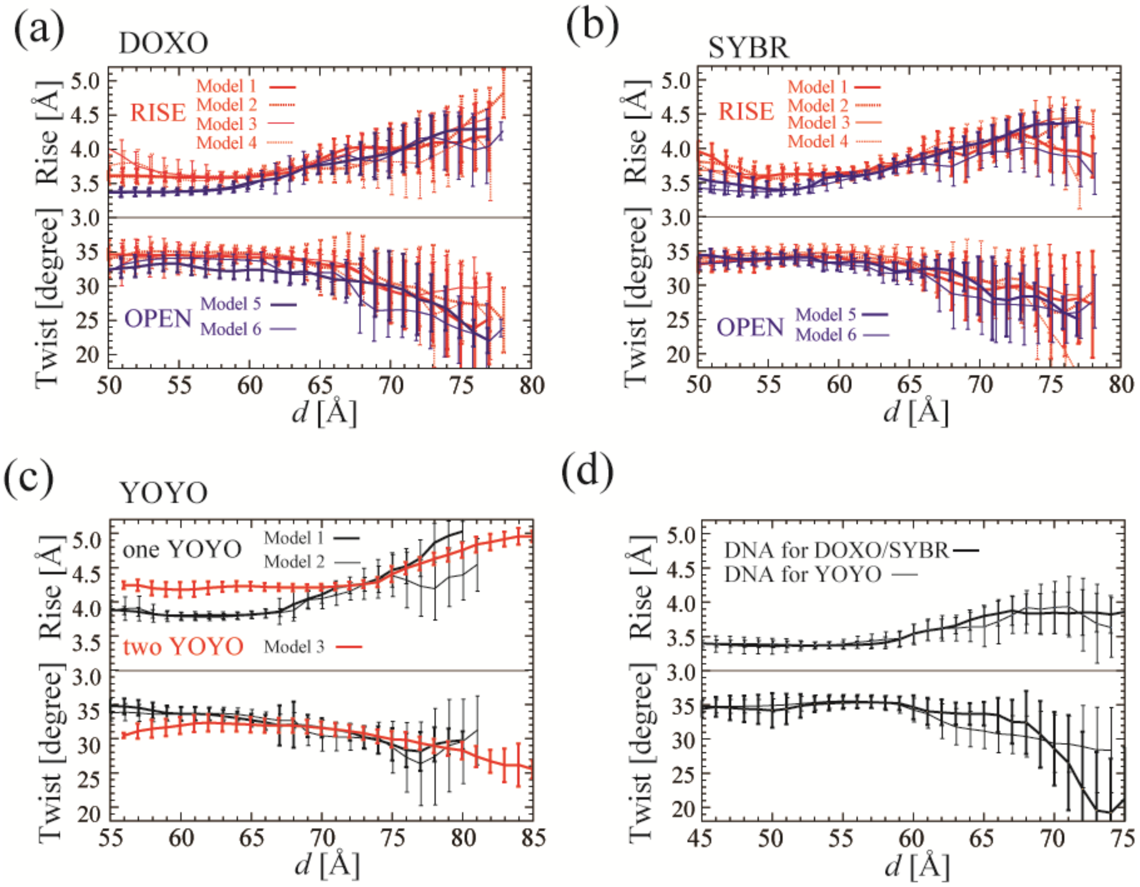

Figure S11: *Rise* and *Twist* of the intercalated DNA along *d*.

*Rise* and *Twist* of the intercalated DNA along *d* for (a) DOXO, (b) SYBR, (c) YOYO, and (d) the free dsDNA, respectively.

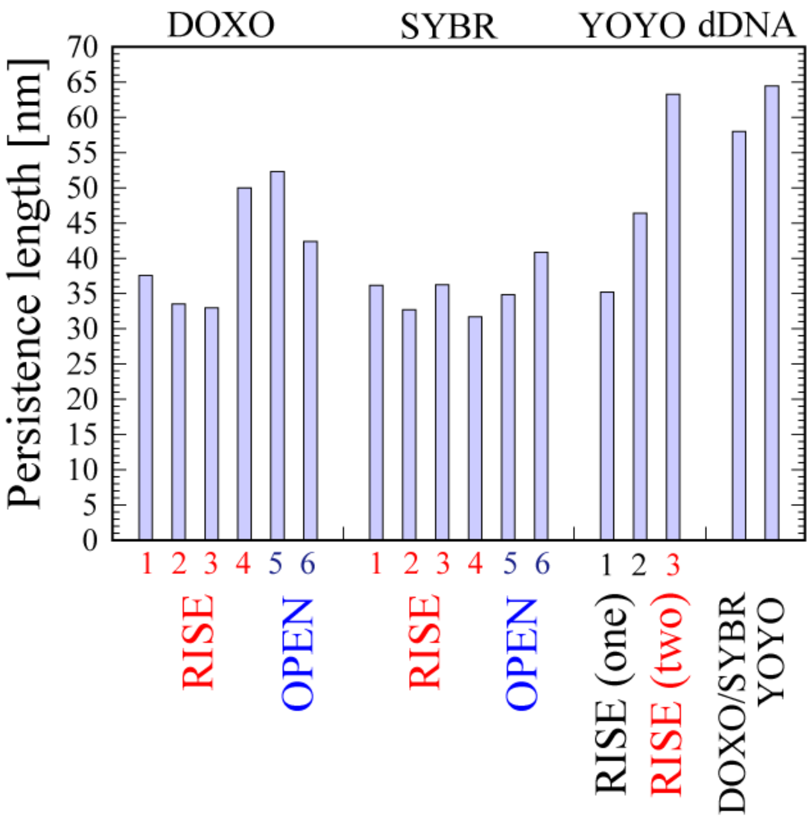

Figure S12: The persistence lengths of the intercalated DNA

The persistence lengths of the intercalated DNA for DOXO, SYBR, and YOYO, and the free dsDNA are shown.

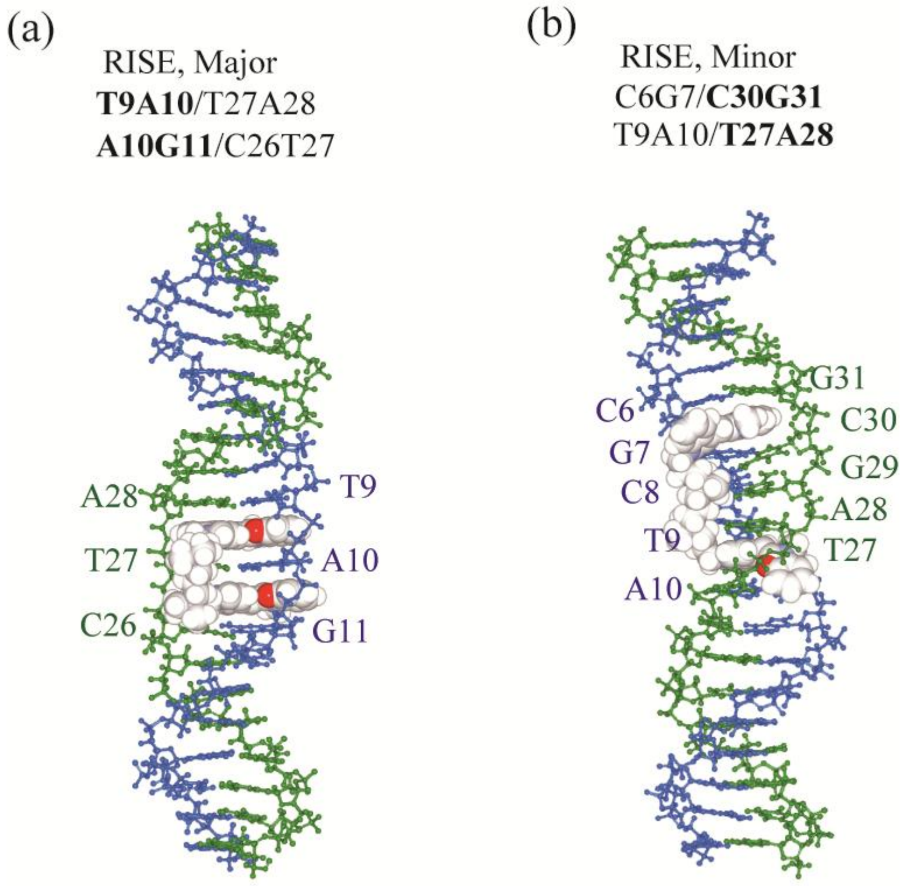

Figure S13: Bis-intercalation of YOYO with (a) one base pair and (b) three base pairs between the two YO-moieties of YOYO

**Table SI. The summary of the procedure of the ABMD and umbrella sampling simulations for intercalator -DNA complex**

| Simulation | Number of particles | Number of replicas for (1) and (2), and number of windows for (3) and (4) | Simulation time for each replica and window [ns] | Simulation time [ns] |
|---|---|---|---|---|
| (1) ABMD and conventional MD to generate the intercalated DNA (Fig. 2) | ~113,000 | 24 for each $\tau$ (1000, 2500, 5000 ps) and the conventional MD) | 700 | 67,200 (DOXO)<br>67,200 (SYBR)<br>67,200 (YOYO) |
| (2) ABMD to estimate the biasing potential for the intercalated DNA | ~113,000 | 24 × 6 models (DOXO)<br>24 × 6 models (SYBR)<br>24 × 3 models (YOYO) | 30 | 4,320 (DOXO)<br>4,320 (SYBR)<br>2,160 (YOYO) |
| (3) Umbrella sampling simulations to refine the biasing potential | ~113,000 | Same as above | 20–40 | ~ 4,320 (DOXO)<br>~ 4,320 (SYBR)<br>~ 2,160 (YOYO) |
| (4) Umbrella sampling simulation for production run | ~113,000 | Same as above | 500 | 72,000 (DOXO)<br>72,000 (SYBR)<br>36,000 (YOYO) |
| Total | | | | 147,840 (DOXO)<br>147,840 (SYBR)<br>107,520 (YOYO) |

Table SII. The summary of intercalation rate

| Intercalator, Base-pair step and intercalation types | Expected fraction of intercalation at random | Fraction of intercalation in ABMD | Fraction of intercalation in conventional MD |
|---|---|---|---|
| **DOXO** | | | |
| AA/TT | 3/17=17.6% | 5.2% | 0.3% |
| AC/GT | 3/17=17.6% | 32.6% | 99.7% |
| CA/TG and TG/CA | 1/17=5.9% and 1/17=5.9% | 0.9% and 4.7% | ND and ND |
| GA/TC | 3/17=17.6% | 32.0% | ND |
| AT/AT | 1/17=5.9% | 7.5% | ND |
| CG/CG | 3/17=17.6% | 8.3% | ND |
| GC/GC | 2/17=11.8% | 8.7% | ND |
| A:T and C:G | 8/18=44.1% and 10/18=55.6% | 47.8* % and 52.2% (RISE-type)76.0% and 24.0% (OPEN-type) | 50.1% and 49.9% ND (OPEN-type) |
| RISE-type and OPEN-type | | $1.19\times10^{-1}$ and $1.31\times10^{-2}$ (90.0% and 10.0%) | $1.89\times10^{-2}$ and 0 (100% and 0%) |
| Major and Minor (RISE-type) | | $5.37\times10^{-2}$ and $6.48\times10^{-2}$ (45.3% and 54.7% ) | $1.89\times10^{-2}$ and 0 (100% and 0%) |
| Major and Minor (OPEN-type) | | ($1.26\times10^{-2}$ and $5.36\times10^{-4}$ (95.9% and 4.1%) | ND |
| **SYBR** | | | |
| AA/TT | 3/17 = 17.6% | 28.7% | 0.2% |
| AC/GT | 3/17 = 17.6% | 15.6% | ND |
| CA/TG and TG/CA | 1/17 = 5.9% and 1/17 = 5.9% | 10.2% and 16.7% | 27.4% and 39.1% |
| GA/TC | 3/17 = 17.6% | 3.5% | 33.2% |
| AT/AT | 1/17 = 5.9% | 1.1% | ND |

| | | | |
|---|---|---|---|
| CG/CG | 3/17 = 17.6% | 13.0% | ND |
| GC/GC | 2/17 = 11.8% | 11.2% | ND |
| A:T and C:G | 8/18=44.1% and 10/18=55.6% | 60.7% and 39.3% (RISE-type)<br>83.1% and 16.9% (OPEN-type) | 50.1% and 49.9% (RISE-type)<br>ND (OPEN-type) |
| RISE-type and OPEN-type | | $1.44\times10^{-1}$ and $2.90\times10^{-2}$ (83.3% and 16.7%) | $2.59\times10^{-2}$ and 0 (100% and 0%) |
| Major and Minor (RISE-type) | | $3.23\times10^{-2}$ and $1.12\times10^{-1}$ (22.4% and 77.3%) | $7.09\times10^{-3}$ and $1.88\times10^{-2}$ (27.4% and 72.6%) |
| Major and Minor (OPEN-type) | | $1.98\times10^{-2}$ and $9.17\times10^{-3}$ (68.3% and 31.7%) | ND |
| **YOYO** | | | |
| AG/CT and CT/AG | 1/17=5.9% and 1/17=5.9% | 9.3% and 8.8% | ND and 0.6% |
| TA/TA | 1/17=5.9% | 8.5% | ND |
| CG/CG | 6/17=35.3% | 67.9% | 99.4% |
| GC/GC | 8/17=47.1% | 5.4% | ND |
| A:T and C:G | 2/18 = 11.1% and 16/18 = 88.9% | 17.6% and 82.4% (RISE-type)<br>ND (OPEN-type) | 0.3% and 99.7% (RISE-type)<br>ND (OPEN-type) |
| RISE-type and OPEN-type | | $3.47\times10^{-1}$ and 0 (100% and 0%) | $1.22\times10^{-1}$ and 0 (100% and 0%) |
| Major and Minor (RISE-type) | | $1.53\times10^{-1}$ and $1.94\times10^{-1}$ (44.1% and 55.9%) | $4.58\times10^{-2}$ and $7.67\times10^{-2}$ (37.4% and 62.6%) |
| Major and Minor (OPEN-type) | | ND | ND |

47.8% was calculated from 5.2 % from AA/TT, 32.6 / 2 % from AC/GC, (0.9 + 4.7) / 2 % from CA/TG and TG/CA, 32.0 / 2 % from GA/TC, 7.5 % from AT/AT.

**Table SIII. The summary of kinetics of intercalation**

| Intercalator | Method | τ [ps] | Fiting parameter [s] First and second intercalation | Estimated value [$s^{-1}$ $M^{-1}$] | Experimental value [$s^{-1}$ $M^{-1}$] |
|---|---|---|---|---|---|
| DOXO | ABMD | 1,500 | $2.71\times10^{-6}$ | $1.24\times10^{8}$ | $7.0\times10^{6}$ at 0.15 M NaCl [34] |
| | | 2,500 | $3.94\times10^{-6}$ | $8.55\times10^{7}$ | |
| | | 5,000 | $3.25\times10^{-6}$ | $1.04\times10^{8}$ | |
| | Conventional | | ND | ND | |
| SYBR | ABMD | 1,500 | $2.04\times10^{-6}$ | $1.65\times10^{8}$ | $1.2 \pm 0.6\times10^{7}$ at 0.1 M NaCl [9] |
| | | 2,500 | $2.38\times10^{-6}$ | $1.41\times10^{8}$ | $2.1 \pm 0.8\times10^{6}$ at 1.0 M NaCl [9] |
| | | 5,000 | $2.85\times10^{-6}$ | $1.18\times10^{8}$ | |
| | Conventional | | ND (no data) | ND | |
| YOYO | ABMD | 1,500 | $5.08\times10^{-7}$ and $4.34\times10^{-6}$ | $6.63\times10^{8}$ and $7.76\times10^{7}$ | YO $3.2 \pm 1.7\times10^{7}$ at 0.1 M NaCl [9] |
| | | 2,500 | $4.62\times10^{-7}$ and $8.07\times10^{-6}$ | $7.29\times10^{8}$ and $4.17\times10^{7}$ | YOYO $3.3 \pm 2.8\times10^{5}$ at 1.0 M NaCl [9] |
| | | 5,000 | $5.58\times10^{-7}$ and $5.39\times10^{-6}$ | $6.03\times10^{8}$ and $6.25\times10^{7}$ | YOYO $3.8 \pm 0.7\times10^{5}$ [33] |
| | Conventional | | $1.62\times10^{-6}$ and ND | $2.08\times10^{8}$ and ND | |

The concentration of the intercalator for the kinetic analysis was assumed to be 5940 μM / 2 = $2.970\times10^{-3}$ M as only two intercalator molecules out of four were involved in intercalation. $k_{on}$ for the second intercalation of the second YO-moiety (bis-intercalation of YOYO) was measured from the time for the first intercalation of the first YO-moiety.

**Table SIV. The summary of physical properties of the intercalated DNA**

| Intercalator and Model | Intercalation site | Persistence length [nm] (See Fig. 11 and Fig. S11) | Stretch modules estimated from contraction and extension [pN] (See Fig. 7 and Fig. S2), and force at the first peak for contraction and extension [pN] (See Fig.9 and Fig. S5) | Δ*Twist* [degree] (See Fig. S9) | $d_{min}$ and elastic range [Å] and correspond values for the contour length |
|---|---|---|---|---|---|
| **DOXO** | | | | | |
| **Model 1** | **G9A10**/T27C28 | 37.58 | 1676 and 2706 (2668) | –14.7 (=) | 60.2 and 58.2–62.3 |
| RISE, Major | (A11:T26) | | –40.7 (58.3) and 94.4 (62.7) | | (64.3 and 63.9–65.5) |
| **Model 2** | **A11C12**/G25T26 | 33.51 | 1730 and 3916 (4077) | –4.7, 1.9, and –7.0 (+) | 60.7 and 58.1–61.8 |
| RISE, Major | (G13:C24) | | –59.3 (58.2) and 79.9 (62.5) | | (64.1, 63.9–64.7) |
| **Model 3** | A7C8/**G29T30** | 32.99 | 2029 and 3938 (2283) | –6.1, 4.4, and –9.6 (+) | 60.7, 58.4 and 61.7 |
| RISE, Minor | (G9:C28) | | –60.5 (58.6) and 51.9 (62.1) | | (63.8 amd 63.8–65.4) |
| **Model 4** | **G13A14**/T23C24 | 49.96 | 2295 and 2533 (1842) | –9.7, 3.4, and –10.9 (+) | 60.3 and 58.7–62.8 |
| RISE, Minor | (C12:G25) | | –43.9 (58.8) and 86.8 (62.6) | | (64.3 and 64.3–65.3) |
| **Model 5** | **A10C12**/G25T27 | 52.32 | 1703 and 1826 (692) | –42.8 (=) | 57.2 and 55.6–59.2 |
| OPEN, Major | (G13:C24) | | –36.6 (55.3) and 47.3 (59.1) | | 61.9 and 60.9–63.7 |
| **Model 6** | A10C12/**G25T27** | 42.40 | 1566 and 2506 (1620) | –18.0 (=) | 57.1 and 54.9–59.7 |
| OPEN, Minor | (G13:C24) | | –60.6 (53.0) and 93.0 (59.4) | | (60.6 and 59.9–62.8) |
| **SYBR** | | | | | |
| **Model 1** | A7C8/**G29T30** | 36.14 | 1280 and 1922 (1862) | –16.4 (=) | 60.3 and 58.6–63.5 |
| RISE, Major | | | –31.6 (58.1) and 84.2 (63.1) | | (65.4 and 64.5–66.3) |
| **Model 2** | **A10A11**/T26T27 | 32.72 | 1297 and 2095 (1283) | –13.5 (=) | 60.1 and 58.3–63.0 |
| RISE, Major | | | –32.6 (57.9) and 81.3 (62.7) | | (64.5 and 64.2–67.9) |

| | | | | | |
|---|---|---|---|---|---|
| **Model 3** | A7C8/**G29T30** | 36.25 | 2469 and 2803 (2283) | –2.2, 3.13, and –5.4 (+) | 60.3 and 58.8–62.6 |
| RISE, Minor | | | –46.5 (58.6) and 87.0 (62.5) | | (64.5 and 64.1–67.2) |
| **Model 4** | C8G9/**C28G29** | 31.70 | 1497 and 2430 (1898) | –3.7, –2.9, and –5.8 (+) | 60.0 and 58.1–62.0 |
| RISE, Minor | | | –36.8 (57.6) and 61.1 (61.8) | | (64.3 and 64.2–66.5) |
| **Model 5** | **A7G9**/C28T30 | 34.86 | 2146 and 1376 (1928) | –24.0 (=) | 56.4 and 55.0–58.4 |
| OPEN, Minor | | | –51.8 (53.5) and 25.5 (57.8) | | (60.5 and 60.4–63.2) |
| **Model 6** | C8A10/**T27G29** | 40.84 | 1892 and 1522 (1127) | –30.2 (=) | 55.9 and 54.5–58.2 |
| OPEN, Minor | | | –51.5 (53.6) and 47.1 (57.8) | | (60.2 and 60.1–64.2) |
| **YOYO** | | | | | |
| **Model 1** | C6G7/**C30G31** | 35.21 | 1308 and 2103 (1980) | –23.9 (=) | 63.0 and 61.3–66.3 |
| RISE, Major | **C8T9**/A28G29 | | –28.3 (60.5) and 91.9 (66.4) | –12.0 (+) | (67.6 and 67.6–70.8) |
| | | | | 2.0 at G7C8/C30G29 | |
| **Model 2** | C6G7/**C30G31** | 46.41 | 1465 and 2810 (2639) | –11.2 (=) | 63.2 and 61.3–66.4 |
| RISE, Minor | **C8T9**/A28G29 | | –40.4 (60.3) and 136 (66.7) | –10.8 (+) | (67.8 and 67.1–68.8) |
| | | | | –6.6 at G7C8/C30G29 | |
| **Model 3** | C4G5/**C32G33** | 63.24 | 2880 and 2710 (2228) | –21.9 (=) | 70.1 and 68.8–73.3 |
| RISE, Major | C6G7/**C30G31** | | –36.4 (68.7) and 117 (73.6) | –7.0 (+) | (74.8 and 74.3–76.7) |
| | | | | –2.4 at G5C6/C32G31 | |
| | **A10G11**/C26T27 | | | –16.8 (=) | |
| RISE, Minor | **C12G13**/C24G25 | | | –9.9 (+) | |
| | | | | –5.2 at G11C12/C26G25 | |
| DNA for | | 57.98 | 2315 and 3277 (2238) | | 56.8 and 55.0–58.8 |
| SYBR/DOXO | | | –67.3 (54.2) and 94.4 (58.5) | | (60.3 and 60.3–62.4) |
| DNA for | | 64.42 | 1991 and 3449 (2351) | | 56.7 and 54.7–58.2 |
| YOYO | | | –53.8 (54.6) and 92.9 (58.7) | | (60.4 and 60.2–62.3) |

The elastic regions for extension and contraction were defined so that the quadratic fitting from $d = d_{\min}$ to the largest and smallest $d$ where the RMSD of the free energies < than 0.02 kcal/mol.

(+) and (=) in the column of Δ*Twist* represent perpendicular and parallel orientations of the intercalator, respectively. For DOXO and SYBR, Δ*Twist* (+) are at the ($i$-1)th, $i$-th, and ($i$+1)-th base-pair steps, and Δ*Twist* (=) are at the $i$-th base-pair step (intercalation site).

The values in parentheses at the stretch modules represent the estimated stretch modulus derived from the contour length (see Eq. (6)).

The values in parentheses at force at the first peak are the average force after the first peak.

The values in parentheses at $d_{\min}$ are the corresponding values of the contour length.

The values in parentheses at elastic range the corresponding ranges of the contour length.

For DOXO, the base-pair step in bold (**G9A10** in model 1**)** indicates that the O4 atom of DOXO is oriented towards it. The base pair in parenthesis (A11:T26 in model 1) indicates that the amino group of DOXO is oriented towards it.

For SYBR, the base-pair in bold (**T30G29** in model 1) indicates that the oxygen atom attached to the quinoline group in SYBR is oriented towards it.

For YOYO, the base-pair in bond (**G31C30** and **C8T9** in model1) indicates that the oxygen atom of the benzoxazole group in YOYO is oriented towards it.
